# Placental invasion mismatch underlines pregnancy disorders and cancers

**DOI:** 10.64898/2026.07.06.736876

**Authors:** Xin Li, Ruixue Chen, Yangyi Zhang, Yuchen Sun, Wenqiang Du, Dan Yu, Yanlin Lu, Yiqing Yang, Xupeng Bi, Yijun Yang, Jintian Zhu, Kailin Sun, Jing Liang, Lin Jiang, Yunqiu He, Liqun Sun, Junhua Shen, Kshitiz, Dan Zhang, Guojie Zhang

**Affiliations:** Women’s Hospital, Zhejiang University School of Medicine & Center for Evolutionary & Organismal Biology, Liangzhu Laboratory, Zhejiang University, Hangzhou, China; Department of Biomedical Engineering, University of Connecticut Health, Farmington, CT 06030; Villum Center for Biodiversity Genomics, Department of Biology, University of Copenhagen, Copenhagen 2100, Denmark

## Abstract

Mammalian placentas vary dramatically in invasiveness, parallel aggressive cancers, and are dysregulated in pregnancy disorders, yet whether they share regulatory architecture remains unclear. We investigated single-cell transcriptomes of the maternal-fetal interface across nine mammals spanning all major placental morphotypes and integrated it with thirteen cancers and five pregnancy complications. A conserved cellular framework is deployed through three discrete regulatory programs: a cancer-like program in hemochorials, endothelial-cooperation program in endotheliochorials, and collagen-rich invasion-suppressing program in epitheliochorials. Aggressive cancers selectively converge on hemochorial program, and we functionally validated share invasion regulators including the VGLL3-TEAD1 interaction and *APOE*. Pregnancy disorders are partially, mismatched deployments of these programs; placental *APOE* knockdown in mice phenocopies preeclampsia with concurrent collapse of both M1/M2 macrophage programs. These findings unify placental diversity, cancer convergence, and obstetric disorders under a regulatory-mismatch principle, whereby evolved placental invasion programs becomes pathological when deployed outside their evolutionary context.

## Introduction

The evolution of viviparity and placentation represents a profound innovation that transformed mammalian reproductive strategies and empowered their diversification^1–3^. At the heart of this innovation lies the biological paradox that fetal cell-lineages must invade maternal tissues for nutrient acquisition while this invasion must be finetuned to prevent tissue damage and immune rejection^4–6^. Different mammalian lineages have solved this challenge in dramatically diverse ways, producing wide diversity in placental invasion depth^7^. Hemochorial placentas of primates and rodents support deep trophoblast invasion into maternal blood, whereas epitheliochorial placentas of ungulates leave maternal tissue layers intact, and endotheliochorial placentas of carnivores fall between these extremes^8^. Phylogenetic analyses indicate that the ancestral eutherian placenta was likely deeply invasive, with reduced invasion evolving repeatedly along several lineages under lineage-specific selection^9^. Evolutionary medicine has framed this divergence as the joint outcome of parent-offspring conflict^4,10,11^ and reciprocal coevolution between fetal invasion and maternal stromal resistance^12,13^. These evolutionary considerations also bear on human medicine, since trophoblast invasion is pathologically altered in major obstetric disorders^14^ and the invasive properties of aggressive cancers have long been compared to those of trophoblasts^15,16^. Preeclampsia, fetal growth restriction, and recurrent miscarriage involve inadequate trophoblast invasion, whereas placenta accreta spectrum involves excessive invasion^17–20^. Their clinical phenotypes resemble natural states across mammalian placentation. At the molecular level, human preeclampsia-risk loci overlap with genes under selection in lineages of convergent placental-invasion reduction^21^, framing obstetric disease as a developmental recapitulation of phylogenetic shifts. Invasive trophoblasts and malignant tumors have long been recognized to share epithelial-mesenchymal transition, matrix degradation, immune evasion, and angiogenesis^15,22^, suggesting that aggressive cancers in mammals redeploy programs evolved for controlled invasion. Together, these observations raise a question that has remained unanswered at cellular and molecular resolution. What features distinguish the ancestral placental configurations that support viable pregnancy in mammalian lineages from their pathological reactivation in human disease, and does the same regulatory-mismatch principle account for the convergence between aggressive cancer invasion and the most invasive placental regime?

Single-cell transcriptomics has begun to converge on a coherent picture of the maternal-fetal interface on which three conclusions are broadly accepted. The cellular machinery for trophoblast invasion is deeply conserved across mammals^23–25^, even as the transcription-factor networks regulating it vary substantially across placental morphologies^26^, implying that placental diversity reflects differential regulation of shared cellular components rather than gain or loss of invasive cell types. This regulation has been shaped reciprocally by maternal and fetal evolution, with maternal endometrial gene expression scaling with depth of placental invasion, stromal cells varying across lineages in their resistance to invasion, and fetal-maternal signaling exhibiting coopetition rather than pure antagonism^2,13,27^. The long-noted parallels between trophoblast and tumor invasion are tied to this same evolutionary architecture, with cancer malignancy rates across mammals scaling with placental invasiveness and cancer transcriptomes identifying shared oncofetal regulatory programs^12,28^. These conclusions imply three open components of the question above, namely whether placental regulatory diversity falls into qualitatively distinct molecular regimes or varies continuously, whether the cancer-placenta correspondence converges specifically with the most invasive configuration or extends generically across morphotypes, and what distinguishes viable deployment of ancestral configurations from their pathological reactivation in human disease. Addressing these requires comparative placental single-cell data at the invasion-active developmental window across all morphotypes, integrated with human pregnancy-disorder data and multi-cancer transcriptomes, an integration has not previously been assembled.

Here we address these questions through an integrated comparative single-cell analysis of the maternal-fetal interface. We first establish the cellular composition and evolutionary divergence of cell types across the nine species, identifying which cellular compartments are most variable across the spectrum of placental invasion depth. We next characterize the molecular regulatory programs of trophoblasts across morphotypes, testing whether placental invasion diversity is organized as discrete molecular regimes or as continuous tuning of a shared program. The regulatory network underlying the cancer-like invasive program in hemochorial placentas was functionally validated in three-dimensional stromal invasion assays. We then analyze the intercellular signaling architectures at each maternal-fetal interface, comparing the empirical patterns with predictions from parent-offspring conflict and reciprocal coevolution. We further apply this comparative framework to human pregnancy disorders, identifying the molecular and cellular configurations into which preeclampsia and recurrent miscarriage revert, comparing these with the natural states of mammalian lineages in which similar configurations support viable pregnancy. Finally, we integrate the placental data with single-cell transcriptomes from thirteen human cancer types to ask whether cancer-placenta convergence is generic across mammalian placentation or specific to particular regulatory configurations, and functionally validate predicted regulatory nodes by *in vivo* knockdown experiments. Together, these analyses provide a unified comparative framework for understanding how the cellular logic of placental evolution informs both human obstetric pathology and the long-noted parallels between placental invasion and aggressive cancer.

## Results

### Coordinated convergence of cell-type expression at the maternal-fetal interface by placental morphotype

To elucidate the molecular logic linking conserved cellular machinery to morphological diversification of placental invasion, we generated single-cell transcriptomic atlases of maternal-fetal interfaces during active trophoblast invasion across nine mammalian species (Fig. 1a, and Extended Data Fig. 1a), This atlas extends two recent cross-species single-cell resources, the six-species mid-gestation atlas of Stadtmauer et al. (2025)^25^ and the ten-species late-gestation atlas of Tan et al. (2026)^26^, in three respects. We sample the early- and mid-gestation window during which trophoblast invasion is actively progressing across all three eutherian morphotypes. We also add Chinese tree shrew as an endotheliochorial reference and one of the convergently derived lineages of reduced placental invasiveness^7^; and sugar glider as a marsupial outgroup. In total, the atlas comprises approximately 906,855 high-quality cells with a mean of 100,762 cells per species, spanning hemochorial (humans, cynomolgus macaque, and mouse), endotheliochorial (tree shrew, dog), epitheliochorial (sheep, goat, pig), and marsupial sugar gilder placentation (Extended Data Fig.1b, and Supplementary Table 1).

**Fig. 1.**
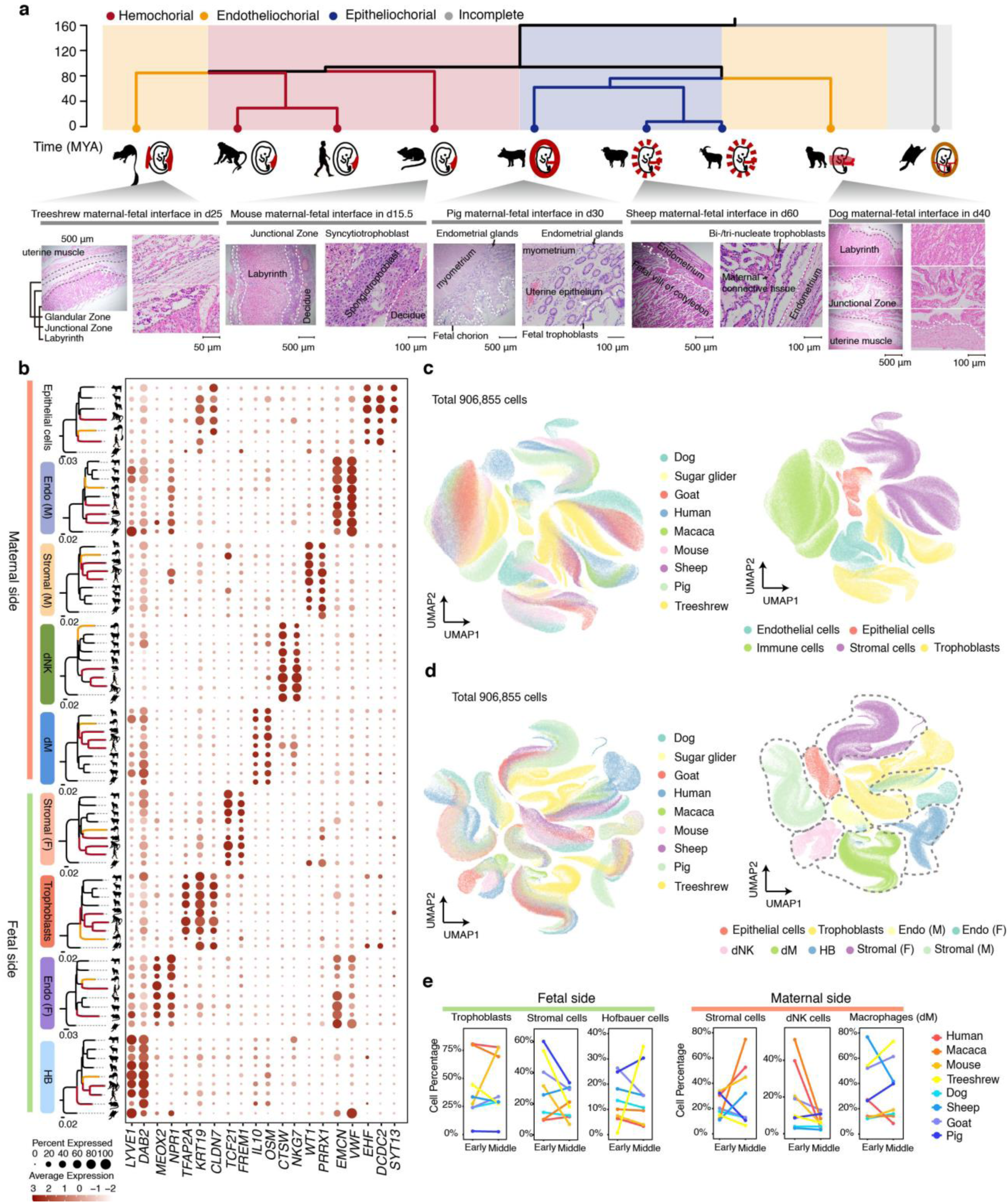
Single-cell transcriptomic profiling of nine mammalian species representing diverse maternal-fetal interfaces. **a,** The evolution and structure of maternal-fetal interfaces. Top: a phylogenetic tree of the mammalian species sampled. Bottom: Histology of the maternal-fetal interface of house mouse (*Mus musculus*), Chinese tree shrew (*Tupaia belangeri chinensis*), Beagle dog (*Canis lupus familiaris*), Hu sheep (*Ovis aries*) and Suziblack pig (*Sus scrofa domestica*) from the mid-gestation. **b,** Gene expression trees based on pseudo-bulk transcriptomes and marker genes expression for each cell type. **c,d,** UMAP visualization of integrated maternal–fetal interface cells from nine species generated using SATURN based on major cell lineages (c) and refined cell types (d). Colors indicate different species (left) and cell types (right). **e,** Line charts showing the proportion of major fetal and maternal cell types in the early-gestation and mid-gestation across eight species.

Unbiased clustering and marker-gene annotation identified nine major cell types present across all species: trophoblasts, maternal and fetal stromal cells, maternal and fetal endothelial cells, maternal epithelial cells, and three immune populations, decidual macrophages (dM), decidual natural killer cells (dNK), and fetal macrophages (HB)) (Fig. 1b–d, Extended Data Fig. 2a, b, 3a and Supplementary Table 2). Cross-species integration using SATURN^29^, which aligns single-cell transcriptomes through protein language model embeddings, recovered consistent cell-type correspondences across all species (Fig. 1c, d). These observations confirm, and extend to additional species, the cross-eutherian conservation of materal-fetal inference cell type architecture established by previous studies^25,26^, and indicate that placental morphological diversity does not arise from the presence or absence of distinct interface cell types.

We next compared cellular compartment composition across morphotypes and developmental windows. Trophoblasts and maternal stromal cells expanded coordinately in hemochorial species, reaching substantially higher trophoblast proportions than endotheliochorial and epitheliochorial placentas (Fig. 1e). Notably, the mouse trophoblast compartment expanded rapidly to converge on the proportions seen in humans and macaque, despite mice being phylogenetically more distant from primates than tree shrews. This coordinated trophoblast-stromal expansion is consistent with hemochorial placentation requires both enhanced trophoblast populations and a permissive maternal stromal environment^30,31^. Immune cell dynamics also varied with placental morphotype. In hemochorial placentas, dNK cells were proportionally enriched during early gestation and equalized by mid-gestation (Fig. 1e), whereas in non-hemochorial placentas, decidual macrophages (dM) remained elevated at mid-gestation (Fig. 1e). This pattern suggests that hemochorial species concentrate immune regulation during the active-invasion window while non-hemochorial species maintain a sustained immune presence at the maternal-fetal interface. Conversely, fetal macrophages (HB) showed divergent trajectories, declining modestly from early to mid-gestation in human, macaque and mouse, but increasing substantially in tree shrews. Epithelial, endothelial, and fetal stromal cells proportions did no exhibit consistent morphotype-based patterns (Extended Data Fig. 3b). These temporal cross-species comparisons indicate that the fetal trophoblast, maternal stromal, and innate-immune compartments are each tuned to distinct temporal logics across placental morphotypes.

The conservation of cellular architecture across morphotypes raised an interesting question that whether the molecular identity of each conserved cell type is itself species-phylogenetically conserved, or has it converged by placental morphotype. To distinguish these alternatives, we reconstructed gene expression phylogenies from pseudo-bulk transcriptomes for each cell type (Fig. 1a, b, and Extended Data Fig. 4). For four cell types, trophoblasts, dNK, dM and maternal stromal(dS), the expression phylogeny placed humans, cynomolgus monkeys, and mice (all hemochorial, separated by approximately 87 million years of independent evolution^32^) into a monophyletic clade, with tree shrew positioned outside this clade despite its phylogenetic affinity to primates (Fig. 1a, b, and Extended Data Fig. 4). That four cell lineages of distinct developmental origin converge on the same morphotype-based grouping is consistent with coordinated co-evolution of the cells that physically and molecularly interact during placentation, with placental morphotype acting as a shared evolutionary attractor on the maternal-fetal interface as an integrated module. This molecular coordination parallels the compositional coordination we observed at the cellular level, where trophoblast and maternal stromal expansion track together with invasion depth (Fig. 1e).

### Three distinct regulatory programs underlie placental invasion diversity

We next asked whether the diversification of placental invasion depth reflects quantitative tuning of a shared molecular program or qualitatively distinct molecular programs that couple trophoblast programs to placental morphotype. We focused this cross-morphotype molecular comparison on extravillous trophoblasts (EVT) and its homologs across all species, since EVTs are the fetal cells that physically invade maternal tissue and thereby define placental architecture. Standard trophoblasts subtypes: cytotrophoblast (VCT), syncytiotrophoblast (SCT), and extravillous trophoblast (EVT) were annotated using established markers in humans and macaques (Extended Data Fig. 5a–o, Supplementary Tables 3 and 4), and propagated to less-characterized species through cross-species expression alignment (Supplementary Table 5). We confirm, and extend to additional taxa that, the conserved invasive-trophoblast signature identified by Stadtmauer et al. (2025)^25^. EVT-like cells expressing the canonical EVT marker set (*FN1, ITGA1, SERPINE1, COL4A1* and *TGFBR2*) are present not only in hemochorial species but also in the epitheliochorial species (pig, sheep, goat) that prior cross-species analyses did not sample (Fig. 2a, b). Other trophoblast subtypes (VCT, SCT) likewise showed conservation across all species (Extended Data Fig. 5a and 6). The cellular architecture of trophoblast invasion is therefore conserved across eutherian morphotype spectrum, indicating that placental diversity arises from differential regulation of conserved cellular components.

**Fig. 2.**
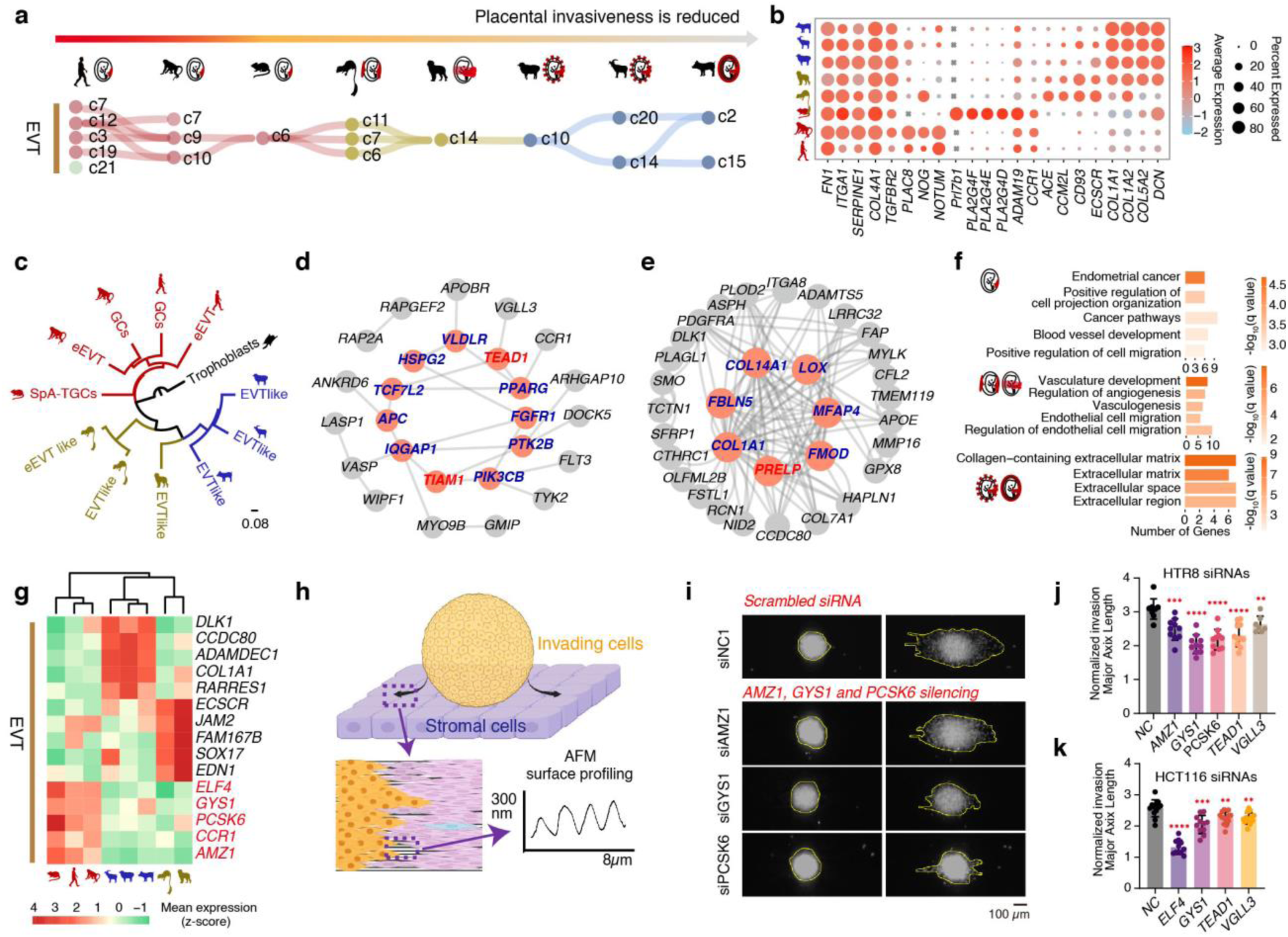
Cross-species comparison of extravillous trophoblast (EVT) and invasion assay. **a,** SAMap mapping relationships between EVTs with an alignment threshold of 0.3. **b,** Marker genes expression of homologue EVTs ascross species. **c,** Expression-based phylogenetic trees of homologous EVTs across eight species, with the trophoblasts of sugar glider (*Petaurus breviceps*) as an outgroup. **d,e,** Up-regulated genes in hemochorial placentas (d) and epitheliochorial placentas (e) with protein-protein interaction network constructions, respectively. Genes marked in red circles are key genes in the interaction network, while genes highlighted in red bold font are hub genes. These were derived based on the combined scores generated by MCODE using STRING. **f,** Pathway enrichment analysis of up-regulated key genes for hemochorial, endotheliochorial and epitheliochorial placentas, respectively. **g,** Heatmap expression of standout candidates DEGs based on combined CPM and Z-score metrics and a stringent cutoff of log_2_FC ≥ 4 for each placenta type. **h,** Schematic for spheroid Accelerated Nanopatterned Stromal Invasion Assay (ANSIA), in which spheroid invasion into surrounding stromal cells, pre-aligned on a nanopatterned substrate to minimize invasion variability arising from stromal collective migration patterns, is monitored by time-lapse imaging. **i,** Fluorescent images showing in-situ *AMZ1, GYS1 and PCSK6* genes silencing on human EVT (HTR8-H2B-tdTomato) spheroid invasion into hESF monolayer (black; unlabeled); Spheroid boundaries are manually outlined. **j,k,** Graph showing normalized major axis length of invading EVT (j) and HCT116 (k) spheroids with scrambled (NC; n = 9 spheroids), or gene silenced for *AMZ1* (*n* = 10 spheroids), *GYS1* (*n* = 10 spheroids), *PCSK6* (*n* = 10 spheroids), *TEAD1* (*n* = 11 spheroids), and *VGLL3* (*n* = 7 spheroids). Statistical significance of differences using unpaired t test between the NC group and the *AMZ1*-, *GYS1*-, *PCSK6*-, *TEAD1*-, and *VGLL3*-silenced groups is indicated by asterisks, where **** denotes *P* < 0.0001, *** denotes *P* < 0.001, and ** denotes *P* < 0.01.

To quantify how EVT expression differs across morphotypes, we constructed expression-based phylogenies from the top 20 EVT-specific marker genes in each species. Strikingly, EVTs from hemochorial species clustered together, while epitheliochorial EVT-like cells formed a separate clade (Fig. 2c). Endotheliochorial species did not cluster together, consistent with these two endotheliochorial species reaching intermediate invasion depth through different gross-anatomic routes, the dogs possess zonary placentas while tree shrews exhibit discoid placentas^33,34^. We next identified genes specifically upregulated in each placenta morphotype through differential expression analysis, revealing 66 hemochorial-enriched genes, 175 endotheliochorial-enriched genes, and 54 epitheliochorial-enriched genes (Supplementary Table 6, and Methods). Protein-protein interaction network analysis identified three distinct hub structure. The hemochorial network is centered on 11 hub genes including *PPARG*, *TEAD1*, and *TIAM1* (Fig. 2d), the endotheliochorial network on 16 hubs including *ENG*, *EDN1* and *ROBO4* (Extended Data Fig.7), and the epitheliochorial network on 7 hubs including *LOX*, *PRELP*, and *COL1A1* (Fig. 2e). The hemochorial program was mostly enriched for cancer-associated pathways (endometrial cancer and hippo-pathway) together with positive regulation of cell migration and blood vessel development, the endotheliochorial program was enriched for vascular and endothelial development, and the epitheliochorial program was enriched for collagen-containing extracellular matrix and ECM-related compartment (Fig. 2f, and Supplementary Table 7). These results define three distinct evolutionary solutions to maternal-fetal exchange: aggressive invasion resembling cancer metastasis in hemochorial species, endothelial cooperation and vascular integration in endotheliochorial species, and matrix stabilization enhancing nutrient exchange through tightly apposition rather than invasion in epitheliochorial species.

To characterize the hemochorial program in functional detail, we focused on *VGLL3* and *TEAD1*, the upstream regulator and central seed-hub gene of the hemochorial network respectively (Fig. 2d). *TEAD1* is a downstream effector of Hippo signaling^35^, one of the cancer-related pathways enriched in the hemochorial program (Fig. 2f, and Supplementary Table 7), making the VGLL3–TEAD1 axis a candidate mechanistic link between the hemochorial cancer-like signature and trophoblast invasion biology. The VGLL3–TEAD1 axis has recently been independently identified by Plazyo et al. (2026)^36^ as dysregulated in human preeclamptic placentas, with placenta-specific *VGLL3* overexpression in mice causing maternal hypertension. Our cross-species data positions *VGLL3-TEAD1* as a constitutive hub of the hemochorial regulatory program (Fig. 2d), so its dysregulation in preeclampsia reflects perturbation of a regulatory module required for hemochorial-specific deep invasion rather than a generic placental machinery. To test the functional requirement of this axis for trophoblast invasion, we used the Accelerated Nanopatterned Stromal Invasion Assay (ANSIA), in which human EVT spheroids invade into a pre-aligned monolayer of human endometrial stromal fibroblasts under time-lapse imaging (Fig. 2h)^12,37^. Knockdown of either *VGLL3* or *TEAD1* significantly reduced EVT spheroids invasion depth (Fig. 2j and Extended Data Fig. 8). Combined with previous reports that *VGLL3* binds *TEAD1* to promote cancer cell proliferation and that *TEAD1* transactive HLA-G expression required for EVT migration^38,39^, supports a conserved *VGLL3*-*TEAD1-*trophoblast invasion pathway that defines the hemochorial regulatory program.

Beyond the network hubs, we identified four hemochorial-enriched genes (*ELF4*, *GYS1*, *PCSK6*, and *AMZ1*) with exceptional expression specifically in hemochorial EVTs (Fig. 2g, Supplementary Table 8 and Methods). The broader hemochorial-enriched gene set also includes regulators with established trophoblast roles, notably *CCR1*^40^ and PCSK6^41^, confirming that the hemochorial program recovers established invasion biology. Using ANSIA, we functionally tested the four newly characterized drivers in human EVT spheroids. Knockdowns of *GYS1*, *PCSK6*, or *AMZ1* significantly reduced spheroid invasion depth (Fig. 2i, j). *ELF4* knockdown completely abolished spheroid formation, precluding direct invasion depth quantification but indicating that this ETS-family transcription factor is required for the trophoblast invasive program at an earlier developmental step. These genes likely support complementary aspects of the invasion process that *GYS1* (glycogen synthase 1) provides metabolic flexibility, energy buffering under hypoxic or nutrient-limited conditions characteristic of deep invasion^42^, *PCSK6* (proprotein convertase) activates extracellular proteases that remodel maternal matrix facilitating tissue invasion^43^, and *AMZ1* (secreted metalloprotease) directly degrades maternal matrix barriers^44^. Together, these findings reveal a hierarchical invasion program in hemochorial placentas, where *ELF4* establishes cellular architecture, *VGLL3*-*TEAD1* initiates the invasive phenotype, and metabolic enzymes (*GYS1*) and protease (*PCSK6*, *AMZ1*) execute matrix penetration. To extend the comparison to cancer cells, we tested the same regulators in the human colon cancer cell line (HCT116) colorectal carcinoma spheroids. Silencing of *ELF4*, *GYS1*, PCSK6, or *TEAD1/ VGLL3* each reduced cancer-cell invasion (Fig. 2k). These functional results extend the cancer-like character of the hemochorial program from pathway-level enrichment to shared functionally validated regulators of invasion in both trophoblasts and cancer cells.

### Deep placental invasion is supported by cooperative rather than antagonistic intercellular signaling

Having identified three distinct regulatory programs operating at the maternal-fetal interface to achieve different invasion depths, we next asked how these programs translate into intercellular communication patterns between fetal and maternal cells at the interface, the battleground where fetal demands for resources meet maternal control over access. Cell-cell communication analysis based on upregulated ligand and receptor genes across cell types revealed structurally divergent signaling architectures across the three morphotypes (Fig. 3a, d, h, and Supplementary Table 9). In hemochorial placentas, EVTs communicated with surrounding populations predominantly through FN1-αVβ5 integrin and PGF-FLT1/NRP1 pathways (Fig. 3b), both promoting cell migration, angiogenesis, and tissue invasion^45–47^. Epitheliochorial EVT-like cells engaged neighboring cells through extensive collagen–integrin and ephrin–Eph interactions (Fig. 3e, f), creating a matrix-anchoring and boundary-reinforcing environment that restricts trophoblast penetration^48,49^. Endotheliochorial placentas exhibited a hybrid pattern combing collagen–integrin signaling with vascular-related interactions (Fig. 3i), including ESAM-mediatedr endothelial junction integrity^50^ and EDN1/VEGFA/VWF-mediated vascular tone^51^, angiogenesis^52^, and endothelial adhesion^53^.

**Fig. 3.**
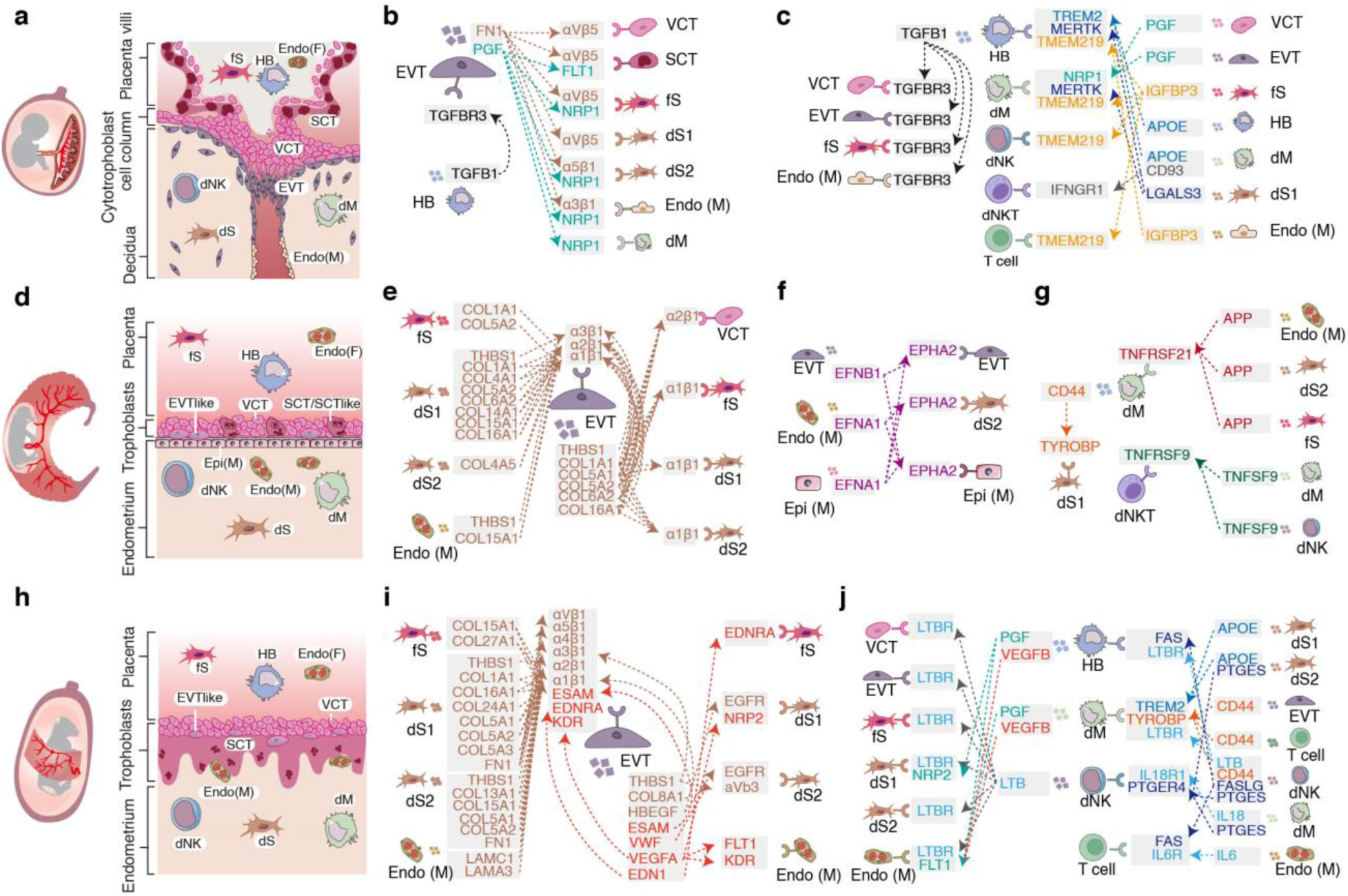
Ligand–receptor signaling across distinct maternal-fetal interfaces. **a,d,h,** Schematic representation of placental morphology and the maternal-fetal interface at the cellular level for hemochorial (a), epitheliochorial (d) and endotheliochorial (h) placenta, respectively. In hemochorial placentas, trophoblast cells invade both the maternal decidual stroma and spiral arteries, while in endotheliochorial placentas, trophoblast cells primarily disrupt maternal epithelial cells and make direct contact with endothelial cells. In epitheliochorial placentas, trophoblast cells are arranged parallel to the maternal epithelial cells. **b,e,f,i,** The interactions between EVT and other cells in different placental types. Genes involved in extracellular matrix construction/remodeling are marked in brown, those involved in cell invasion are marked in green, genes related to vascular development are marked in red, and genes associated with resistance to invasion are marked in purple. **c,g,j,** The interactions between immune cells and other cells in different placental types. Identical ligands and receptors are color-coded consistently.

The immune signaling landscape mirrored these structural differences. Hemochorial placentas were enriched ligand–receptor interactions promoting immune tolerance and anti-inflammatory macrophage function^54–58^, with APOE-TREM2 the significant pair and additional tolerance interactions including CD93–IFNGR1, IGFBP3–TMEM219, LGALS3–MERTK, and TGFB1–TGFBR3 (Fig. 3c). The APOE–TREM2 interaction recapitulates at the signaling-network level the APOE-dependent macrophage tolerance demonstrated *in vivo* above, indicating that *APOE* acts not only intrinsically within macrophages but also as a sender ligand to *TREM2* on neighboring cells. Epitheliochorial placentas instead upregulated pro-inflammatory signaling through APP–TNFRSF21, CD44–TYROBP, and TNFSF9–TNFRSF9 (Fig. 3g), amplifying inflammatory responses and creating barriers to trophoblast invasion^59–61^. Endotheliochorial placentas displayed a dual immune signaling profile, combining pro-inflammatory (IL18–IL18R1, IL6–IL6R, and CD44–TYROBP)^61,62^ and anti-inflammatory signaling (APOE–TREM2) (Fig. 3j), consistent with a balanced immune environment supporting moderate invasion.

These divergent communication architectures raised a question about the evolutionary logic of maternal-fetal signaling. Parent-offspring conflict theory^63^ predicts that deeper trophoblast invasion, by intensifying competition over maternal resources, should be accompanied by intensification of antagonistic signaling between fetal ligands and maternal receptors^4^. To test this prediction directly, we modeled the co-evolution of fetal ligands and their cognate maternal receptors across species using phylogenetically independent contrasts (PICs)^64^, classifying each ligand-receptor pair as antagonistic (negatively co-evolving), cooperative (positively co-evolving), or neutral. The result inverts Haig’s prediction: hemochorial placentas showed the smallest antagonistic ligand–receptor fraction (5.48%), epitheliochorial placentas the largest (30.76%), endotheliochorial placentas intermediate (16.67%) (Extended Data Fig. 9, and Supplementary Table 10). Across the morphotype spectrum, antagonism thus decreases progressively as invasiveness increased, the most invasive placentas exhibit the most cooperative signaling, while non-invasive placentas maintain the greatest molecular conflict. Rather than escalating conflict, deep invasion appears to resolve it through extensive immune tolerance mechanisms^65^, whereas superficial attachment preserves antagonistic signaling that restricts fetal access to maternal resources. This suggests that invasion depth itself may be an evolutionary solution to, rather than a cause of, maternal-fetal conflict.

Alternatively, these observations also admit a second theoretical reading. Antagonism predicted by Haig may not be eliminated but instead embedded within the tolerance circuitry itself, such that perturbation of tolerance, rather than overt antagonism, is the form in which conflict ultimately manifests. Under this reading, pregnancy disorders that compromise tolerance should fail in specific and predictable ways, exhibiting incomplete reversion toward less-invasive molecular states accompanied by deficits in the compensatory programs that normally sustain hemochorial pregnancy. We test this prediction in the following sections, in the contexts of preeclampsia and recurrent miscarriage.

### Preeclampsia recapitulates an epitheliochorial anti-invasion program without compensatory IGF signaling

Major pregnancy disorders associated with abnormal trophoblast invasion affect millions of pregnancies and remain incompletely understood at the cellular and molecular level. While Elliot and Crespi (2015)^21^ showed that preeclampsia risk loci recapitulate the genetic signatures of convergent invasiveness-reducing eutherian lineages, the cellular populations, regulatory programs, and compensatory features that distinguish viable epitheliochorial pregnancies from pathological reversion in human preeclampsia have remained unresolved. To address these questions, we generated snRNA-seq of human placenta and decidua from healthy control (∼39 weeks) and four major pregnancy complications, including fetal growth restriction (FGR, ∼37 weeks), gestational diabetes mellitus (GDM, ∼39 weeks), early-onset severe pre-eclampsia (PE, ∼28 weeks), and placenta accreta spectrum disorders (accreta, ∼37 weeks) (Supplementary Table 11). Across 158,852 high-quality cells (Supplementary Table 12), we identified all major cell types described above (Extended Data Fig. 10a–f) and used them to evaluate how each disorder deploys the cross-species hemochorial and epitheliochorial molecular features established in the preceding sections.

Clustering of trophoblast populations across disease conditions revealed disorder-specific lineage disruptions (Fig. 4a, c). In FGR, EVTs failed to cluster with control EVTs, instead grouping with the SCT lineage (Fig. 4c), consistent with arrested EVT differentiation. VCT and EVT from GDM, and SCT from PE, formed distinct clusters separate from their healthy counterparts (Fig. 4c), indicating altered trophoblast speciation or state. By contrast, all three trophoblast subtypes from accreta clustered appropriately with their healthy counterparts, suggesting that excessive invasion in acreta arises from functional alterations within correctly specified cell types rather than lineage mis-specification.

**Fig. 4.**
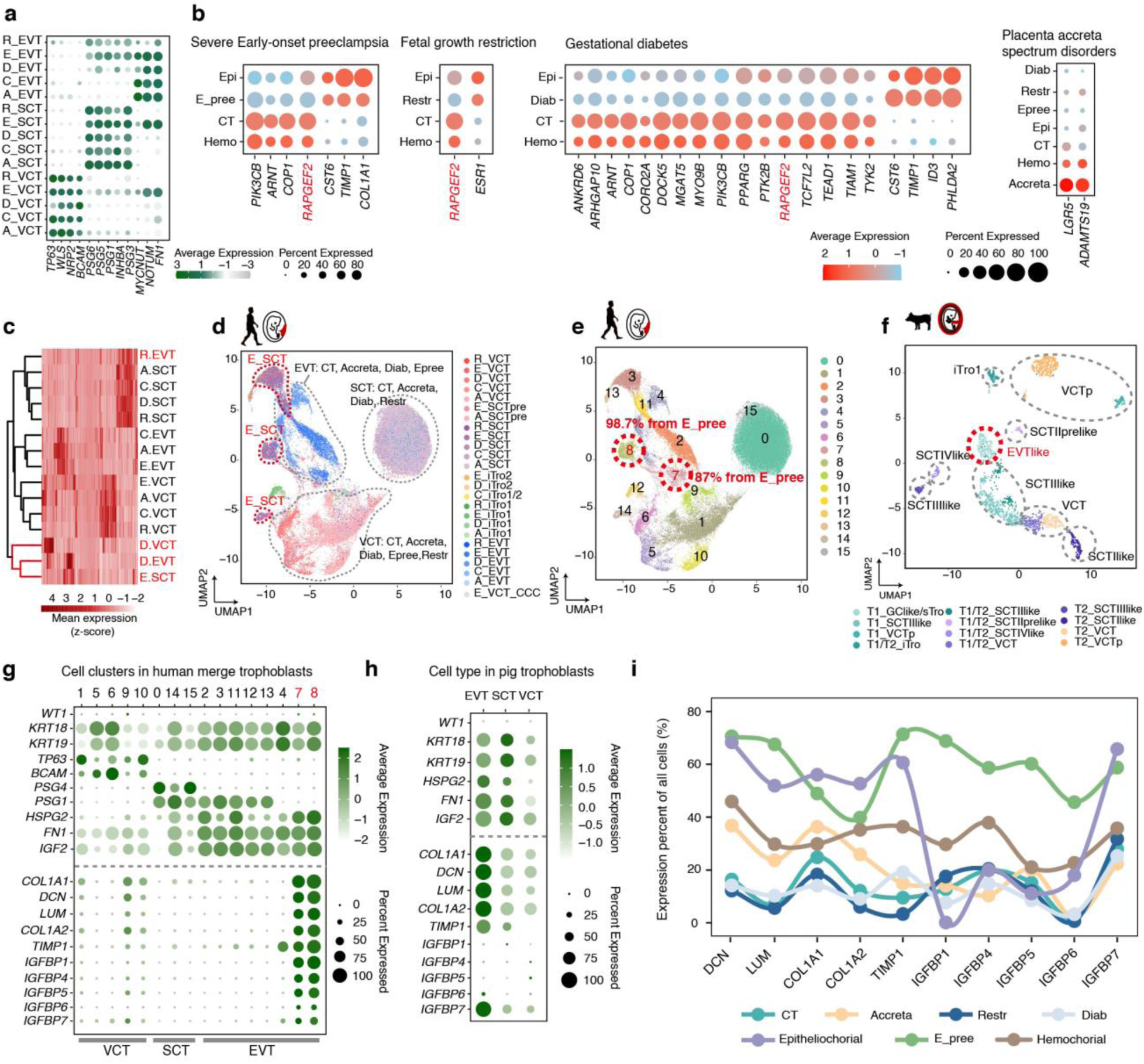
Integrated single-cell atlas of pregnancy disorders reveals essential genes in hemochorial placenta and commonalities with epitheliochorial placenta. **a,** Marker genes of trophoblasts (i.e., VCT, SCT, and EVT) for normal pregnancy (C_), fetal growth restriction (R_), gestational diabetes (D_), early-onset severe pre-eclampsia (E_) and placenta accreta spectrum disorders (A_). **b,** EVT differential expression analysis between hemochorial (Hemo) versus epitheliochorial (Epi) and normal pregnancy (CT) versus pregnancy-related diseases (i.e., fetal growth restriction (Restr), gestational diabetes (Diab), early severe pre-eclampsia (E_pree) and placenta accreta (Accreta)). **c,** Heatmap clustering of VCT, SCT, and EVT from normal pregnancy and different disease conditions based on marker gene expression. **d,** UMAP visualization of integrated trophoblasts from normal pregnancy and all disease conditions. Cell annotations in the UMAP are based on the cell-type classifications previously assigned within each individual disease dataset before integration. **e,** Cells re-clustering based on the integrated UMAP embedding, and the resulting cell populations are denoted by numeric identifiers. **f,** UMAP visualization of trophoblasts from pigs. **g,** Marker genes expression of all cell re-clusters of integrated trophoblasts from normal pregnancy and all disease conditions. **h,** Marker genes expression of trophoblasts in pigs. **i,** The fraction of all cells expressing marker genes of clusters c7 and c8 is shown across normal pregnancy, different disease conditions, hemochorial and epitheliochorial placentas.

Differential expression analysis in EVTs from three shallow-invasion disorders (FGR, GMD, PE) revealed that genes upregulated in healthy controls overlapped significantly with the hemochorial-enriched gene set defined cross-species above (Fig. 2d, 4b, Supplementary Tables 13 and 14), indicating that these disorders share a partial loss of the hemochorial molecular program. Among these, *RAPGEF2*, which encodes a guanine nucleotide exchange factor that activates the invasion-promoting GTPase Rap1, was consistently downregulated across all three shallow-invasion disorders^66^. Conversely, accreta EVTs showed marked upregulation of *LGR5*, a Wnt target gene associated with stemness and invasiveness^67–69^, and *ADAMTS19,* a metalloproteinase involved in extracellular matrix degradation^70,71^, consistent with their hyper-invasive phenotype of accreta. Additionally, based on the key hemochorial regulatory factors identified above—*AMZ1*, *GYS1*, *ELF4*, and *TEAD1*—we found that all four genes were significantly downregulated in GDM, whereas *AMZ1* and *GYS1* were significantly upregulated in accrete (Supplementary Table 13).

Among the disorders examined, preeclampsia is uniquely defined by shallow trophoblast invasion as its core pathological mechanism and carries the greatest maternal–fetal mortality burden globally^18^. To distinguish PE-specific molecular features from the shared shallow-invasion signature established above, we integrated trophoblasts from all disease conditions and re-examined cluster structure. PE SCTs segregated into three distinct populations, clustering separately with both EVT and VCT populations rather than forming a coherent SCT cluster (Fig. 4d). Re-clustering identified two novel sub-clusters (cluster 7 and cluster 8) composed almost exclusively of PE cells (i.e., 87% and 98.7%, respectively) (Fig. 4e). These clusters expressed canonical trophoblast markers (i.e., *KRT18* and *KRT19*) and EVT markers (i.e., *HSPG2*, *FN1* and *IGF2*), but additionally expressed of a set of marker genes shared with porcine epitheliochorial EVT-like trophoblasts including *COL1A1*, *DCN*, *LUM*, *COL1A2*, *TIMP1* and *IGFBP7* (Figs. 4f–4h). This suggested that preeclampsia trophoblasts had adopted a molecular program resembling the anti-invasive regulatory program employed by high degree of similarity to the EVT-like trophoblast phenotype observed in pigs.

To test whether this epitheliochorial-like signature extended beyond trophoblasts, we examined expression of the c7/c8 markers across all cell types in each condition. In healthy controls and other pregnancy disorders, these genes were predominantly expressed in stromal populations (fetal and maternal stromal cells) as expected (Extended Data Fig. 11a–d and 12). In PE, however, they showed broad, elevated expression across decidual macrophages, decidual NK cells, maternal endothelial cells, and maternal epithelial cells (Extended Data Fig. 11a, b and 12). The fraction of cells expressing these genes across the whole placenta in PE (an average of 59.66%) approached the levels seen in porcine epitheliochorial placentation (an average of 57.94%), and exceed the levels seen in hemochorial placentation (an average of 35.43%), healthy controls (an average of 13.93%), or other pregnancy-related disorders (an average of 9.08%-27.51%) (Fig. 4i and Extended Data Fig. 11a–f). Functionally, *COL1A1*, *DCN*, *LUM* and *COL1A2* promote collagen deposition and inhibit matrix remodeling^72–74^, *TIMP1* inhibits matrix metalloproteinase^75^, and *IGFBP7* promotes fibrotic microenvironments^76,77^, together constituting a collagen-rich, anti-invasive matrix program. Excessive collagen deposition is a well-documented feature of human preeclamptic placentas, and collagen I is sufficient to suppress trophoblast proliferation and invasion experimentaly^78^. PE therefore deploys, across multiple cell types, the same matrix-stabilizing molecular program that operates as the stable evolutionary regulatory program of epitheliochorial placentation cross-species.

Yet PE and epitheliochorial placentation diverged in one critical respect, Despite their shared collagen-rich, anti-invasive matrix signature, PE showed markedly elevated expressions of the insulin-like growth factor binding proteins, *IGFBP1*, *IGFBP4*, *IGFBP5*, and *IGFBP6* relative to epitheliochorial placentas (Fig. 4i). These binding proteins sequester IGFs and suppress IGF signaling^79^, which is essential for trophoblast survival, metabolism, and endocrine function. In epitheliochorial placentation, matrix-mediated invasion restriction is paired with intact IGF signaling that sustains trophoblast viability. PE combines the same matrix restriction with simultaneous IGF sequestration, generating a doubly restrictive environment that impairs both invasion and trophoblast function. Together, these results specify what distinguishes a viable evolutionary configuration from a pathological one. Preeclampsia recapitulates the matrix-stabilizing regulatory program that pigs, sheep, and goats have convergently evolved as an anti-invasive epitheliochorial state, but without the compensatory IGF-axis tuning that makes shallow trophoblast invasion compatible with successful pregnancy in those species.

### Recurrent miscarriage recapitulates epitheliochorial immune architecture without compensatory fetal responses

Early recurrent miscarriage (ERM) affects 1-3% of pregnancies^80^ and is characterized by reduced trophoblast invasion and elevated maternal immune activation^81^. Epitheliochorial mammalian species sustain successful pregnancies despite similar immunological features, raising the question of whether ERM reflects failure of hemochorial-specific mechanisms or incomplete recapitulation of an epitheliochorial-like state, and what compensatory features distinguish viable epitheliochorial pregnancy from pathological ERM in humans. Previous study revealed that invasive trophoblasts exhibited significant association with recurrent pregnancy loss^82^, but the relationship between this signature and the immune architecture of less-invasive mammalian placentation has not been examined. To address these questions, we performed compared the immune cell composition from scRNA-seq of ERM placentas^83^ with our cross-species data (Methods).

The analyses revealed coherent patterns distinguishing hemochorial from non-hemochorial placentation. Fetal macrophages (HB) in hemochorial placentas were exclusively M2-type (anti-inflammatory) (Fig. 5a, and Supplementary Table 15), while non-hemochorial placentas contained both M1 (pro-inflammatory) (average 5.57% of total immune cells) and M2 (∼18.65%) populations (Supplementary Table 15). Maternal macrophages (dM) followed a parallel pattern, with M1 cells comprising ∼14.42% of immune cells in non-hemochorial placentas but only ∼7.65% in hemochorial species (Extended Data Fig.13a, and Supplementary Table 15). The lymphoid compartment displayed an inverse pattern. Hemochorial placentas exhibiting remarkable lymphoid diversity, with five distinct dNK /dNKT/T cell subtypes comprising 40.38% of immune cells on average, whereas non-hemochorial placentas showed only 1-3 subtypes, representing just 6.75% of immune populations (Fig. 5a, and Supplementary Table 15). We also detected two trophoblast subpopulations expressing immune markers across species (Extended Data Fig. 5b–o). One subset (iTro1) mainly expressed macrophage marker genes and clustered with dM cells, while another (iTro2) predominantly expressed NK cell marker genes and clustered with dNK cells. Notably, iTro2 appeared exclusively in hemochorial placentas (Extended Data Fig. 13b), suggesting it represents an adaptation specific to deep invasion.

**Fig. 5.**
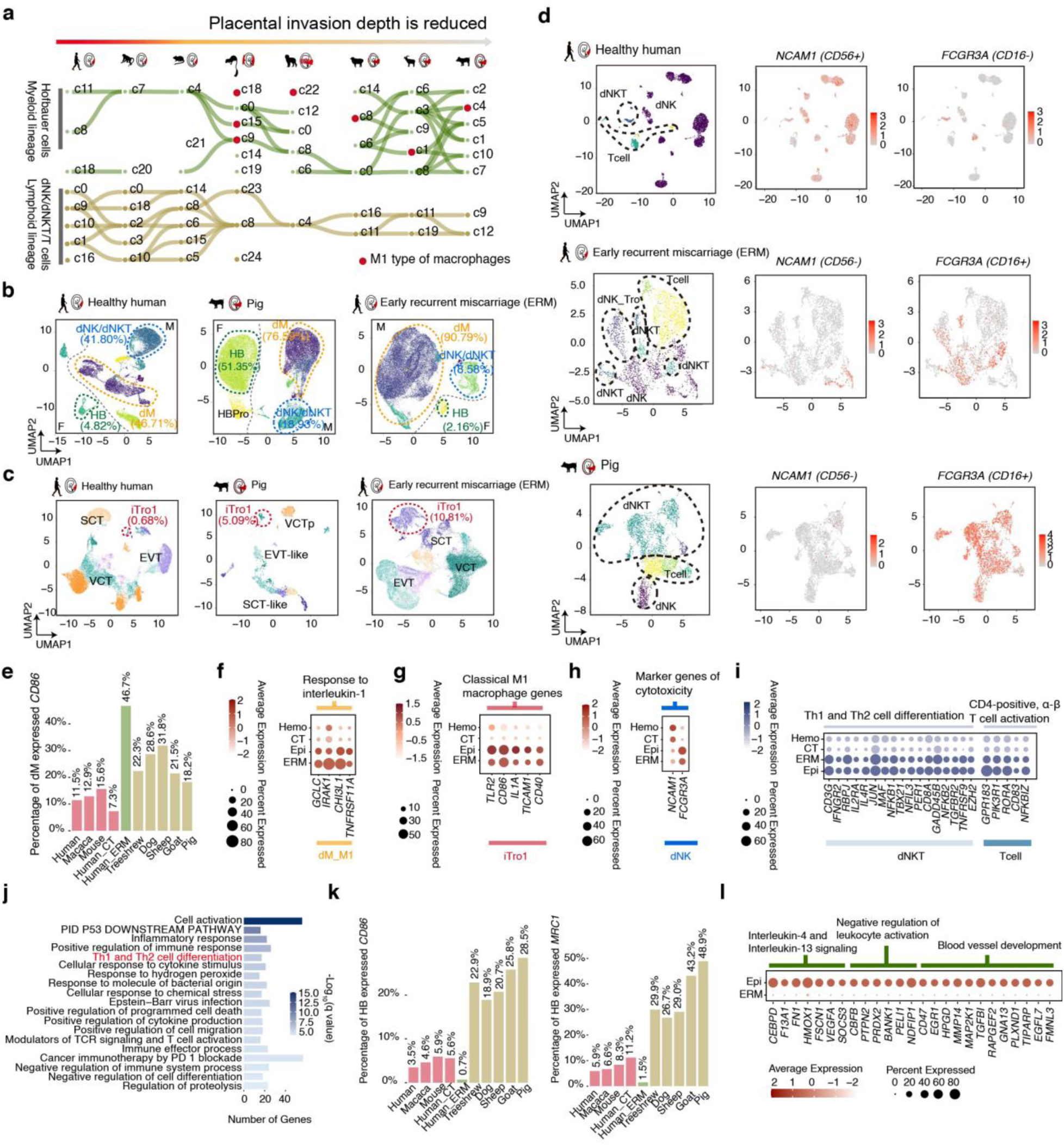
Immune cell expression characteristics across different species and early recurrent miscarriage (ERM). **a,** SAMap mapping relationships of myeloid lineage and lymphoid lineage between eight mammalian species with an alignment threshold of 0.3. **b,c,** UMAP visualization of immune cells (b) and trophoblasts (c) in healthy human, pigs and ERM, respectively. **d,** UMAP visualization of lymphoid lineage, and expression of *NCAM1* (*CD56*) and *FCGR3A* (*CD16*) in healthy human, pigs and ERM, respectively. **e,** The proportion of *CD86*⁺ decidual macrophages (dM) among immune cells across eight mammalian species, normal controls (CT), and ERM. **f,g,h,i,** Overlapped up-regulated genes expression between epitheliochorial placentas and ERM for M1 type decidual macrophages (dM_M1) (f), iTro1 (g) and decidual natural killer cells (dNK) (h), dNKT and T cells (i), respectively. **j,** Pathway enrichment analysis of overlapping upregulated genes in dNK cells between epitheliochorial placentas and ERM. **k,** The proportion of *CD86*⁺ and *MRC1*^+^ Hofbauer (HB) among immune cells across eight mammalian species, normal controls (CT), and ERM. **l,** Upregulated genes in epitheliochorial HB cells *versus* ERM, which are significantly enriched in Interleukin-4 and Interleukin-13 signaling, negative regulation of leukocyte activation, and blood vessel development pathways.

Applying this cross-species comparison to ERM revealed striking immunological convergence with epitheliochorial placentation rather than healthy humans, particularly in maternal immune cell composition and inflammatory status. Decidual macrophages (dM) dominated the maternal immune compartment in both pigs (76.59%) and ERM placentas (90.79%), whereas they comprised only 46.71% in healthy human placentas (Fig. 5b). By contrast, decidual NK (dNK) and dNKT cells were markedly enriched in healthy humans (41.80%), but were significantly reduced in pigs (18.93%) and ERM (8.58%) (Fig. 5b). The immune-like trophoblasts (iTro1) were similarly enriched in pigs (5.09%) and ERM placentas (10.81%) compared with healthy humans (0.68%) (Fig. 5c), indicating a shared inflammatory trophoblast phenotype between non-hemochorial placentation and pathological human pregnancy.

At the functional level, dM cells from both non-hemochorial placentas and ERM samples exhibited elevated expression of the M1 pro-inflammatory marker *CD86* (Fig. 5e), along with significant enrichment of interleukin-1 response pathway genes (Fig. 5f, and Supplementary Table 16). iTro1 cells from both epitheliochorial and ERM placentas showed high expression of classical M1 macrophage markers (Fig. 5g), supporting a pro-inflammatory trophoblast phenotype. The lymphoid compartment reinforced this inflammatory signature. dNK cells from pigs and ERM placentas displayed reduced expression of *NCAM1* (*CD56*), a marker of cytotoxic restraint and immune tolerance, with concomitant upregulation of *FCGR3A* (*CD16*) (Fig. 5d, h, and Supplementary Table 16), which enhances cytotoxic and antibody-dependent cellular activity. Moreover, genes shared between epitheliochorial and ERM dNK/dNKT/T cells were significantly enriched in immune activation pathways, including inflammatory response, Th1 and Th2 cell differentiation, and CD4-positive αβ T cell activation, further indicating a pro-inflammatory immune milieu (Fig. 5i, j). Together, these findings indicated that ERM placentas had adopted a pro-inflammatory immune environment characteristic of epitheliochorial placentation, dominated by activated macrophages and cytotoxic NK cells rather than the tolerogenic populations that characterize healthy hemochorial pregnancy.

This convergence raised a critical question that if ERM placentas recapitulate immunological architecture of epitheliochorial placentation, why does epitheliochorial placentation sustain successful pregnancy across multiple mammalian species while ERM fails? Our results indicate that, despite their similarities on the maternal side, ERM and epitheliochorial placentas display differed dramatically in fetal macrophage populations. In pigs, fetal macrophages (HB) constitute a substantial fraction of total immune cells (51.35%) (Fig. 5b), representing a robust fetal immune presence. In ERM placentas, by contrast, HB cells only constitute a small proportion (2.16%) of immune populations (Fig. 5b). This depletion affected both M1-like (CD86⁺) and M2-like (MRC1⁺ and ARG1/2^+^) subsets (Fig. 6k, and Supplementary Table 17), indicating a complete collapse of the fetal macrophage compartment rather than a polarization shift between subsets.

**Fig. 6.**
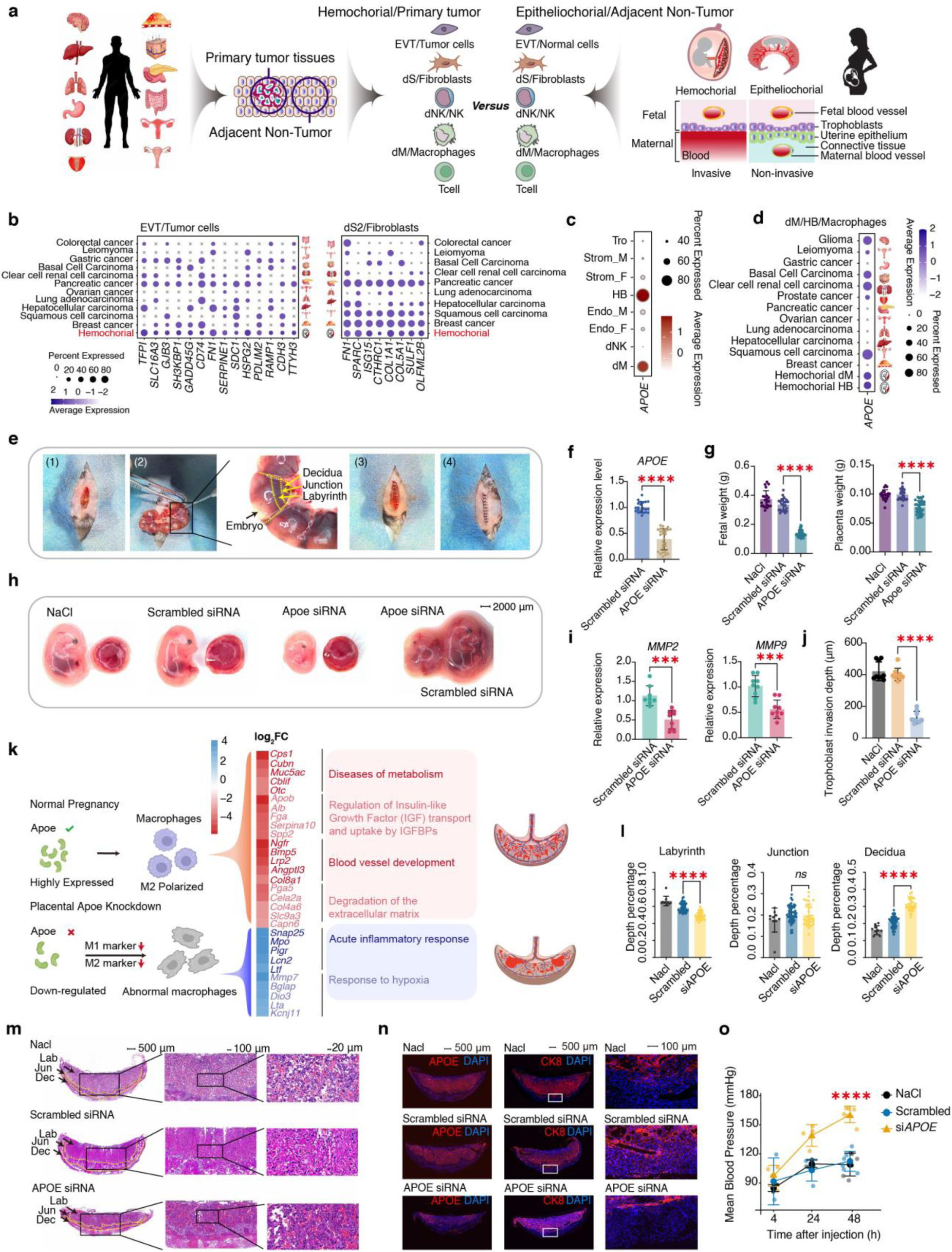
Comparative differential expression analysis of cell populations across hemochorial and epitheliochorial placenta types and tumor tissue states. **a,** Schematic illustration of cell-type–specific comparisons between tumor tissues and different placental types in this study. **b,** Expression of genes that are upregulated in at least five tumor types and overlap with hemochorial-upregulated genes across EVT/tumor cells and dS2/fibroblasts, respectively. **c,** Expression of *APOE* across cell types in the mouse placenta. **d,** Expression of genes that are upregulated in at least five tumor types and overlap with hemochorial-upregulated genes across macrophages/HB/dM. **e,** Uterine laparotomy procedure. The procedure included shaving the dam’s abdomen, making an approximately 2-cm incision in the abdominal skin (1), exteriorizing and gently manipulating the uterine horns through the incision and injecting siRNA into E12.5 placentas (2), followed by complete suturing of the peritoneum (3) and the abdominal skin (4). **f**, qPCR analysis of *APOE* expression in E15.5 placentas treated with scrambled siRNA (negative control, *n* = 7 with three technical replicates) and APOE siRNA (*n* = 9 with three technical replicates). **g,** Fetal and placental weights at E15.5 following normal saline (NaCl, vehicle control, *n* = 21), scrambled siRNA (negative control, *n* = 25), and *APOE* siRNA treatment (*n* = 37). **h,** Representative gross morphology and quantification of E15.5 embryos and placentas from NaCl, scrambled siRNA, and *APOE* siRNA treated groups. **i,** qPCR analysis of *MMP2*, and *MMP9* in E15.5 placentas treated with scrambled siRNA (negative control, *n* = 3 with three technical replicates) and *APOE* siRNA (*n* = 3 with three technical replicates). **j,** Quantification of trophoblast invasion depth in E15.5 placentas from NaCl, scrambled siRNA, and *APOE* siRNA treated groups. **k,** Schematic overview of *APOE* function during placental development. **l,** Relative depth (percentage) of the labyrinth, junctional zone, and decidua in E15.5 placentas across NaCl (*n* = 50), scrambled siRNA (*n* = 50), and *APOE* siRNA (*n* = 30) groups. **m,** Hematoxylin and eosin (H&E) staining of E15.5 placentas from NaCl, scrambled siRNA, and *APOE* siRNA treated groups. Black arrowheads indicate the labyrinth (Lab), junctional zone (Jun), and decidua (Dec). **n,** Immunofluorescence staining for APOE (red), CK8 (red) of E15.5 placental sections at 8μm thickness from NaCl, scrambled siRNA, and *APOE* siRNA-treated groups. **o,** Systolic blood pressure (SBP) in pregnant mice across NaCl (*n* = 12), scrambled siRNA (*n* = 13), and *APOE* siRNA (*n* = 14) groups at 4, 24, and 48 hours after injection at E12.5. Statistical significance was assessed using an unpaired t test comparing scrambled siRNA (negative control) and *APOE* siRNA groups. For Systolic blood pressure (SBP), all data are presented as the mean ± SEM. Statistical significance was determined using a two-way ANOVA with Sidak’s multiple comparisons test. Asterisks indicate significance levels: \*\*\*\**P* < 0.0001, \*\*\**P* < 0.001, \*\**P* < 0.01.

Differential expression analysis further revealed that genes enriched in epitheliochorial HB cells but downregulated or absent in ERM were significantly associated with Interleukin-4 and Interleukin-13 signaling, negative regulation of leukocyte activation, and blood vessel development (Fig. 5l). These pathways regulate macrophage M1-to-M2 polarization, immune tolerance, and placental vascular remodeling. Their near-complete loss in ERM-associated HB cells suggests that fetal macrophages in epitheliochorial species actively counterbalance maternal inflammation, creating a permissive environment for pregnancy despite elevated maternal immune activation. ERM thus represents not simply excessive maternal immune activation but an incomplete recapitulation of epitheliochorial immune architecture. Epitheliochorial species have evolved into a balanced system where maternal pro-inflammatory responses are offset by robust fetal macrophage populations that promote tolerance and vascular development. ERM adopts the maternal inflammatory component without the compensatory fetal response, creating an immunologically hostile environment that cannot support pregnancy.

### Placenta–cancer convergence identifies *APOE* as regulator of hemochorial invasion

The immune-tolerance mechanisms that permit deep trophoblast invasion in hemochorial pregnancy, and whose breakdown underlies ERM, are also a defining feature of invasive malignancy^84^. This parallel, together with the cancer-pathway enrichment and shared invasion-promoting regulators we identified in the hemochorial program, prompted us to test systematically the parallels between trophoblast and tumor invasion across cellular compartments, and to ask whether hemochorial placentation and cancer converge not only on invasion but also on shared mechanisms of immune tolerance. Yu et al. (2024)^28^ recently integrated single-species (human) placental scRNA-seq with multi-cancer scRNA-seq and identified the onco-fetal molecule B7-H4 as a shared immune-tolerance checkpoint, establishing the integration approach but leaving open whether the convergence is hemochorial-specific or generic across placental morphotypes and whether it has cross-species evolutionary structure. To address these questions, we integrated our cross-species placental data with single-cell transcriptomes from 13 human tumor types (∼ 894,815 cells from primary tumors and adjacent non-tumor tissues (Supplementary Table 18)), and compared transcriptional profiles between matched cell-type pairs across placentas and tumors (Fig. 6a, and Supplementary Table 19).

The comparison revealed remarkable transcriptional convergence across multiple cellular compartments. Across all cell types examined, genes upregulated in tumor tissues showed significant overlap with genes upregulated in hemochorial placentas compared to epitheliochorial placentas (Extended Data Fig. 14). This pattern held consistently across diverse cancer types, demonstrating that the molecular signature distinguishing invasion from non-invasive placentation extends beyond the invasive cells to encompass the entire tissue microenvironment. In this framework, we define cancer invasiveness as the capacity for local tissue invasion, the regulated remodeling and degradation of the extracellular matrix and migration through stromal barriers, which constitutes the initiating step of the invasion–metastasis cascade^85^. This is also the program most directly comparable to trophoblast behavior, since extravillous trophoblasts likewise execute controlled local invasion into the maternal decidua without distant dissemination^86^.

Focusing on genes consistently upregulated in at least five tumor types that were also elevated in hemochorial placentas (Fig. 6b, d), we characterized the shared programs of two compartments in detail. Within EVT/tumor cells, the shared invasion machinery upregulated was enriched for hemostasis (e.g., *SDC1*, *TFPI*, *FN1*, and *SERPINE1*), positive regulation of cell motility (e.g., *CD74*, *FN1*, and *SERPINE1*) and extracellular matrix organization (*FN1*, *HSPG2*, *SERPINE1*, *SDC1*) (Fig. 6b, and Supplementary Table 20), supporting matrix remodeling, cytoskeletal rearrangement, and force generation required for both trophoblast invasion into maternal decidua and tumor cell dissemination through stromal barriers. Within the stromal pair, decidual stromal cells in hemochorial placentas and cancer-associated fibroblasts shared upregulation of ECM organization genes (*COL1A1*, *COL5A1*, *SULF1*, *OLFML2B*, and *CTHRC1*) and positive regulation of cell migration (*COL1A1*, *FN1*, and *SPARC*) (Fig. 6b, and Supplementary Table 20), supporting stromal remodeling and formation of a microenvironment permissive to invasive growth. These multi-compartment overlaps are consistent with a refinement of the long-standing trophoblast-cancer convergence framework^12,15,22^ from scattered single-molecule co-option toward coordinated redeployment of the multi-cellular invasion program characteristic of mammalian hemochorial placentation.

Interestingly, among all cancer types examined, pancreatic ductal adenocarcinoma displayed the largest number of genes shared with the hemochorial regulatory program across both the EVT-tumor and stromal-fibroblast compartment (Fig. 6b). The PDAC–hemochorial overlap encompassed both the invasion machinery shared in the EVT–tumor pair and the stromal program shared with cancer-associated fibroblasts. This dual-compartment concordance is consistent with PDAC’s well-characterized desmoplastic invasive phenotype, in which coordinated remodeling of stromal extracellular matrix sustains aggressive tumor-cell invasion^87^, paralleling the coordinated trophoblast invasion and maternal stromal remodeling characteristic of hemochorial placentation.

Immune tolerance is another shared feature of both hemochorial placentation and aggressive tumor microenvironments, with macrophages central to its establishment in both contexts^88,89^. Consistently, decidual/Fetal macrophages and tumor-associated macrophages across multiple cancer types all upregulated *APOE* (Fig. 6d, and Supplementary Table 21), a gene involved in lipid metabolism and macrophage polarization toward anti-inflammatory states, contributing to immune tolerance^90^. *APOE*+ macrophages are now well-established as an immunosuppressive population across multiple cancer types, including pancreatic, hepatocellular, lung, renal, and colorectal cancers, where they support tumor progression and immunotherapy resistance through anti-inflammatory polarization, regulatory T cell recruitment, and CD8+ T cell suppression^91–93^. The placental *APOE*+ macrophage population has received far less attention and its prior literature in human pregnancy disorders is limited to genotype-severity association in preeclampsia^94^ and transcriptomic differential expression in PE Hofbauer cells^95^, with no prior cell-autonomous functional perturbation in the placenta. The cross-compartment convergence in tolerance-associated macrophage populations nominated *APOE* for functional validation *in vivo*.

To test whether *APOE* is functionally required for trophoblast invasion *in vivo*, and given its preferential expression in macrophages relative to other cell types (Fig. 6c), we delivered *APOE-*targeting siRNA (or scrambled control) directly into mouse placentas by uterine laparotomy at E12.5 (Fig. 6e, and Extended Data Fig. 15a) and assessed phenotypes at E15.5. The siRNA achieved efficient *APOE* knockdown in placental tissue at both the transcript level by qPCR (Fig. 6f, and Supplementary Table 22) and the protein level by immunofluorescence and western blot (Fig. 6n, and Extended Data Fig. 15b, c). Placenta-targeted *APOE* knockdown reduced both fetal and placental weight significantly (Fig. 6g, and Supplementary Table 23) and altered embryo and placental gross morphology (Fig. 6h and Extended Data Fig. 16). Histological analysis of E15.5 placentas by hematoxylin and eosin (H&E) staining showed disrupted placental architecture (Fig. 6m, and Extended Data Fig. 17). Quantitatively, placentas exhibited a pronounced thinning of the labyrinth zone accompanied by a marked expansion of the decidual layer (Fig. 6l, and Supplementary Table 24), and CK8^+^ trophoblasts invasion depth into the decidua was significantly fewer and shallower compared with controls (Fig. 6n, j and Extended Data Fig. 18). The reduced trophoblast invasion was accompanied by reduced placental expression of the invasion-promoting markers (*MMP2* and *MMP9*) (Fig. 6i). Notably, pregnant mice with placental *APOE*-knockdown exhibited a significantly elevated systolic blood pressure (mean, 160 mmHg) relative to scrambled siRNA controls (mean, 112 mmHg) (Fig. 6o). Together, these phenotypes recapitulate the cardinal features of human preeclampsia, with fetal growth restriction, impaired trophoblast invasion, and maternal hypertension, demonstrating that placental *APOE* is functionally required for normal trophoblast invasion and maternal cardiovascular homeostasis in hemochorial pregnancy.

Given the critical role of *APOE* in macrophage development, polarization, and the establishment of immune tolerance, we hypothesized that placental *APOE* sustains invasion by maintaining the polarization state of resident macrophages. We tested this by measuring canonical M1 (pro-inflammatory) and M2 (anti-inflammatory) macrophage program markers in *APOE*-knockdown placentas (Extended Data Fig.19, and Supplementary Table 22), and found *APOE*-knockdown coordinately suppressed both M1 and M2 gene programs. This dual suppression is consistent with either coordinated failure of both polarization programs or coordinated loss of placental macrophage abundance under *APOE* loss, a distinction that single-cell perturbation analysis will resolve. In either case, placental *APOE* is required to sustain an immune-tolerant macrophage compartment at the maternal-fetal interface. Notably, this phenotype closely mirrors macrophage function and immune tolerance described in early recurrent miscarriage, and is consistent with a model in which placental *APOE* sustains an immune-tolerant macrophage compartment that permits normal trophoblast invasion. Bulk RNA-seq profiling of scrambled-control *versus APOE*-knockdown placentas provided further mechanistic context (Fig. 6k, Extended Data Fig.20a, b, and Supplementary Table 25). In scrambled controls, the most significantly upregulated pathways included blood vessel development, regulation of IGF transport and uptake by IGF-binding proteins, extracellular matrix degradation, and diseases of metabolism, programs collectively consistent with the normal vascular elaboration, trophoblast invasion, and fetal growth support observed in this group. Conversely, *APOE* knockdown placentas showed preferential upregulation of response to hypoxia and acute inflammatory response pathways (Fig. 6k, Extended Data Fig. 20b, and Supplementary Table 26), indicating that loss of placental *APOE* triggers a shift from a growth- and invasion-permissive transcriptional state toward hypoxic stress and sterile inflammation. Together, these transcriptional signatures are consistent with a model in which *APOE* coordinates an integrated lipid–macrophage–vascular–invasion axis: lipid-metabolic failure under *APOE* loss arrests macrophage function, inhibits blood vessel development, attenuates invasion, and provokes hypoxia-driven inflammation, although whether the macrophage and trophoblast phenotypes are causally linked or arise as parallel *APOE*-dependent outputs will require macrophage-specific perturbation to distinguish.

Together, these findings establish that the transcriptional programs distinguishing hemochorial from epitheliochorial placentation are recapitulated across multiple cell types in the tumor microenvironment, and that one of the regulators nominated by this cross-compartment convergence, *APOE* in tolerance-associated macrophages, is functionally required for normal hemochorial trophoblast invasion and maternal cardiovascular homeostasis *in vivo*. Token together, the multi-compartment transcriptional convergence and the functional validation of *APOE* support the evolutionary hypothesis that cancer invasion involves coordinated reactivation of deeply conserved developmental programs that mammalian evolution has optimized for controlled tissue invasion.

## Discussion

The evolution of placental invasion presents a paradox in which fetal tissues must invade maternal tissues to establish nutrient exchange while invasion is controlled to prevent pathology^18,96–98^. Haig’s parent-offspring conflict theory^4^ predicts that invasion depth reflects the equilibrium between fetal extraction pressures and maternal countermeasures. Recent comparative single-cell work established the conserved cellular architecture of the maternal-fetal interface between eutherian hemochorial mammals and non-invasive marsupials^24^, and the cross-species regulatory landscape of trophoblast diversification at late gestation^26^. Our single-cell atlas across mammalian species representing all major placental types at the active-invasion window of mid-gestation extends these prior studies by showing that conserved cellular machinery is deployed under three distinct regulatory programs, cancer-like invasion, endothelial-cooperative programs, and collagen-rich extracellular matrix stabilization, representing multiple evolutionary solutions to the invasion-control problem. Cross-species phylogenetically-independent contrasts of ligand-receptor co-evolution invert Haig’s prediction. The most invasive placentas show the lowest antagonistic ligand-receptor fraction and the most cooperative signaling, while non-invasive placentas maintain the greatest antagonism. Together with the higher expression-evolution rates of fetal invasion lineages relative to maternal stromal and immune lineages, consistent with directional versus stabilizing selection, these findings suggest that deep invasion evolved not through escalated conflict but through mutual disarmament via coordinated immune tolerance, positioning invasion depth as an evolutionary solution to maternal-fetal conflict rather than a cause of it.

This evolutionary perspective transforms our understanding of pregnancy disorders from isolated pathologies into predictable consequences of regulatory circuit dysfunction. Our work specifies the cellular and molecular features being recapitulated and identifies the species-typical compensations whose absence distinguishes viable epitheliochorial pregnancy from pathological deployment in humans. Our study revealed that preeclampsia exhibits transcriptional convergence with epitheliochorial placentation through collagen-rich, anti-invasive matrix programs, but additionally suppress IGF signaling that epitheliochorial species maintain to sustain trophoblast viability under shallow invasion, creating a doubly restrictive environment that impairs both invasion and trophoblast function. Similarly, recurrent miscarriage recapitulates the pro-inflammatory maternal immune environment of epitheliochorial placentation, but lacks the robust fetal macrophage population whose tolerance-promoting and pro-vascular gene programs counterbalance maternal inflammation in epitheliochorial species. These patterns support an evolutionary mismatch model^99^ in which pathology disorders arise not from random molecular failures but from activation of evolutionarily coherent invasion control programs in inappropriate species contexts^100,101^, with critical co-evolved compensations failing to engage. This framework generates testable hypothesis that restoring the missing compensations may be a therapeutic strategy worth investigating for these disorders.

The convergence between placental invasion and cancer metastasis extends this evolutionary logic to oncology. The trophoblasts-tumors paralle^12,15,22^ has typically been interpreted as cancer hijacking placental programs, and recent single-species placenta-tumor scRNA-seq integration identified the onco-feta molecule B7-H4^28^. Our cross-species integration across thirteen cancer types extends this work in two respects. First, cancers recapitulate not isolated invasion molecules but coordinated multi-cellular transcriptional programs encompassing invasion machinery, stromal remodeling, and immune tolerance. Pancreatic adenocarcinoma shows the strongest concordance with hemochorial placentation across both invasive and stromal compartments. This multi-cellular convergence suggests that reactivation of ancient tissue invasion circuits contributes to cancer metastatic capacity alongside cancer-specific genetic alterations^16^. Second, the cross-compartment convergence identified *APOE* as a shared regulator across decidual, fetal, and tumor-associated macrophages. *APOE*+ macrophages are well established as immunosuppressive populations in cancers^92,102,103^, but no cell-autonomous *in vivo* perturbation in the placental context had been performed. Our demonstration that placental *APOE* knockdown impairs trophoblast invasion and produces a preeclampsia-like maternal phenotype establishes *APOE*-mediated immune tolerance as a functionally required component of hemochorial program shared with aggressive cancers. The cross-species context also places the VGLL3-TEAD1 axis identified by Plazyo et al. (2026)^36^ as dysregulated in human preeclampsia within a broader framing, positioning the disorder as perturbation of a conserved hemochorial regulator rather than a disease-specific molecular event. Cancers therefore appear to redeploy not isolated invasion molecules but the coordinated invasion-tolerance machinery mammalian evolution has shaped for controlled maternal-fetal coexistence.

## Methods

### Samples processing, Ethics statement, single-cell/nucleus sequencing

The experimental pregnant animals in this study include Sugar glider (*Petaurus breviceps*), C57BL/6J mouse (*Mus musculus*), Chinese tree shrew (*Tupaia belangeri chinensis*), Beagle dog (*Canis lupus familiaris*), Hu sheep (*Ovis aries*), Shanghai Chongming White goat (*Capra hircus*) and Suziblack pig (*Sus scrofa domestica*) at early and middle gestation points. Time point chosen were in sugar glider day 9 (*n* = 1) of 15 total gestation, in mouse 10.5 (*n* = 2) and 15.5 (*n* = 2) of 19.5, in tree shrew day 15 (*n* = 3) and 25 (*n* = 3) of ∼45, in dog day 25 (*n* = 3) and 40 (*n* = 3) of ∼63, in sheep day 60 (*n* = 1) and 86 (*n* = 1) of ∼147, in goat day 45 (*n* = 3) and 75 (*n* = 3) of ∼150, and in pig day 30-31 (*n* = 3) and 63-68 (*n* = 3) of ∼114 days gestation. The placenta and decidua/endometrium were collected respectively (Extended Data Fig. 1), and kept at 4℃ in MACS® Tissue Storage Solution for approximately 24h until they could be dissociated into single-cell suspension. Sugar glider was purchased from pet markets, C57BL/6J mice were purchased in Hangzhou hangsi Biotechnology Co., Ltd, tree shrews were obtained at Kunming Institute of Zoology, Chinese Academy of Sciences, sheep and goats were collected at Nanjing Zhushun Biotechnology Co., Ltd. and pigs were sampled at Taizhou Taihe Biotechnology Co., Ltd. The beagle dogs were obtained from local veterinary hospital. After proper animal restraint and anesthesia in the hospital, laparotomy was performed. Following removal of the uterus, the abdominal incision was sutured, and placental and decidual tissues were collected from uterus. Postoperative care and recovery procedures were carried out accordingly. Experimental protocols and animal use in this study were approved by the Zhejiang University Ethics Committee Institutional Review Board (ZJU20230191 and ZJU20240087). Healthy human (*Homo sapiens*) placenta and decidua at gestation of 14 week were obtained at Women’s Hospital, Zhejiang University, with appropriate maternal written consent and approval form the Women’s Hospital, Zhejiang University Ethics Committee (IRB-20250065-R). These were integrated with a human atlas generated from 4-13 weeks^104^, cynomolgus macaque samples generated from stages E29 and E45 of a 162-day gestation^105^, early recurrent miscarriage samples^83^, gestational diabetes^106^ and public cancer scRNA data (Supplementary Table S18).

We additionally collected full-term placenta and decidua from control subjects (*n* = 2) and patients were diagnosed as fetal growth restriction (*n* = 1), severe preeclampsia (*n* = 2), gestational diabetes (*n* = 1) and placenta accreta spectrum disorders (*n* = 3) at Women’s Hospital, Zhejiang University to perform single-nucleus RNA sequencing. Written informed consent was obtained from all participants, and the study was approved by the Ethics Committee of the Women’s Hospital, Zhejiang University (IRB-20250065-R). The fetal growth restriction case had no history of chronic hypertension or pregestational diabetes and was confirmed by ultrasound examination. Gestational diabetes was diagnosed according to “Guideline No. 393-Diabetes in Pregnancy”^107^ and pregnant women with obesity or undergoing insulin treatment were excluded. Severe preeclampsia samples were classified according to American College of Obstetricians and Gynecologists guidelines, based on high blood pressure (systolic > 160 mm Hg or diastolic > 100 mm Hg), thrombocytopenia, impaired liver function, renal insufficiency, pulmonary edema and/or cerebral/visual disturbances. For the case diagnosed as increta, the ultrasound profile met the criteria of International Federation of Gynecology and Obstetrics (FIGO). The PAS placenta and decidua were obtained from the removed invaded myometrial tissue, where the invaded myometrial tissue was resected, and myometrial reconstruction was performed. Maternal parameters for each cohort are detailed in Supplementary Table 11.

To generate scRNA-seq data, placenta and decidua/endometrium were minced with scalpel into 0.1∼0.2g cubes in 2 ml of digestive solution containing 0.2 mg/ml Liberase^TM^ TM (Roche, 5401119001) and 0.25% Trypin-EDTA (Gibco, 25200056). Tissue suspensions were heated at 37℃ for 1∼2h and the digestion was terminated by adding 2ml of fetal bovine serum (FBS) (Australia Origin) (Zeta Life, Z7010FBS-500). The disaggregated cell suspension was passed through a 40-μm cell strainer. The flow-through was pelleted by centrifugation at 700 rcf for 5 min and cells were resuspended in 5ml of red blood cell lysis buffer (Invitrogen, 00-4300) for 10 min at ice. At this point, cells were examined on C100 Automated Cell Counter to check for debris and cell death through 0.4% trypan blue stain (Thermo Fisher, T10282). For single-nucleus suspension preparation, collected tissues were washed with 1× PBS (Servicebio, G4202-500ML), rapidly frozen, and stored in liquid nitrogen. Nuclei were isolated using a mechanical extraction approach.

Briefly, tissues were transferred into a 2-mL Dounce homogenizer and thawed in homogenization buffer (20 mM Tris, pH 8.0, 500 mM sucrose, 0.1% NP-40, 0.2 U/μL RNase inhibitor (Beyotime, R0102-2kU), 1× protease inhibitor cocktail (Beyotime, P1005), 1% bovine serum albumin (BSA) (Servicebio, GC305010-100g), and 0.1 mM DTT). The samples were homogenized with pestle A for 10 strokes, passed through a 70-μm cell strainer, then further homogenized with pestle B for 10 strokes and filtered through a 30-μm cell strainer. The homogenate was centrifuged at 500 × g for 5 min at 4°C to pellet nuclei, which were subsequently resuspended in blocking buffer (1% BSA, 0.2 U/μL RNase inhibitor in 1× PBS). After a second centrifugation under the same conditions, the nuclei pellet was resuspended in Cell Resuspension Buffer (MGI).

ScRNA-seq/snRNA-seq libraries were prepared using the DNBelab C4 Series Single-Cell Library Prep Set (MGI, 1000021082) as previously described^108^. In brief, single-nucleus/cell suspensions were used for droplet generation, emulsion breakage, bead collection, reverse transcription and cDNA amplification to generate barcoded libraries. Indexed libraries were constructed according to the manufacturer’s protocol. Concentrations were measured with a Qubit ssDNA Assay Kit (Thermo Fisher Scientific, Q10212). Libraries were sequenced on DNBSEQ-T7 platform at the China National GeneBank (Shenzhen, China).

### Processing of scRNA/snRNA-seq data analysis

Raw reads were obtained in FASTQ format from C4 sequencing platform and were processed using an open-source pipeline (https://github.com/MGI-tech-bioinformatics/DNBelab_C_Series_scRNA-analysis-software) to obtain expression matrices. Briefly, all samples were performed sample de-multiplexing, barcode processing, and single-cell 3’ unique molecular identifier (UMI) counting with default parameters. Processed reads were then aligned to reference genomes using STAR (version 2.5.1b) ^109^. *Petaurus breviceps*, *Homo sapiens*, *Mus musculus*, *Tupaia belangeri chinensis*, *Canis lupus familiaris*, *Ovis aries*, *Capra hircus* and *Sus scrofa domestica* were mapped to their respective NCBI genome annotations (PetGlider_PUasm1.0 (GCF_028583685.1), GRCh38.p14 (GCF_000001405.40), GRCm39 (GCF_000001635.27), TupChi_1.0 (GCF_000334495.1), ROS_Cfam_1.0 (GCF_014441545.1), ARS_UI_Ramb_v3.0 (GCF_016772045.2), ARS1.2 (GCF_001704415.2) and Sscrofa11.1 (GCF_000003025.6), respectively). Valid cells were then automatically identified based on the UMI number distribution of each cell by using the “barcodeRanks()” function of the DropletUtils tool to remove background beads and the beads that had UMI counts less than the threshold value. Finally, we used PISA to calculate the gene expression of cells and create a matrix for each library. The expression matrices of human placenta and decidua at 4-13 weeks, cynomolgus macaque samples generated from stages E29 and E45, placenta and decidua from early recurrent miscarriage patients, gestational diabetes and public cancer (i.e., lung adenocarcinoma, colorectal cancer, gastric cancer, breast cancer, clear cell renal cell carcinoma, pancreatic cancer, leiomyoma, prostate cancer, ovarian cancer, hepatocellular carcinoma, high-grade squamous intraepithelial lesion, basal cell carcinoma and glioma) scRNA data were obtained from E-MTAB-6701 at ArrayExpress^104^, GSE180637^105^, GSE174399^83^, GSE173193^106^, GSE123904^110^, E-MTAB-8410^111^, GSE134520^112^, NGDC: PRJCA008495^113^, GSE210038^114^, GSE212966^115^, GSE162122^116^, GSE181294^117^, E-MTAB-8107^118^, GSE242889^119^, GSE208653^120^, E-MTAB-13085^121^, and GSE249263^122^.

For each sample, we used CreateSeuratObject function from Seurat (version 5.1.0)^123^ R package to generate a Seurat object from the expression matrix. Biological replicates were integrated using the merge function in R. Cells with fewer than 200 detected genes and for which the total mitochondrial gene expression exceed 5% were removed. DoubletFinder (version 2.0.5) was used to detect doublets cells with default parameters^124^. Data normalization was conducted using the NormalizeData function, and the top 2,000 most variable genes were identified with the FindVariableFeatures function. Data were then scaled and batch effects were regressed out using ScaleData function. Principal component analysis (PCA) was performed on the variable features and the batch correction was performed by RunHarmony function in harmony (version 1.2.0)^125^. Subsequently, the batch-corrected embeddings were used to compute a UMAP for visualization by RunUMAP function. Cell–cell relationships were constructed with a shared nearest neighbor (SNN) graph based on the first 10 Harmony dimensions by FindNeighbors function, and clusters were finally identified using the FindClusters function at a resolution of 0.6. For subclustering of a specific cell population, cells belonging to this major cluster were re-extracted, and clustering was reperformed using the FindClusters function in Seurat with the resolution parameter set to 1. After obtaining cell clusters, marker gene identification for each cluster was performed using the FindAllMarkers function with default parameters.

For cell type annotation, we identified the main cell-type populations using known marker genes mostly from human and mouse^104,126–128^ and their respective 1:1 orthologs in the other species based on OrthoFinder (version 2.5.5) (Fig. 1b and Extended Data Fig. 3a)^129^. In detail, for stromal cells, we used e.g., *LUM*, *DCN*, and *COL5A2* to mark. After subclustering, we distinguished fetal/maternal stromal cells based on sample position, and also the marker genes *TCF21* and *WT1*, respectively. We then annotated the cell subtypes of maternal stromal cells and clustered them into two major groups, dS1 (*IGFBP6*, *MFAP5*, *SCARA5*, and *WT1*) and dS2 (*APOLD1*, *GPR4*, *KCNJ8*, *ACTA2*, and *TAGLN*). For immune cells, e.g., *TYROBP*, *RGS1*, and *C1QC* were used to mark them. After subclustering, *LYVE1* marks hofbauer (HB) cells, *NKG7* mark decidual natural killer (dNK) cells, *IL10* and *OSM* mark decidual macrophages (dM). For trophoblasts, *TFAP2A*, *KRT19* and *CLDN7* were used to mark. For endothelial cells, we used e.g., *PLVAP, EMCN* and *KDR* to mark them and distinguished fetal/maternal side based on sample position and also the marker genes *MEOX2* and *VWF*, respectively. For epithelial cells, *EHF*, *DCDC2*, and *SYT13* were used to mark them. All detailed cell type descriptions are provided in Supplementary Table 2. In addition, we calculated the Z-score of the marker genes (i.e., top 20 genes of log_2_FC for each cluster) for human annotated cells and those unclassified cells in other species. Specifically, the average expression values were obtained using the AverageExpression function in Seurat, followed by z-score normalization across genes. The normalized expression profiles were integrated across species based on one-to-one orthologs, and then a heatmap was generated to visualize their clustering relationships. Cell types were further annotated by referencing the corresponding human cell populations.

In addition, to systematically identify macrophage subtypes resembling M1 or M2 polarization states of HB/dM, we implemented a quantitative scoring framework integrating curated marker gene sets with differential expression profiles across clusters. Briefly, two reference gene lists were constructed to represent canonical M1-like and M2-like macrophage signatures, respectively (https://www.biocompare.com/Editorial-Articles/566347-A-Guide-to-Macrophage-Markers/; Supplementary Table 27). Each list was compiled from orthologous gene mappings between species, where each entry linked an ortholog group to its corresponding gene symbol. For each cluster in the single-cell RNA-seq dataset, marker gene statistics were obtained from the differential expression table, which included average log₂ fold change (avg_log2FC), expression percentages in the cluster (pct.1) and in other clusters (pct.2), and adjusted p-values. To estimate the relative contribution of each macrophage subtype, we calculated a contribution value for each gene according to the formula:

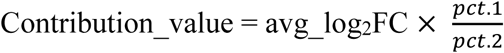

This value reflects both the magnitude and specificity of expression enrichment. For each cluster, total M1 and M2 contribution scores were computed as the sum of individual contribution values of genes belonging to the M1 or M2 reference sets, respectively. The relative contributions were then normalized to obtain subtype-specific proportions:

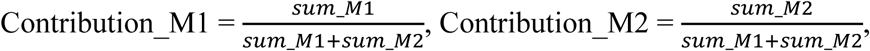

Clusters were classified as M1-like or M2-like when their corresponding subtype contribution exceeded 0.6. Clusters not meeting either threshold were designated as unclassified.

### Gene expression trees construction across mammals

Pseudo-bulk samples were generated from groups of cells using the AverageExpression function of the Seurat R package. For each cell type conserved across species, we constructed gene expression trees following the approach of Brawand et al. (2011)^130^. Pairwise expression distance matrices were first computed between the corresponding pseudo-bulk samples, where the distance was defined as 1 − ρ, with ρ representing Spearman’s correlation coefficient calculated using the cor function of the R stats package. Neighbour-joining trees were then inferred using the ape R package (v5.8)^131^. The robustness of branching patterns was evaluated through bootstrap analyses, in which 10,573 one-to-one orthologs were randomly sampled with replacement 1,000 times. Bootstrap values represent the proportion of replicate trees that support the branching patterns observed in the majority-rule consensus tree. To calculate the total tree length, intra-species variability among individuals was excluded, following the approach of Murat et al. (2023)^132^. The total tree length for each species phylogeny was scaled to account for differences in the number of taxa. After excluding intra-species variability, the standardized tree length was calculated as the total branch length of the pruned tree divided by (*n* − 1), where *n* is the number of taxa included in the tree. This normalization ensures that tree length comparisons are not biased by tree size.

### Integration of scRNA-seq data across species

To integrate maternal-fetal interface cells across multiple mammalian species, we applied SATURN^29^, which couples the biological properties of protein language models (pLMs) with single-cell RNA expression. To generate protein embedding for each species, cleaned protein FASTA files were embedded using the ESM-1b transformer model (esm2_t48_15B_UR50D)^133^ through ‘extract.py’ in esm (https://github.com/facebookresearch/esm). Protein embeddings were mapped back to gene symbols, and converted into gene-level embeddings using ‘convert_protein_embeddings_to_gene_embeddings.py’ in SATURN (https://github.com/snap-stanford/SATURN/blob/main/protein_embeddings). ScRNA-seq data from Seurat objects onto h5ad format using the SaveH5Seurat and Convert functions within the SeuratDisk package in R. Counts were then processed into AnnData objects and paired with the corresponding gene-level embeddings. Cell type annotations were provided as label metadata. To balance dataset sizes, we randomly downsampled the dataset using the ‘preprare_h5ad.clean.py’ script. Finally, SATURN was trained using ‘train-saturn.py’ based on the following parameters: pretrain_batch_size= 10240, batch_size=10240, --num_macrogenes=6, model_dim=256, --polling_freq=201, --epochs=50, --hv_genes=0, --hv_span=1, --pretrain_epochs=200, --pe_sim_penalty=1.0, --l1_penalty=0, --centroid_score_func=default. For larger datasets, we scaled up training with GPU support and adjusted max_split_size_mb to optimize CUDA memory allocation. The integrated embeddings were examined by PCA and UMAP to assess overlap of cell populations across species.

SAMap (version 1.0.16) was used to calculate gene homology-aware similarity scores between cells of different species^134^. For each species, processed single-cell RNA-seq datasets were stored in .h5ad format, and cell type annotations were included as metadata columns. Protein homology relationships between the pairwise species were obtained from pre-computed BLAST (version 2.16.0) results and supplied to SAMap as external gene-to-gene correspondence maps. The mapping procedure was run with neighborhood graphs constructed from both species (neigh_from_keys = True). The resulting SAMap object, containing the cross-species alignment, was stored in a serialized file (samap.pkl) for downstream analysis. The SAMap objects generated from pairwise species comparisons were subsequently analyzed to quantify and visualize cross-species cell type correspondences. For each SAMap run, the serialized result object (samap.pkl) was reloaded in Python. Mapping scores between cell types across species were computed using the get_mapping_scores() function from the SAMap analysis module. To visualize the degree of alignment, we generated Sankey plots using the sankey_plot() function in SAMap, with an alignment threshold (align_thr) of 0.3 for trophoblasts, immune cells and 0.2 for stromal cells.

### Differential expression analysis for homologous cell subtype across different placental type, pregnancy diseases and cancers

We identified homologous cell subtype of VCT, SCT, EVT, HB, fS, dS1, dS2, dM, dNK, dNKT, Tcell across species based on the gene markers and the integrated results of SATURN and Samap with the alignment threshold (align_thr) of 0.3 for trophoblasts, immune cells and 0.2 for stromal cells. For each homologous cell subtype, cross-species single-cell transcriptomic integration and differential expression analysis were performed using Seurat (version 5.1.0) and Harmony (version 1.2.0). To ensure comparability, the intersection between expressed genes and the orthologous gene list was determined based on 11,638 one-to-one orthologous genes, and only genes present in all eight species were retained for subsequent analyses. The individual objects were merged into a single Seurat object, followed by standard preprocessing steps: log-normalization, identification of 2,000 variable features, data scaling, and principal component analysis (PCA). Batch effects across species were corrected using Harmony, with species designated as the grouping variable. Species were further grouped three placental types: hemochorial (human, macaque, mouse), endotheliochorial (dog, tree shrew), and epitheliochorial (sheep, goat, pig) based on the layers of maternal and fetal tissues separating the maternal blood from the fetal blood^135^. Differential gene expression analyses were performed for each placental group against the others using the Wilcoxon rank-sum test, as implemented in the FindMarkers function of Seurat. Differentially expressed genes (DEGs) were defined as those detected in ≥10% of cells in the given group (min.pct = 0.1) with log fold-change > 0.75 and *P*_adj_ < 0.05. For pregnancy disorder datasets, we applied the same analytical framework as described above to perform cell type–specific differential expression analysis between disease and control groups.

For tumor datasets, we performed differential expression analysis by comparing tumor with adjacent non-tumor for each cell type. Specifically, tumor cells (i.e., mostly epithelial cancer cells) were analogous to extravillous trophoblasts (EVTs) from hemochorial placentas due to their invasive properties, and differential expression was assessed against corresponding cell types from adjacent non-tumor tissues (Supplementary Table 18). Similarly, cancer-associated fibroblasts were analogous to decidual stromal (dS) cells from the hemochorial placenta, with normal fibroblasts serving as controls. In addition, immune subsets within tumors, including macrophages, natural killer (NK) cells, natural killer T (NKT) cells and T cells, were analyzed in the same framework by comparing tumor-infiltrating immune cells to their counterparts in adjacent non-tumor tissues. DEGs were identified using the FindMarkers function in Seurat, applying thresholds of ≥10% of cells in the given group (min.pct = 0.1), Padj < 0.05 and log fold-change > 0.75. Pathway enrichment analysis for DEGs in this study was performed using Metascape (https://metascape.org).^136^

Finally, we performed permutation test to check if the overlaps between cancer and hemochorial up-regulated genes for each cell type were significantly different from those expected at random. To this end, we used sample function in R to generate simulated gene sets in the human genome by randomly selecting genes of equal number to the observed up-regulated genes in hemochorial, and we replicated this process 10,000 times. We compared the number of overlaps between the observed hemochorial up-regulated and cancer up-regulated genes with the overlaps between the simulated hemochorial up-regulated and cancer up-regulated genes, and calculated the statistical significance of *P* values (i.e., the probability that a higher number of overlaps would be observed by chance).

### Identification and verification of core genes for placenta invasiveness

We further calculated standardized Z-score and normalized counts per million (CPM) of DEGs for EVT and dS pseuodo-bulk expression values across species. To calculate Z-score in each species, we firstly computed the average expression level of each gene across cells grouped by cell type using the AverageExpression function in Seurat. The resulting gene-by-cell-type expression matrix was then scaled on a per-gene basis by subtracting the mean expression and dividing by the standard deviation of that gene across all cell types. This transformation yielded a z-score for each gene in each cell type, reflecting its relative up- or down-regulation compared to the gene’s average expression profile. To calculate CPM in each species, Seurat objects were converted into a SingleCellExperiment object, and cell-level metadata were organized using the prepSCE function in the muscat package^137^. Gene-level count data were aggregated at the cell type level using aggregateData with the sum function. For each cell type, raw counts were then normalized to CPM by dividing gene counts by the total number of counts per column and multiplying by one million. Z-score/CPM of DEGs across eight species were finally integrated into one matrix.

To identify the potential major genes responsible for the placenta invasiveness, we applied additional filters based on both CPM and Z-score values of DEGs. For a given target group, we required that candidate upregulated genes satisfy the following criteria: (*i*) within the target group, the minimum CPM/Z-score value across all species must be greater than the maximum CPM/Z-score value observed in the remaining species; and (*ii*) for Z-scores, all species within the target group must have values greater than 0.5. Genes fulfilling these thresholds were retained as robust group-specific DEGs. For the filtering DEGs of EVT based on Z-scores, a protein-protein interaction network was constructed with combine scores which were retrieved from the STRING version 12.0 database (https://string-db.org/)^138^. Mcode^139^ plug-ins of Cytoscape^140^ were used to analyze the key gene clusters and hub genes.

### Cell-cell communication analysis

We performed cell–cell communication analysis across nine species (sugar glider, human, cynomolgus macaque, mouse, tree shrew, dog, sheep, goat, and pig) using CellPhoneDB v5.0.0^141^. To enable cross-species comparison of intercellular communication networks, we constructed a unified gene expression matrix based on human genes as the reference. OrthoFinder (version 2.5.5) was employed to infer paralogs of the species. For each non-human species, we obtained cell-type–specific gene expression count matrices by LayerData function in Seurat package and summed the counts of all paralogs corresponding to a single-copy human gene within the same orthogroup. The resulting value was then assigned to the respective human gene ID. This procedure yielded a “human-equivalent” gene expression matrix for each species, which was used exclusively for downstream analyses of ligand–receptor interactions. The statistical analysis was performed using CellPhoneDB to identify significant ligand–receptor interactions. Only receptors and ligands expressed in more than 10% of the cells within a given cluster and with a p-value < 0.05 were retained. Interacting pairs that were consistently detected across all species with hemochorial, endotheliochorial, or epitheliochorial placentation, respectively, were preserved. Moreover, lineage-specific interactions were defined when both the receptor and ligand were encoded by genes upregulated in the corresponding cell populations of hemochorial, endotheliochorial, or epitheliochorial species.

### Statistical analysis of escalation

To investigate the co-evolution of fetal ligands and maternal receptors, we use a reduced copy (one to one) of the CellPhoneDB ligand–receptor database (Supplementary Table 28)^25^ to generate the candidate testing pairs by cutting the interactions to only secreted ligands and their cognate ligand-binding subunit genes, excluding co-receptors. For each ligand-receptor pair, we first identified representative expression values within each species. Specifically, the fetal ligand expression was defined as the maximum average expression across all fetal cell types, while maternal receptor expression was the maximum average expression across all maternal cell types. These expression values were then standardized using a z-score transformation of log_2_-transformed Transcripts Per Million (TPM) values (zTPM) to mitigate biases from differing dynamic ranges of gene expression between species.

We tested co-evolution for each placental type, with Petaurus breviceps serving as the outgroup. First, a standard linear regression (receptor_zTPM ∼ ligand_zTPM) was performed on the cross-species expression values to determine the direct regression slope, reflecting the magnitude and direction of the association before phylogenetic correction. Second, to account for shared evolutionary history, we used Phylogenetically Independent Contrasts (PICs). PICs were calculated for the zTPM values of each ligand and its corresponding receptor using the pic() function in the R package ‘ape’, and a time-calibrated phylogeny of our nine species form TimeTree5^32^. The resulting *P* value, representing the statistical significance of the co-evolutionary relationship, was -log10 transformed and plotted on the y-axis. Ligand-receptor pairs with a PIC *P* value < 0.05 and a significant positive (slope > 1) or negative (slope < −1) regression slope were highlighted as instances of symmetric or asymmetric escalation, respectively.

### *In vitro* invasion assays

Spheroid invasion assay was performed as previously described^37,65^. Fibroblasts (ESFs or BJ) were seeded on nanopatterned substrate and cultured until an aligned monolayer was established. ESFs were decidualized in vitro in DMEM/F12 supplemented with 2% FBS, 0.5mM 8-Bromo-cAMP, and 1 µM medroxyprogesterone 17-acetate for 4 days. Extravillous trophoblast (HTR8) and colon cancer (HCT 116) were labelled by lentiviral transduction with H2B-tdTomato (SignaGen). For gene perturbation, siRNAs (including the NC-1 scrambled negative control) were purchased from IDT. Cells were seeded at 50% confluence, and gene knockdown was performed using Lipofectamine RNAiMAX (Thermo Fisher) according to the manufacturer’s instructions. EVT and cancer spheroids were generated in ultra-low attachment round bottom microplate (Corning) and plated onto the pre-aligned fibroblast monolayer for invasion. Time-lapse images were acquired every 2 or 4 h for 24 or 48 h. To quantify invasion, spheroid boundaries at 0 h and 24/48 h were manually outlined, fit to an ellipse, and the major axis length were extracted in Fiji/ImageJ. To account for variation in initial spheroid size, the 24 or 48 h major axis length was normalized to corresponding 0 h value. Statistical analysis was performed using Prism.

### *In utero* placental injection of siRNA encoding *APOE*

Pregnant C57BL/6J mice at embryonic day 12.5 (E12.5) were anesthetized with isoflurane delivered via an anesthetic vaporizer for a maximum duration of 40 min (RWD Life Science Co., Ltd., Shenzhen, Guangdong, China). A midline laparotomy was performed to expose the uterine horns, as previously described^142^. Throughout the surgical procedure, the abdominal cavity, particularly the exposed uterine horns, was continuously moistened with warmed physiological saline, and maternal body temperature was maintained using a heating plate. *In vivo* injections of NaCl, scrambled siRNA or *APOE*-targeting siRNA were performed using glass capillary micropipettes (50 μm inner diameter, 100 μm outer diameter; Lqwlbio, 1.0 × 0.58–100) pulled with a P-2000 laser-based micropipette puller (Sutter Instrument Company, Novato, CA). The micropipette tips were further refined using a PG-22C micropipette grinder (Lqwlbio), and injections were carried out using a FemtoJet 4i microinjector (Eppendorf). Both *APOE*-targeting siRNAs and scrambled control siRNAs were designed and synthesized by RiboBio. The sequences of the *APOE* siRNA were as follows: sense, 5′-GCA ACA ACA UCC AUA UCC A dTdT-3′; antisense, 5′-UGG AUA UGG AUG UUG UUG C dTdT-3′^143^. Scrambled siRNA–treated placentas (*n* = 4) served as negative controls (NC; siBDMV002, RiboBio), while normal saline–treated placentas (*n* = 4) were used as vehicle controls (Innochem, B46918). All *in vivo* siRNAs were chemically modified with 2′-O-methyl ribose substitutions and conjugated to 5′ cholesterol to enhance stability and cellular uptake. To visualize and track the injection process, Cy5 fluorophores were conjugated to the 5′ end of the siRNAs. Each placenta was injected to a depth of approximately 0.5 mm with 4–5 μL of siRNA solution, corresponding to a dose of 5 nmol per placenta. To maximize embryo survival, no more than two placentas per dam were injected. Following injection, the uterine horns were gently returned to the abdominal cavity, and the abdominal wall and skin were closed with sterile absorbable PGA suture (Jinhuan, R516) and silk sutures (WEGO, S43183). At E15.5, only viable fetuses were harvested for gross morphological assessment, quantitative analysis, qPCR, western blotting, immunofluorescence, and hematoxylin and eosin (H&E) staining. Placental tissues were dissected using RNase-free forceps and blades. Each placenta was bisected along the midline: one half was fixed in 4% paraformaldehyde at 4 °C, while the remaining half was further divided into two quarters and snap-frozen at −80 °C in separate tubes, one of which contained RNA preservation reagent for downstream RNA analysis.

### Gene expression and western blot analyses

The total RNA was extracted automatically by HollySys AE2100 using Magbead Tissue RNA kit (with DNase I) (CWBIO, CW3711S) or manually using FastPure Complex Tissue/Cell Total RNA Isolation Kit (Vazyme, RC113-01). The RNA concentrations were exanimated by NanoDrop. The first-strand cDNA was synthesized using the HiScript IV All-in-One Ultra RT SuperMix for qPCR (Vazyme, R433-01) Following the manufacturer’s instruction, 500 to 1000ng of RNA was reverse transcribed as the template for RT-PCR in 20μl volume (including 5μl 4x All-in-One Ultra qRT SuperMix, specifical volume of template RNA (calculated by RNA concentration) and nuclease-free water adding to 20 μl) and a thermocycling condition at50℃ for 5 min, and 85℃ for 5 sec. Subsequently, the qPCR with SupRealQ Purple Universal SYBR qPCR Master Mix (U+) (Vazyme, Q412-02) was performed on the qTOWER^3^G (Analytik-Jena, 844-00554-x) using the first-strand cDNA. The qPCR reactions and conditions were set as those described above. Each qPCR was run three times for one sample as technical replicates. Based on the qPCR results, the expression level was calculated using the 2^−ΔΔCt^ method^144^ and normalized according to the internal control *RPS20* gene. All primers for *APOE, MMP2, MMP9, RPS20, CD86*, *CCL2*, *NOS2*, *IL6*, *MRC1*, *C1QA*, *C1QB*, *C1QC*, *CD163*, *ARG1*, *CSF1R*, and *IL1B* are provided in Supplementary Table 29.

Placental protein extracts were prepared from normal saline–treated, scrambled siRNA–treated, and *APOE* siRNA–treated mice. Placental tissues were extracted using BBproExtra® Total Protein Extraction Kit (BestBio, BB-3101) on ice, and protein concentrations were determined using standard BCA methods. Equal amounts of protein (50μg) from placental samples were mixed with diluted SDS-PAGE Protein Sample Loading Buffer (5×) (Biosharp, BL502A), boiled at 95℃ for 5 min, and loaded onto a 12% Super-PAGE™Bis-Tris Gel (Epizyme, LK305). Following electrophoretic separation, proteins were transferred onto 22 μm polyvinylidene fluoride (PVDF) membranes. Membranes were blocked in blocking solution (5% skimmed milk in TBST) at room temperature for 1 h and then incubated overnight at 4 °C with primary antibodies (Apoe Recombinant monoclonal antibody (1:500, Proteintech, 83728-4-RR), HRP-conjugated Beta Actin Monoclonal antibody (Proteintech, HRP-60008)). After washing 3 times with TBST, membranes were incubated with horseradish peroxidase–conjugated secondary antibodies including HRP-Labeled Goat anti-Rabbit IgG (H+L) (Beyotime, A0208) and HRP-Labeled Goat anti-Mouse IgG (H+L) (Beyotime, A0216). Immunoreactive bands were detected using an enhanced chemiluminescence (ECL Plus) detection system (Beyotime, P0018S). Band intensities were quantified using a blot analysis system (Bio-Rad Laboratories, Marne-la-Coquette, France), and β-actin was used as a loading control. Prestained Multicolor Protein Marker (CWBIO, CW2841) were used to determine protein sizes. For densitometric analysis, band intensities were quantified using ImageJ software^145^. Protein expression levels were normalized to β-actin and expressed as relative values compared with control samples.

### Hematoxylin and eosin (H&E) and immunofluorescence

Tissue samples were first fixed and then rinsed three times in phosphate-buffered saline (PBS; pH 7.4) for 10 minutes each to remove any residual fixative. They were dehydrated through a graded ethanol series, starting with 70% ethanol for 1.5–2 hours, followed by 80% ethanol for 1.5 hours, two changes of 95% ethanol (1.5–2 hours each), and two changes of absolute ethanol (2 hours each). After dehydration, the tissues were cleared in xylene (two changes, 1 hour each) and infiltrated with molten paraffin wax (three changes, 1 hour each) at 60°C, before being oriented and embedded in fresh paraffin blocks. Paraffin-embedded tissue blocks were sectioned at a thickness of 4–5 µm using a rotary microtome (LEICA RM2235). The sections were then floated in a warm water bath (42–45°C), mounted onto positively charged glass slides, and dried using a Slide Dryer (KEDEE Co.,LTD. KD-HΙ Jinhua, Zhejiang, China). For Hematoxylin and Eosin (H&E) staining, slides were deparaffinized in xylene (two changes, 10 minutes each) and rehydrated through an ethanol series (100%, 95%, 80%, 70%, 5 minutes each), followed by a rinse in distilled water. Nuclei were stained with Mayer’s hematoxylin for 3–5 minutes and then washed briefly in running tap water. Cytoplasmic counterstaining was performed using eosin Y solution for 1–2 minutes. After dehydration through an ascending ethanol series (70%, 80%, 95%, 100%, each for 30 seconds) and clearing in xylene (two changes, 2 minutes each), the slides were coverslipped with a synthetic mounting medium. Thickness measurements were performed on sections obtained from the central one-third region of the placenta. The thickness of the decidual layer, junctional zone, and labyrinth zone was quantified using CaseViewer software. For each sample, three consecutive sections were analyzed, and for each section, measurements were repeated at least three times. The average values were used for subsequent statistical analysis.

For immunofluorescence (IF) staining, sections were deparaffinized and rehydrated as described above. Antigen retrieval was carried out by heating the slides in Citrate Antigen Retrieval Solution (pH 6.0, Beyotime, P0081) in a 100°C water bath for 20 minutes, followed by cooling to room temperature. Permeabilization was performed using 0.2% Triton X-100 (Sangon Biotech, A110694-0100) in PBS for 10 minutes at room temperature, and non-specific binding sites were blocked by incubation with 5% bovine serum albumin (BSA, Servicebio, GC305010-100g) in PBS for 1 hour at room temperature. Sections were then incubated overnight at 4°C with primary antibodies, including Anti-Cytokeratin 8 (CK8; 1:500, Abcam, ab53280), and APOE Recombinant Monoclonal Antibody (1:500, Proteintech, 83728-4-RR), all diluted in blocking solution. The following day, slides were washed three times with PBS (5 minutes each) and incubated with species-specific fluorophore-conjugated secondary antibodies (Goat Anti-Rabbit IgG H&L, Alexa Fluor 594; 1:1000, Abcam, ab150080) for 1 hour at room temperature in the dark. After washing three times with PBS, autofluorescence was quenched using the TrueVIEW Autofluorescence Quenching Kit (Vectorlabs, SP-8400-15) for 5 minutes at room temperature. Nuclei were counterstained with 4′,6-diamidino-2-phenylindole (DAPI; 1 µg/mL) for 5 minutes, and the slides were mounted with an anti-fade mounting medium. Finally, stained sections were examined and imaged using a slide scanner (3Dhistech Pannoramic MIDI II and OLYMPUS VS120) with appropriate filter sets for visualization of specific fluorophores, and images were captured using dedicated software. Trophoblast invasion depth was measured as the distance the farthest CK8^+^ trophoblast was found in the decidua relative to the junctional zone. For each sample, measurements were repeated at least three times. The average values were used for subsequent statistical analysis.

### RNA-Seq analysis

Placental tissues were collected from mice injected with scrambled siRNA (*n* = 7) or *APOE* siRNA (*n* = 6) for RNA-seq analysis. Total RNA was extracted using the Total RNA Extractor (Trizol) Kit (B511311, Sangon, China) according to the manufacturer’s instructions and treated with RNase-free DNase I to eliminate genomic DNA contamination. RNA integrity was assessed on a 1.0% agarose gel. High-quality RNA samples were then submitted to Sangon Biotech (Shanghai) Co., Ltd. for library preparation and sequencing. Messenger RNA libraries were constructed using the VAHTS™ mRNA-seq V2 Library Prep Kit for Illumina® following the manufacturer’s protocol and sequenced on a NovaSeq platform (Illumina, San Diego, CA) to generate 150-bp paired-end reads. Raw reads were filtered using fastp (v1.3.2)^146^, aligned to the mouse reference genome (GCF_000001635.27_GRCm39) with HISAT2 (v2.2.1)^147^, converted to sorted BAM files using SAMtools (v1.21)^148^, and quantified at the gene level using featureCounts (v2.1.1).^149^ Gene expression levels were normalized and reported as fragments per kilobase of exon per million mapped fragments (FPKM). Pearson correlation coefficients among samples were calculated using the cor() function in R (v4.3.1). Differentially expressed genes (DEGs) between the scrambled and ApoesiRNA groups were identified through DESeq2 (v1.42.1)^150^. Genes with a log₂(fold change) ≥ 2 and adjusted *P* ≤ 0.05 were defined as up-regulated, whereas those with a log₂(fold change) ≤ −1 and adjusted *P* ≤ 0.05 were defined as down-regulated in the *Apoe* siRNA group. Pathway enrichment analysis of DEGs was performed using Metascape.

## Data availability

The raw and processed data generated in this study have been deposited in the NCBI database under the accession code PRJNA1434137 (https://dataview.ncbi.nlm.nih.gov/object/PRJNA1434137?reviewer=b47a6nvsef4ida7ccf2954f2 2u). The expression matrices of human placenta and decidua at 4-13 weeks, cynomolgus macaque samples generated from stages E29 and E45, placenta and decidua from early recurrent miscarriage patients, gestational diabetes and public cancer (i.e., lung adenocarcinoma, colorectal cancer, gastric cancer, breast cancer, clear cell renal cell carcinoma, pancreatic cancer, leiomyoma, prostate cancer, ovarian cancer, hepatocellular carcinoma, high-grade squamous intraepithelial lesion, basal cell carcinoma and glioma) scRNA data were obtained from E-MTAB-6701, E-MTAB-8107, E-MTAB-8410, and E-MTAB-13085 at ArrayExpress, GSE180637, GSE174399, GSE173193, GSE123904, GSE134520, NGDC: PRJCA008495, GSE210038, GSE212966, GSE162122, GSE181294, GSE242889, GSE208653, and GSE249263 at Gene Expression Omnibus in the NCBI database.

## Code availability

All code generated in the study, including analysis parameters, are available at https://github.com/MsLeexinxin/Placenta_evolution.

## Acknowledgements

We thank Jacobus J. Boomsma and Jiaming Chen for their suggestions and assistance in revising the article. We thank Zheyi Ni and Xiao Sun for technical guidance in the experiments. We thank Baohua Chen, Huimin Cai, Sheng Qian, Guangji Chen and Dongya Wu for their advice and guidance on data analysis. We thank Danyang Sun for the experimental help. We thank Dan Yang from the Core Facilities, Zhejiang University School of Medicine for their technical support. And we also acknowledge the Information Technology Center of Zhejiang University and China Mobile Zhejiang Co., Ltd (Hangzhou Branch) for providing.

G.Z. is supported by the National Key Research and Development Program of China (grant no. 2024YFA1802500), Basic Research Center Program (grant no. 32388102), G.Z. and D.Z. are both supported by Fundamental and Interdisciplinary Disciplines Breakthrough Plan of the Ministry of Education of China (grant no. JYB2025XDXM508). G.Z. is also supported by the New Cornerstone Science Foundation through the XPLORER PRIZE and the K.C. Wong Education Foundation. D.Z. is supported by the National Key Research and Development Program of China (grant no. 2025YFC2708200), and National Natural Science Foundation of China (grant no. 82394424). K. is supported by the National Institute of Child Health and Development (grant no. R01HD112424). X.L. is supported by the National Natural Science Foundation of China (grant no. 32300344), the Postdoctoral Fellowship Program and China Postdoctoral Science Foundation under grant nos. 2023M743037 and GZB20230649.

## Author contributions

G. Z., D. Z. and X. L. conceived this work. X. L., R. C., Y. L., D. Z., J. S., L. J., D. Y., Y. S., Y-Q. Y. and L. S. participated in sampling collection, extraction, library preparation and sequencing. X. L., Y. Z., X.B. and K. S. analyzed the data. X. L., G. Z., K., W. D., Y. Z., D. Y., Y. S., Y-Q. Y., Y-J. Y and Y. H. designed and performed the experiments. J. L. provided technical support. X. L., G. Z., K., W. D., Y. Z. and Y. S. wrote the initial draft of the manuscript with input from all co-authors. G. Z. and D. Z. supervised this project.

## Competing interests

The authors declare no competing interests.

## Additional information

## Supplementary information

The online version contains supplementary material available at xxx.

---

**Extended Data Fig. 1.**
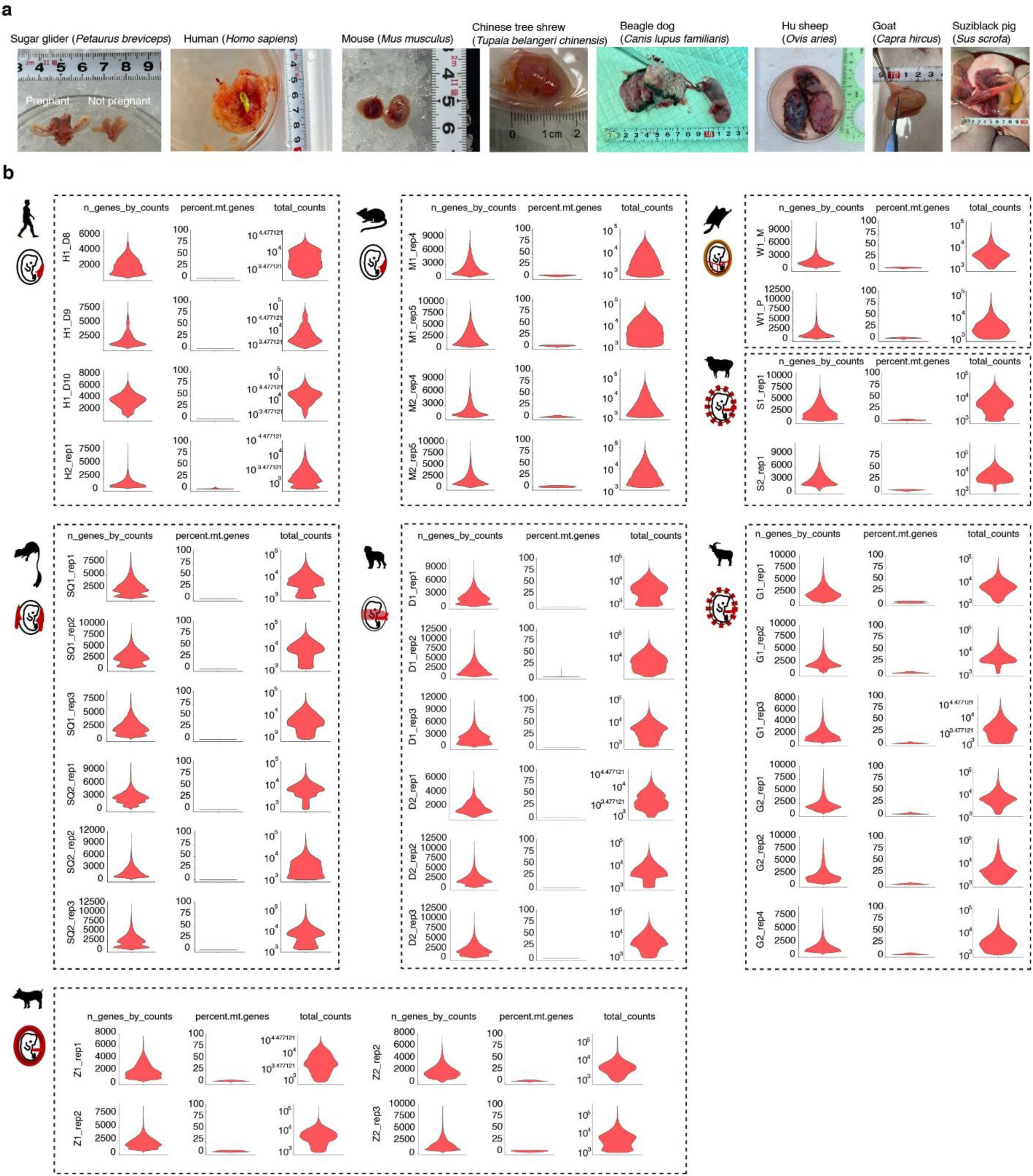
Sampling and single-cell RNA-seq data quality assessment of the maternal-fetal interface excluding macaque samples, which were quality-controlled by the original studies. **a,** Placental morphology of eight species at the time of sample collection. **b,** Violin plots of quality control metrics across all biological replicates for species included in quality control analysis. n_genes_by_counts: number of unique genes detected per cell; total_counts: total unique mRNA counts per cell; pct.counts.mt: percentage of mitochondrial gene counts.

**Extended Data Fig. 2.**
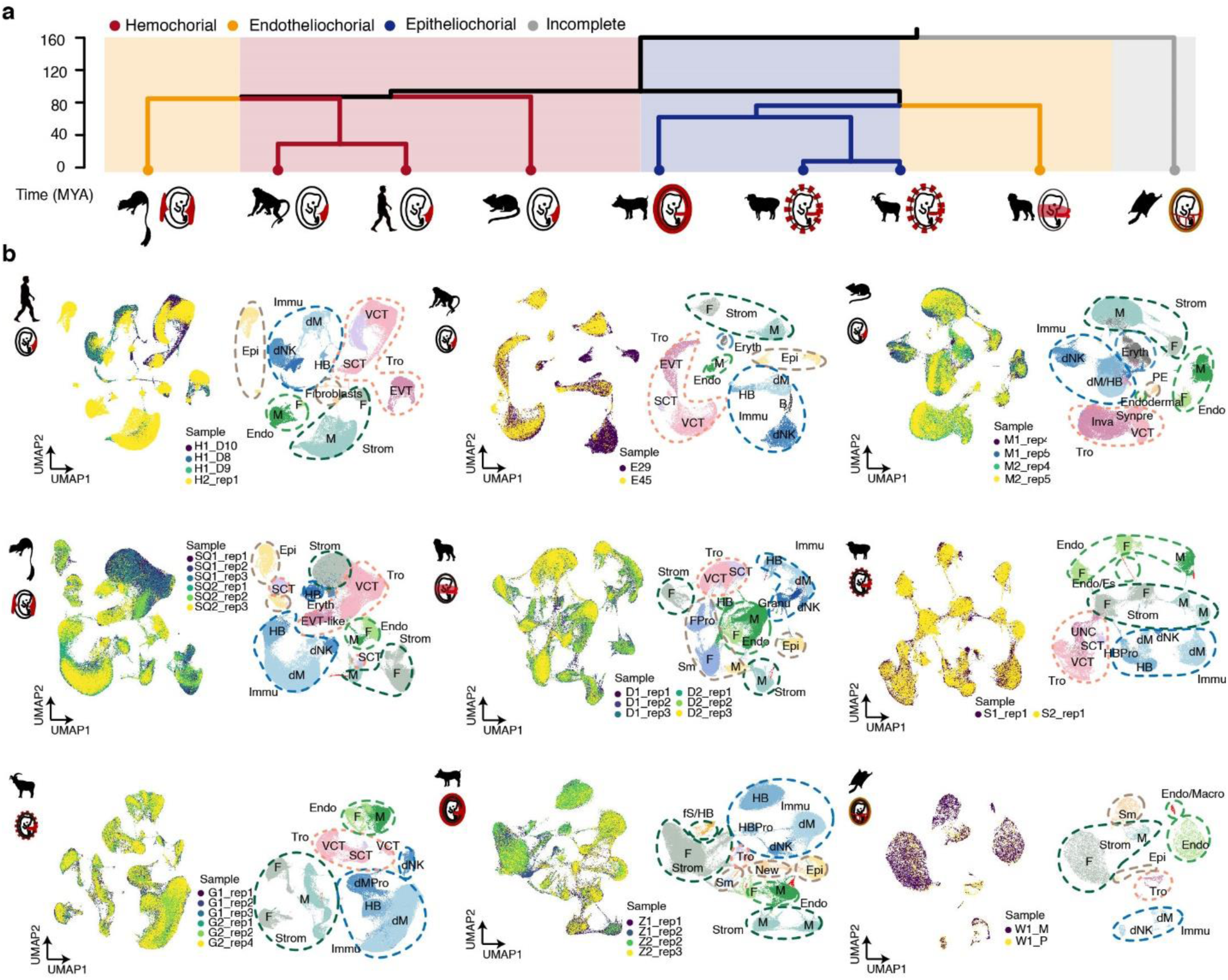
Evolutionary relationships and cross-species single-cell landscapes of the maternal–fetal interface. **a,** Phylogenetic tree illustrating the evolutionary relationships among the mammalian species included in this study. **b,** Species-specific UMAP embeddings of single-cell transcriptomes, colored by biological replicate of origin (left) and annotated cell type (right), across the nine species.

**Extended Data Fig. 3.**
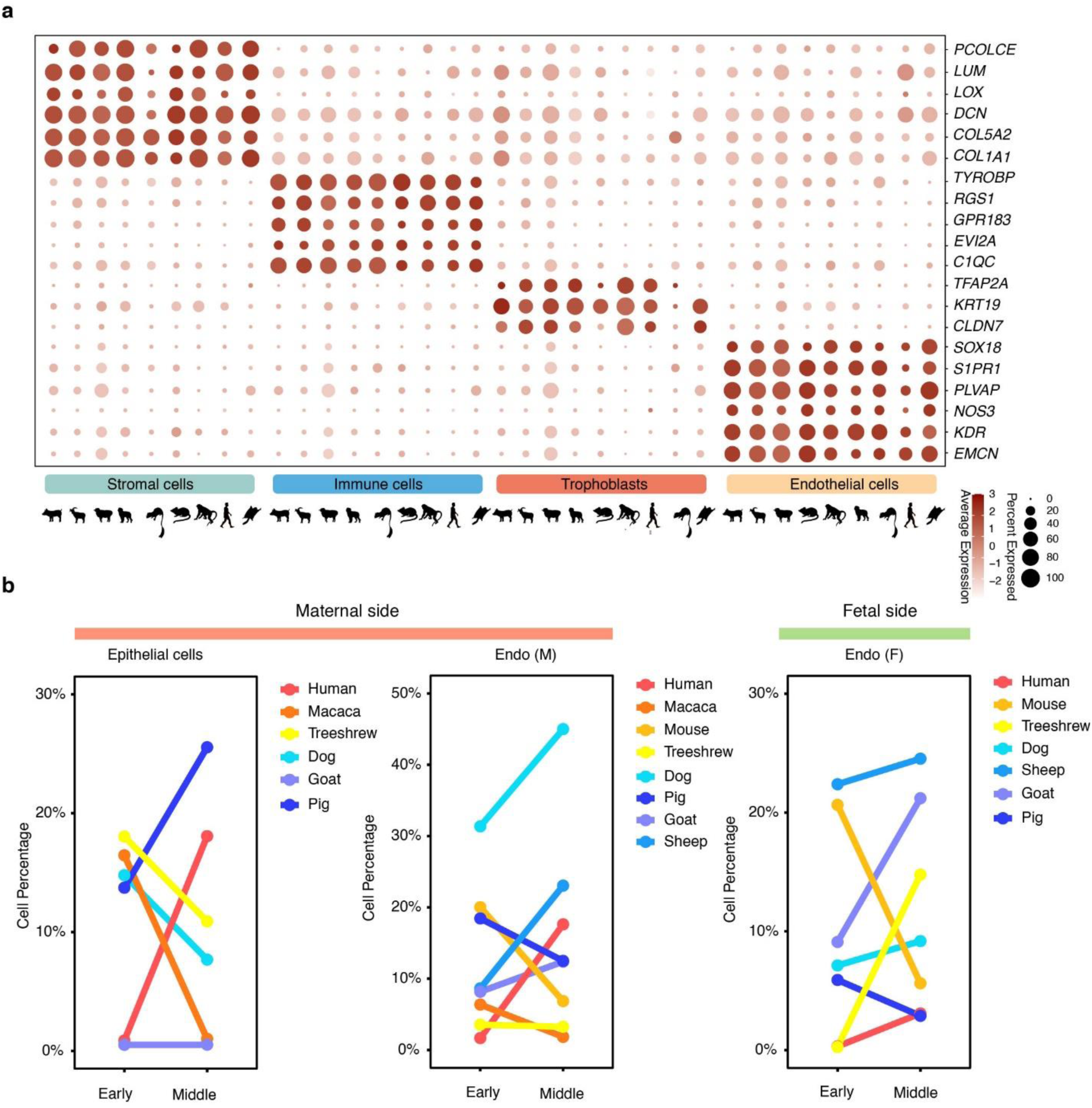
Marker gene expression and cross-species cell-type composition of the maternal–fetal interface. **a,** Bubble plots showing the expression of marker genes for major cell types, including stromal cells, immune cells, trophoblasts, and endothelial cells, across species. **b,** Line charts depicting the proportions of epithelial cells, maternal endothelial cells, and fetal endothelial cells during early and middle gestation across species.

**Extended Data Fig. 4.**
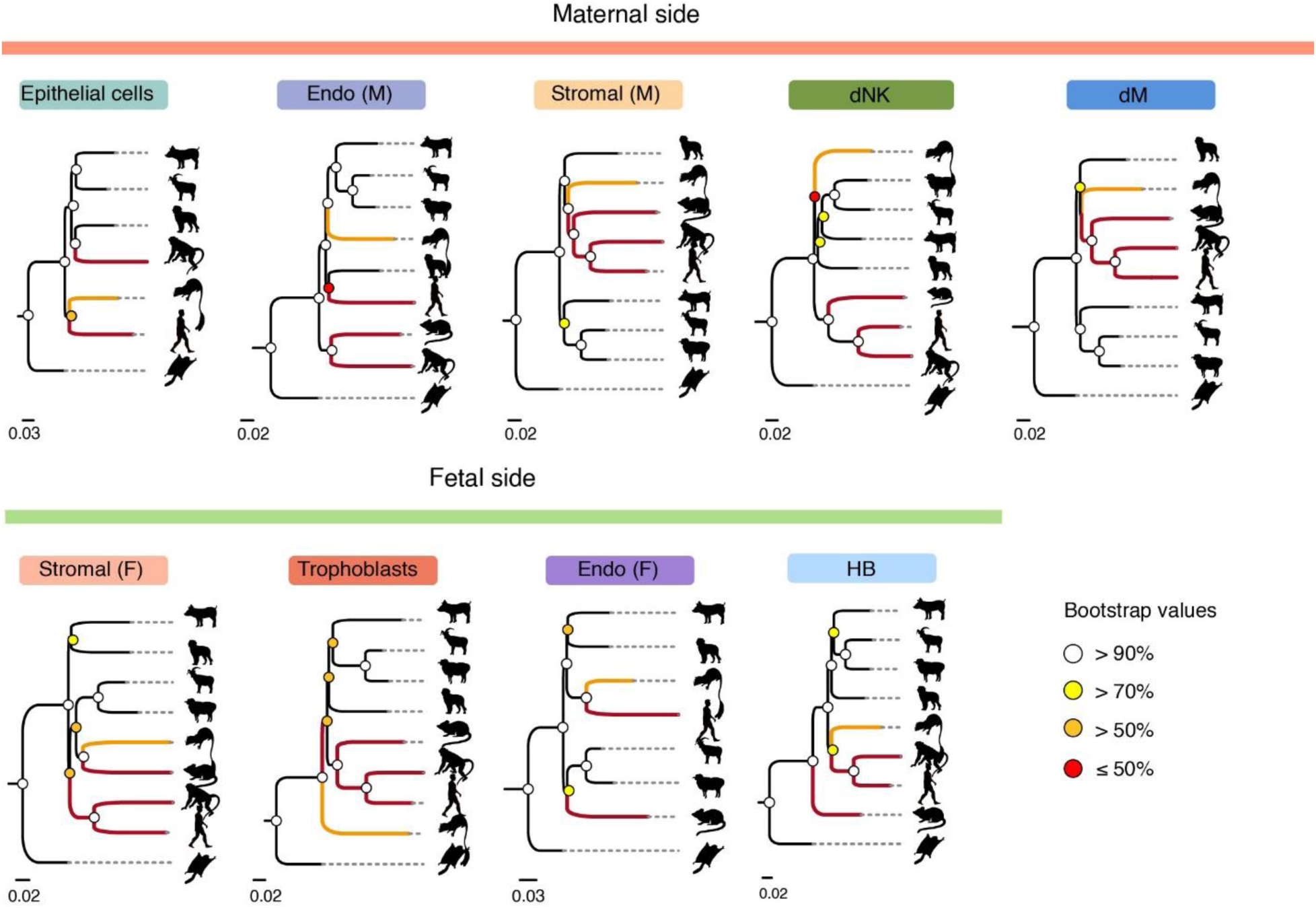
Gene expression-based phylogeny inferred from pseudo-bulk transcriptomes of maternal- and fetal-derived cell types. Bootstrap support values are indicated by circles, with values ≥ 0.9 shown as white-filled circles.

**Extended Data Fig. 5.**
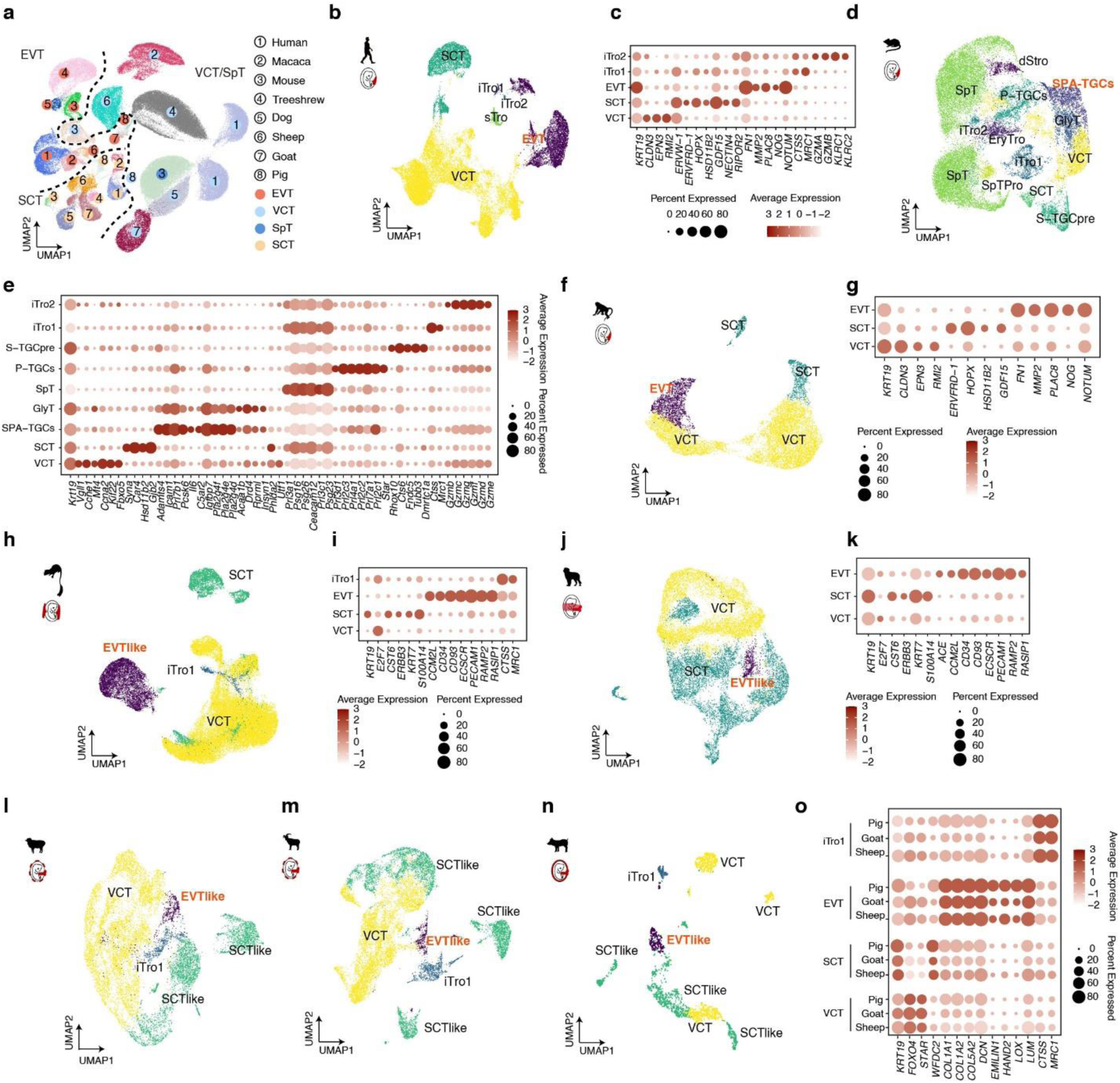
Cross-species single-cell transcriptomic landscapes and marker gene expression of trophoblast subtypes. **a,** Integrated UMAP projection of trophoblast cells from eight species using SATURN. **b–o,** Species-resolved UMAP embeddings and corresponding bubble plots illustrating marker gene expression for distinct trophoblast subtypes in human, macaque, mouse, tree shrew, dog, sheep, goat, and pig, respectively.

**Extended Data Fig. 6.**
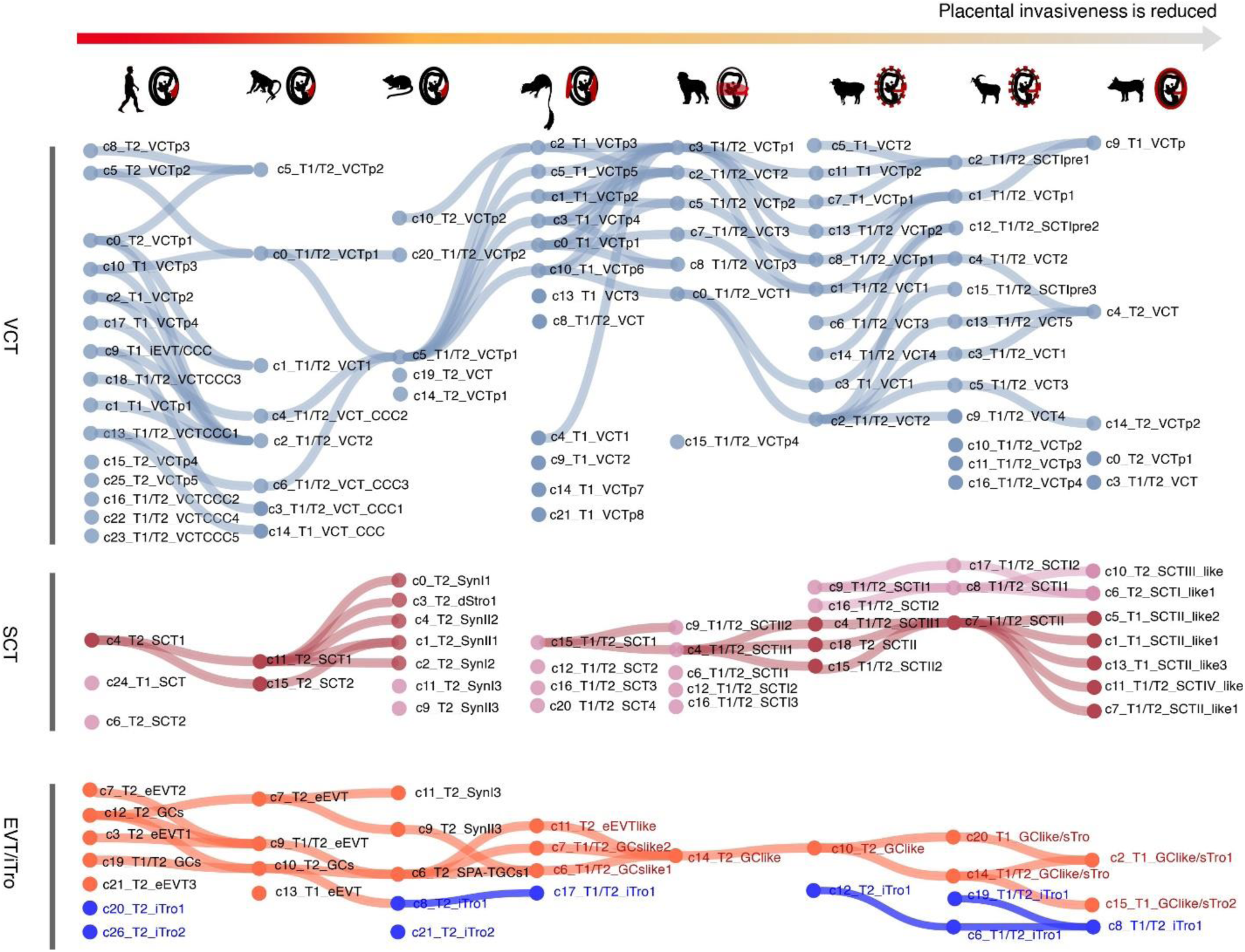
SAMap mapping of cross-species relationships among trophoblast clusters. SAMap was used to map relationships between clusters of villous cytotrophoblasts (VCT), syncytiotrophoblasts (SCT), extravillous trophoblasts (EVT), and trophoblast subpopulations expressing immune markers (iTro) across species, using an alignment threshold of 0.3.

**Extended Data Fig. 7.**
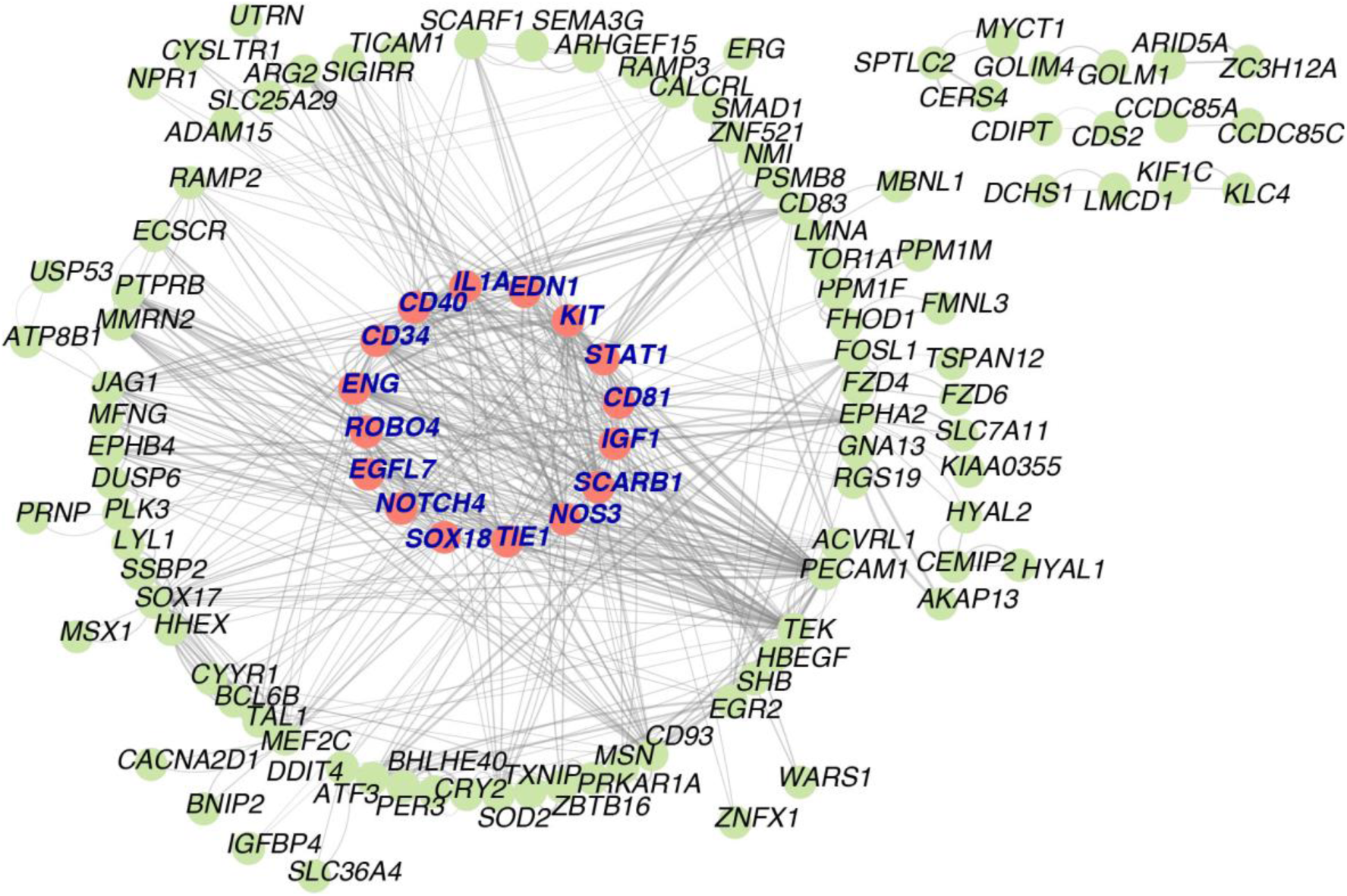
Upregulated genes in EVT-like cells from endotheliochorial placentas and their corresponding protein–protein interaction networks. Genes marked in red circles are key genes in the interaction network, while genes highlighted in red bold font are hub genes. These were derived based on the combined scores generated by MCODE using STRING.

**Extended Data Fig. 8.**
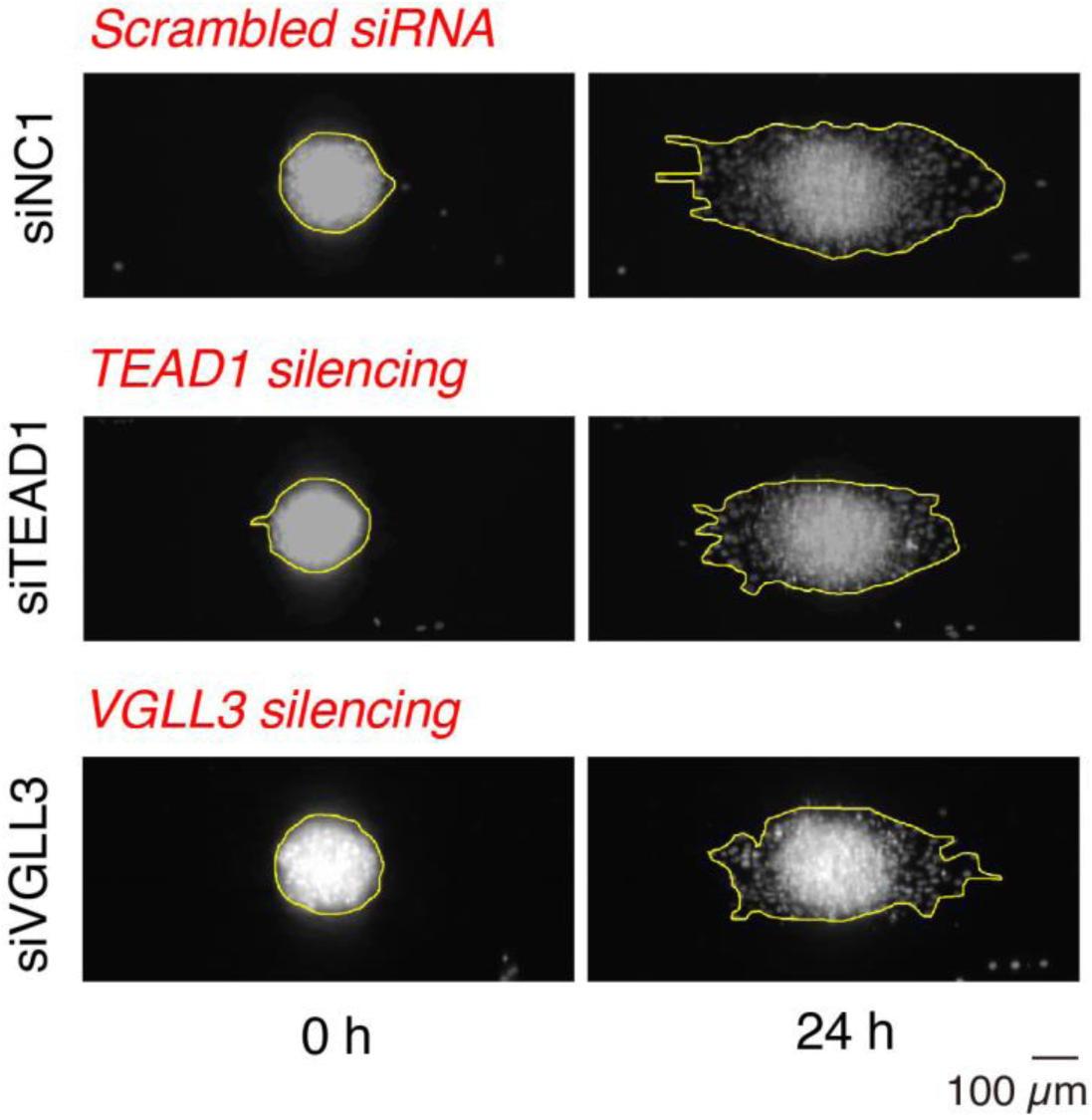
Cross-species comparison of extravillous trophoblasts (EVTs). Fluorescent images showing in-situ *TEAD1* and *VGLL3* genes silencing on human EVT (HTR8-H2B-tdTomato) spheroid invasion into hESF monolayer (black; unlabeled); Spheroid boundaries are manually outlined.

**Extended Data Fig. 9.**
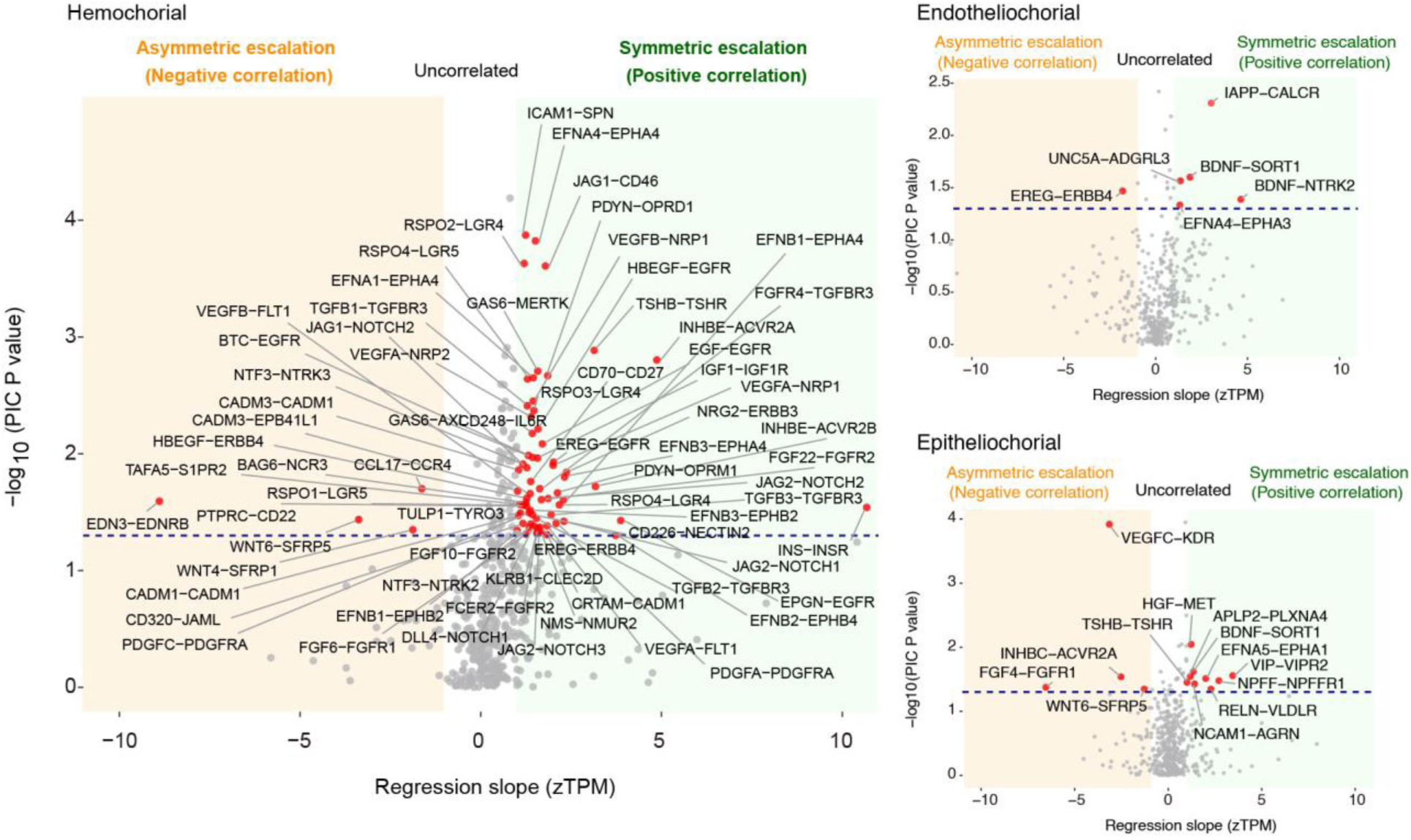
Co-evolution of fetal ligands and maternal receptors assessed by phylogenetically independent contrasts. Volcano plots illustrating co-evolutionary coupling between ligand and receptor expression across hemochorial, endotheliochorial, and epitheliochorial placentas. The vertical axis shows two-sided *P* values testing whether the slope of phylogenetically independent contrasts between ligand and receptor expression differs significantly from zero, while the horizontal axis represents ordinary least squares regression slopes of zTPM-transformed ligand and receptor expression values. Ligand–receptor pairs exhibiting strong evolutionary coupling are highlighted within shaded regions (orange, negative correlation; green, positive correlation), with pairs showing stronger statistical support indicated in red. The blue dashed line denotes a *P* value threshold of 0.05.

**Extended Data Fig. 10.**
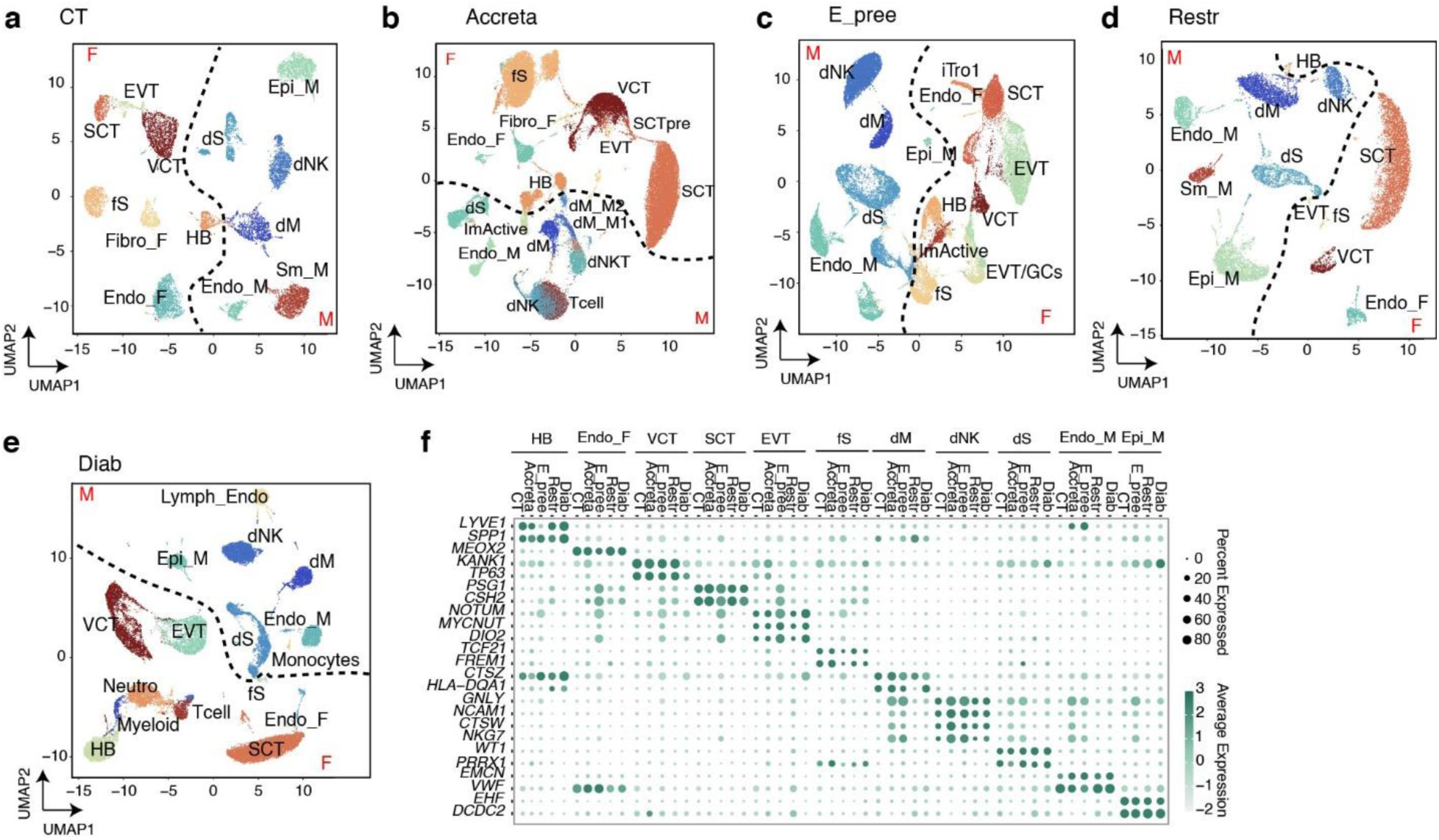
Single-cell transcriptomic landscapes of the maternal–fetal interface in healthy and complicated pregnancies. a–e, UMAP projections of single-cell RNA-seq data, colored by annotated cell type (right), for healthy controls (CT) (a) and pregnancies complicated by placenta accreta spectrum disorders (Accreta) (b), early-onset severe pre-eclampsia (E_pree) (c), fetal growth restriction (Restr) (d), and gestational diabetes (Diab) (e). **f,** Bubble plots illustrating marker gene expression patterns across cell types in healthy and complicated pregnancies.

**Extended Data Fig. 11.**
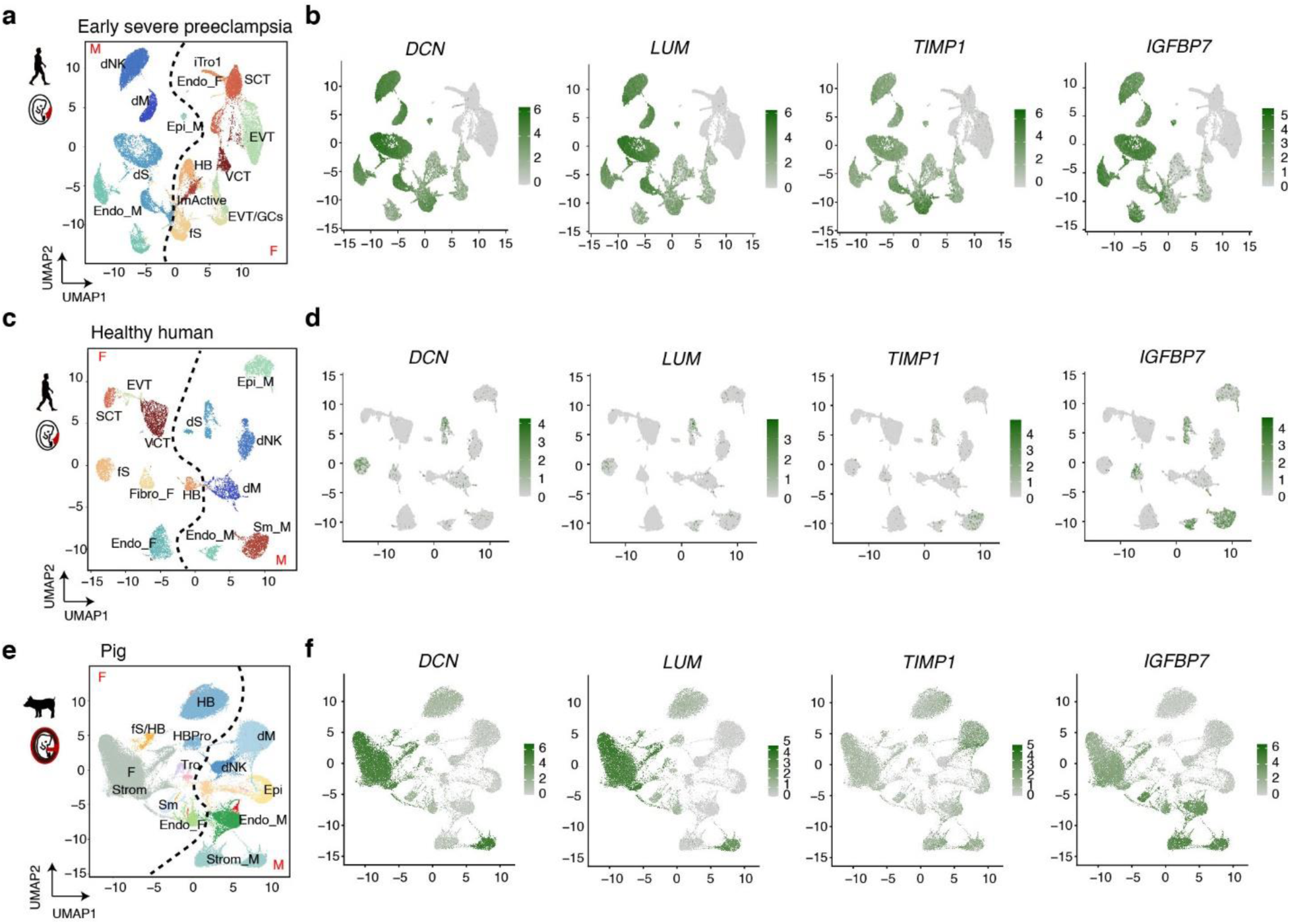
UMAP feature plots of *DCN*, *LUM*, *TIMP1*, and *IGFBP7* expression in healthy controls, early-onset severe pre-eclampsia, and pig placentas. a,c,e. UMAP embeddings with cell-type annotations for early-onset severe pre-eclampsia (a), healthy controls (c), and pig samples (e). **b,d,f** Corresponding UMAP feature plots showing the expression distributions of *DCN*, *LUM*, *TIMP1*, and *IGFBP7* in early-onset severe pre-eclampsia (b), healthy controls (d), and pig samples (f), respectively.

**Extended Data Fig. 12.**
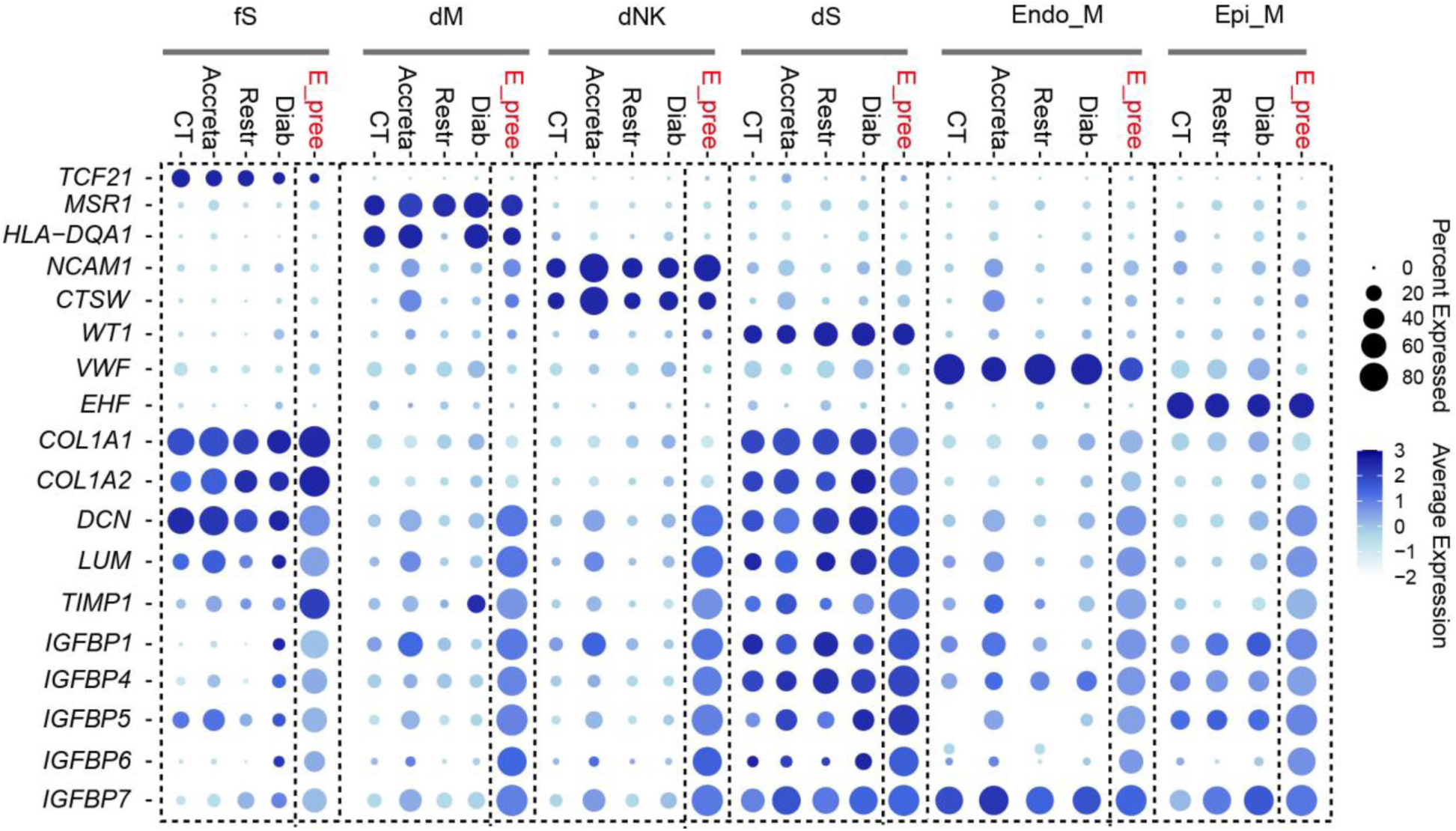
Altered marker gene expression across other cell types in healthy and complicated pregnancies. Bubble plots showing the expression of marker genes for fetal stromal cells (fS), decidual macrophages (dM), decidual NK cells (dNK), decidual stromal cells (dS), maternal endothelial cells (Endo_M), and maternal epithelial cells (Epi_M) across healthy controls (CT) and four major pregnancy complications: placenta accreta spectrum disorders (Accreta), early-onset severe pre-eclampsia (E_pree), fetal growth restriction (Restr), and gestational diabetes (Diab). In addition, the expression of marker genes for clusters c7 and c8 identified in Fig 4e is shown for these cell types in both control and disease conditions.

**Extended Data Fig. 13.**
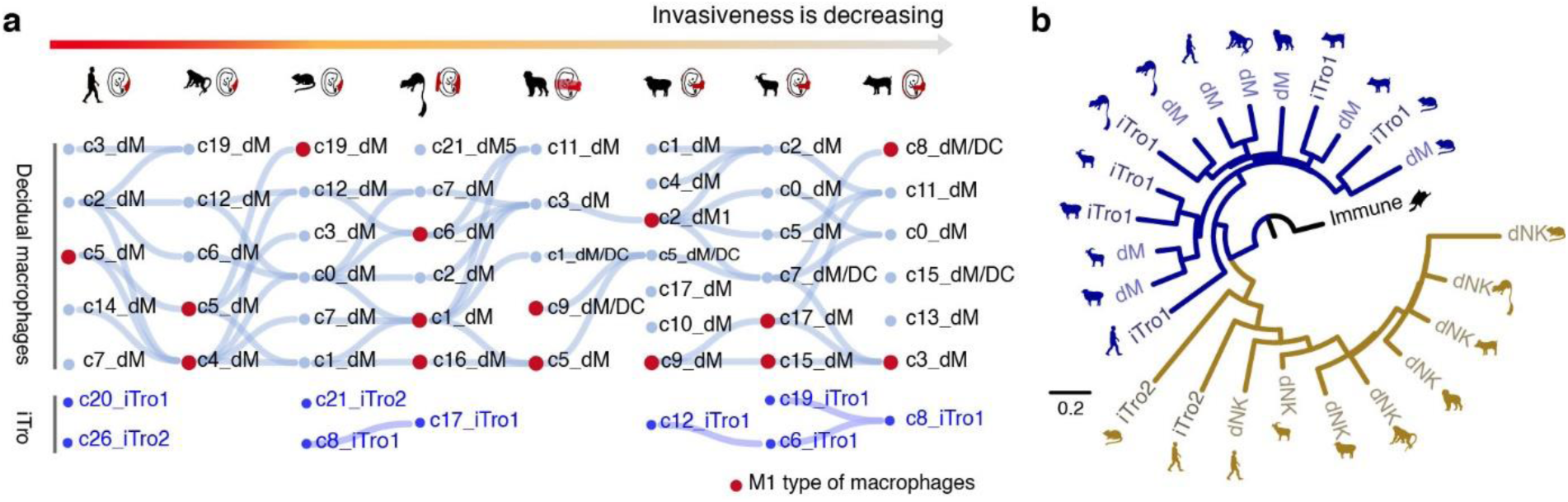
Cross-species relationships between dM, dNK and trophoblast subpopulations (iTro1 and iTro2) expressing immune markers revealed by SAMap and phylogenetic analysis. **a,** SAMap-based mapping of cluster correspondences between dM, iTro1 and iTro2 across species, with connections shown for an alignment threshold of 0.3. **b,** Phylogenetic tree constructed from expression matrices of iTro1, iTro2, dM, dNK across species, using immune cells from sugar glider (*Petaurus breviceps*) as the outgroup.

**Extended Data Fig. 14.**
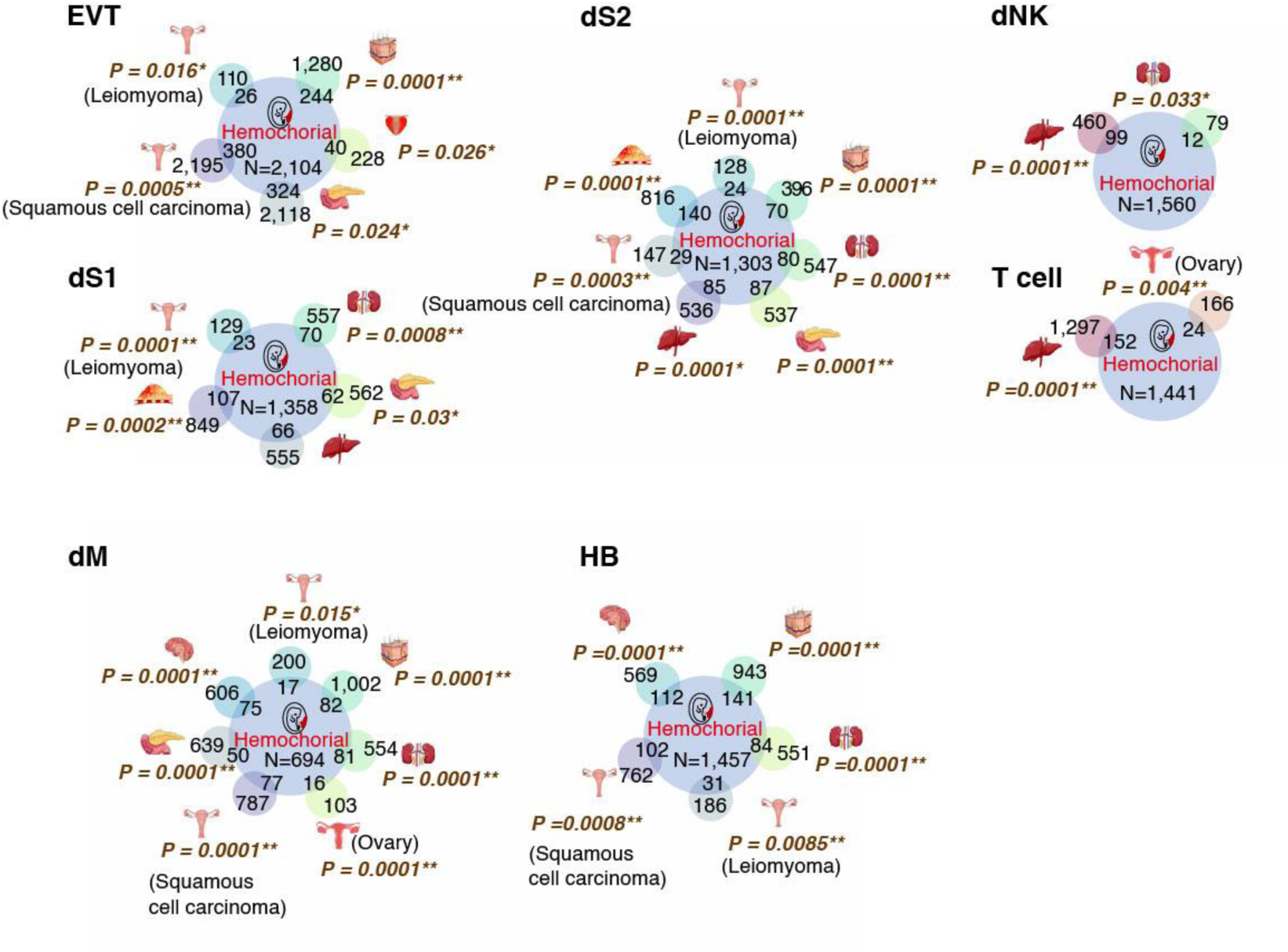
Statistical significance of the overlap between genes upregulated in different tumor types and those upregulated in hemochorial placentas across distinct cell types.

**Extended Data Fig. 15.**
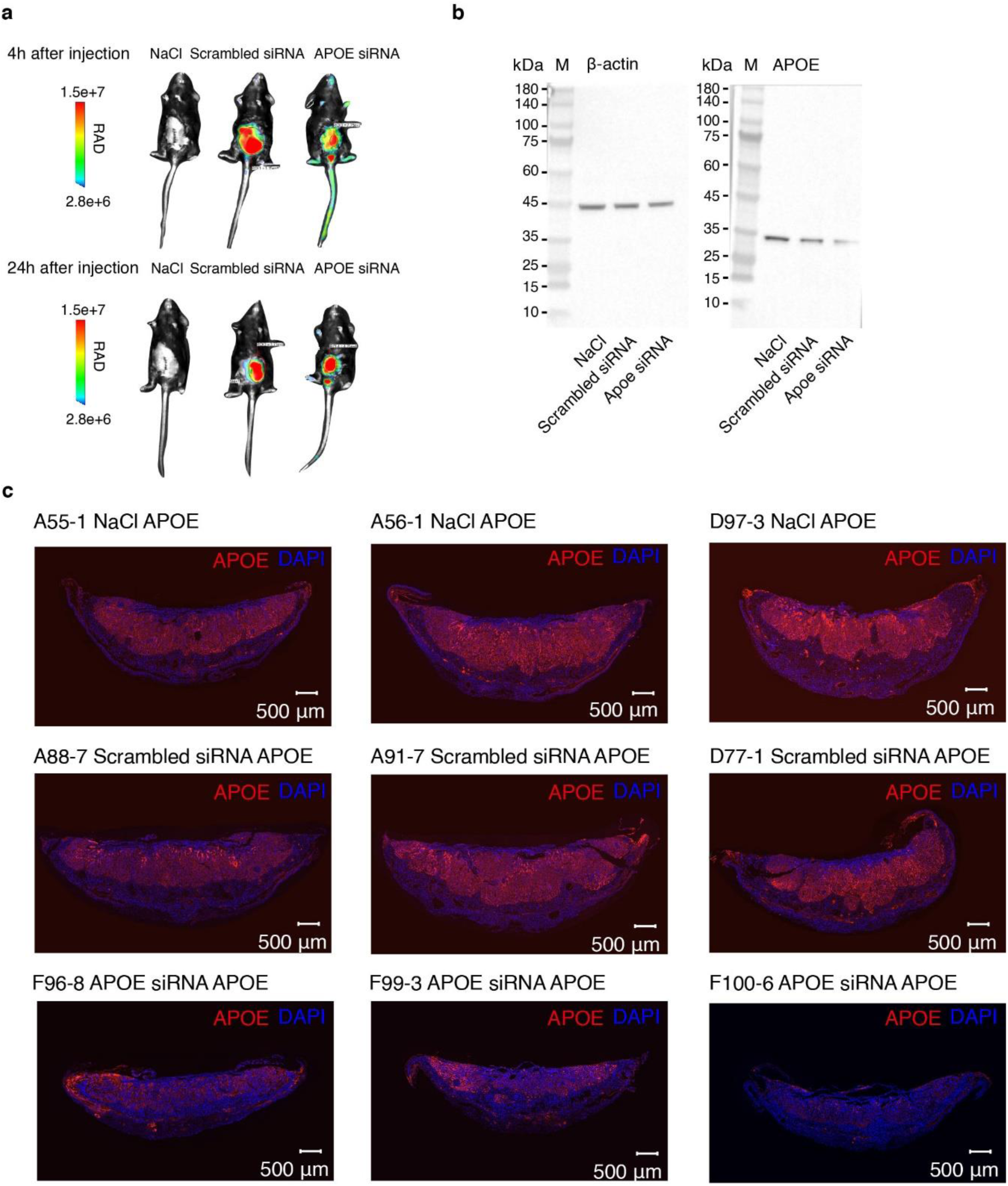
Cy5-traced intra-placental siRNA delivery achieves efficient *APOE* knockdown in mouse placenta as validated by Western blot and immunofluorescence. **a**, Biodistribution of Cy5-siRNA-containing 2′-O-methyl ribose substitutions and 5′ cholesterol modification in NaCl, scrambled siRNA, and APOE-targeting siRNA mice 4 and 24h post *in utero* injection. **b,** The relative expressions of *APOE* and β-actin of E15.5 placenta in NaCl, scrambled siRNA, and APOE-targeting siRNA by western blot. **c,** Immunofluorescence staining for APOE (red) of E15.5 placental sections at 8μm thickness from NaCl, scrambled siRNA, and *APOE* siRNA-treated groups. The prefix “xxx-xxx” refers to the maternal ID, and the suffix refers to the fetal ID.

**Extended Data Fig. 16.**
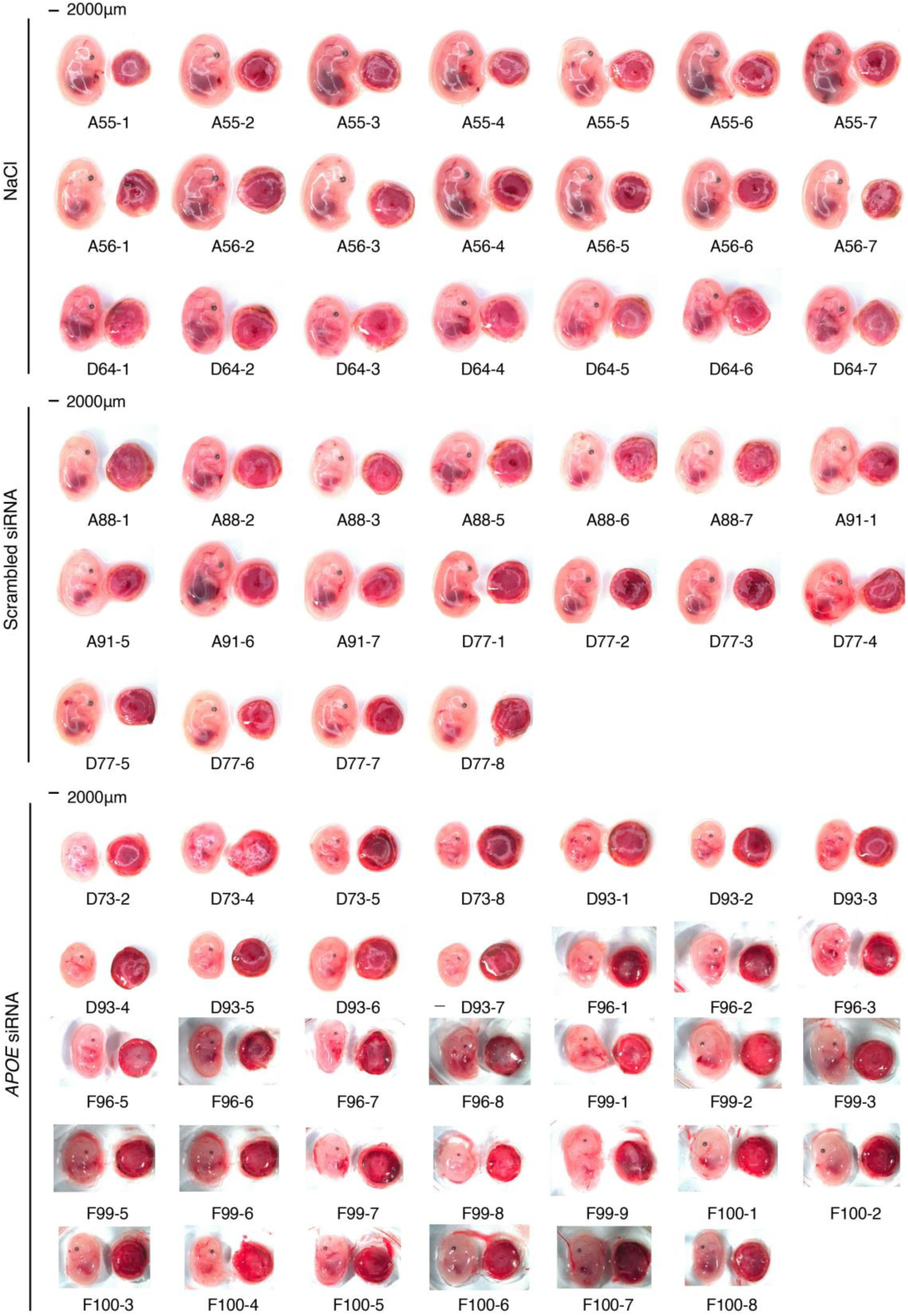
Gross morphology and quantification of E15.5 embryos and placentas from NaCl, scrambled siRNA, and *APOE* siRNA-treated groups. The prefix “xxx-xxx” refers to the maternal ID, and the suffix refers to the fetal ID.

**Extended Data Fig. 17.**
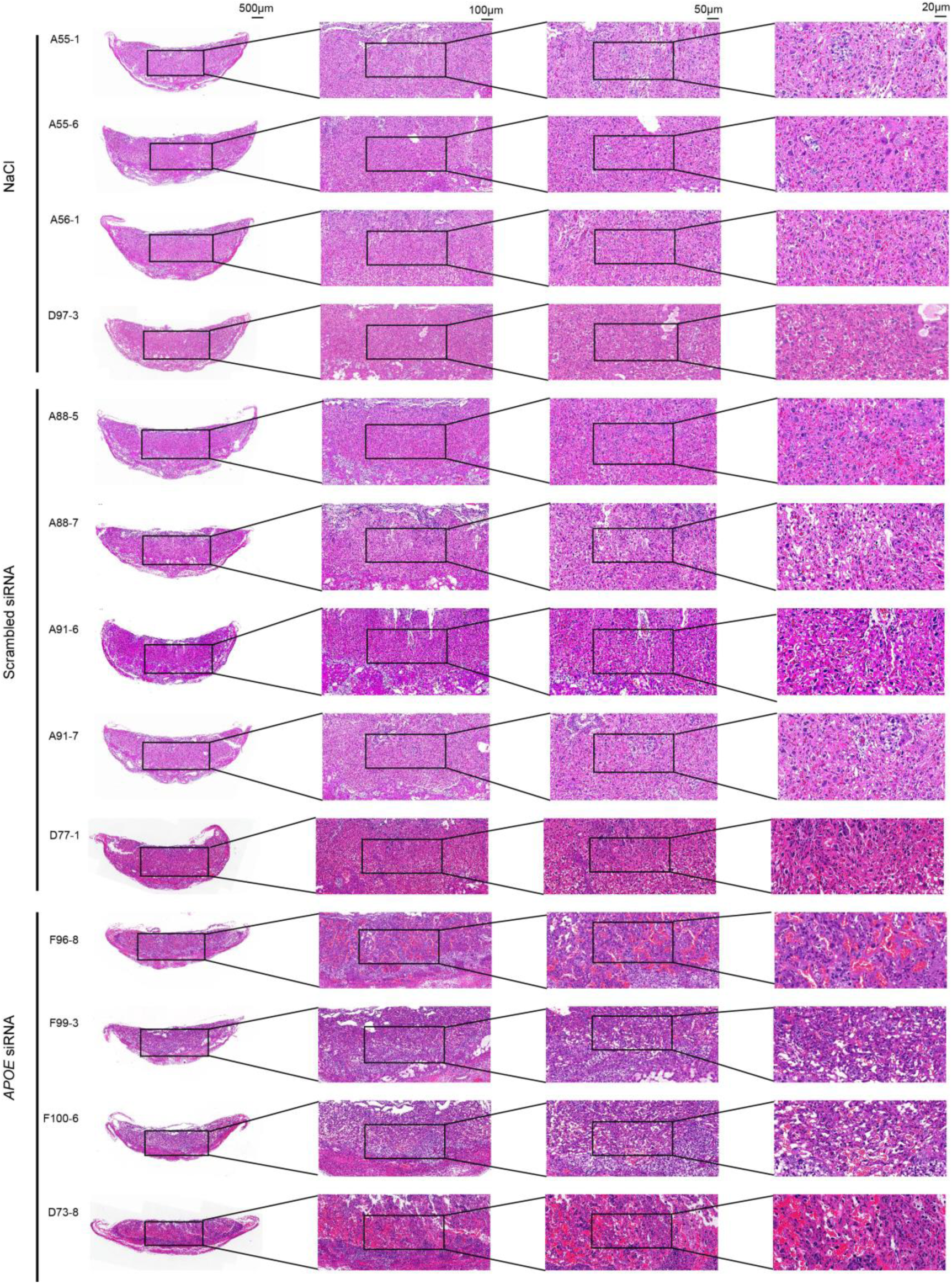
Hematoxylin and eosin (H&E) staining of E15.5 placentas at 8 μm thickness from NaCl, scrambled siRNA, and *APOE* siRNA treated groups. The prefix “xxx-xxx” refers to the maternal ID, and the suffix refers to the fetal ID.

**Extended Data Fig. 18.**
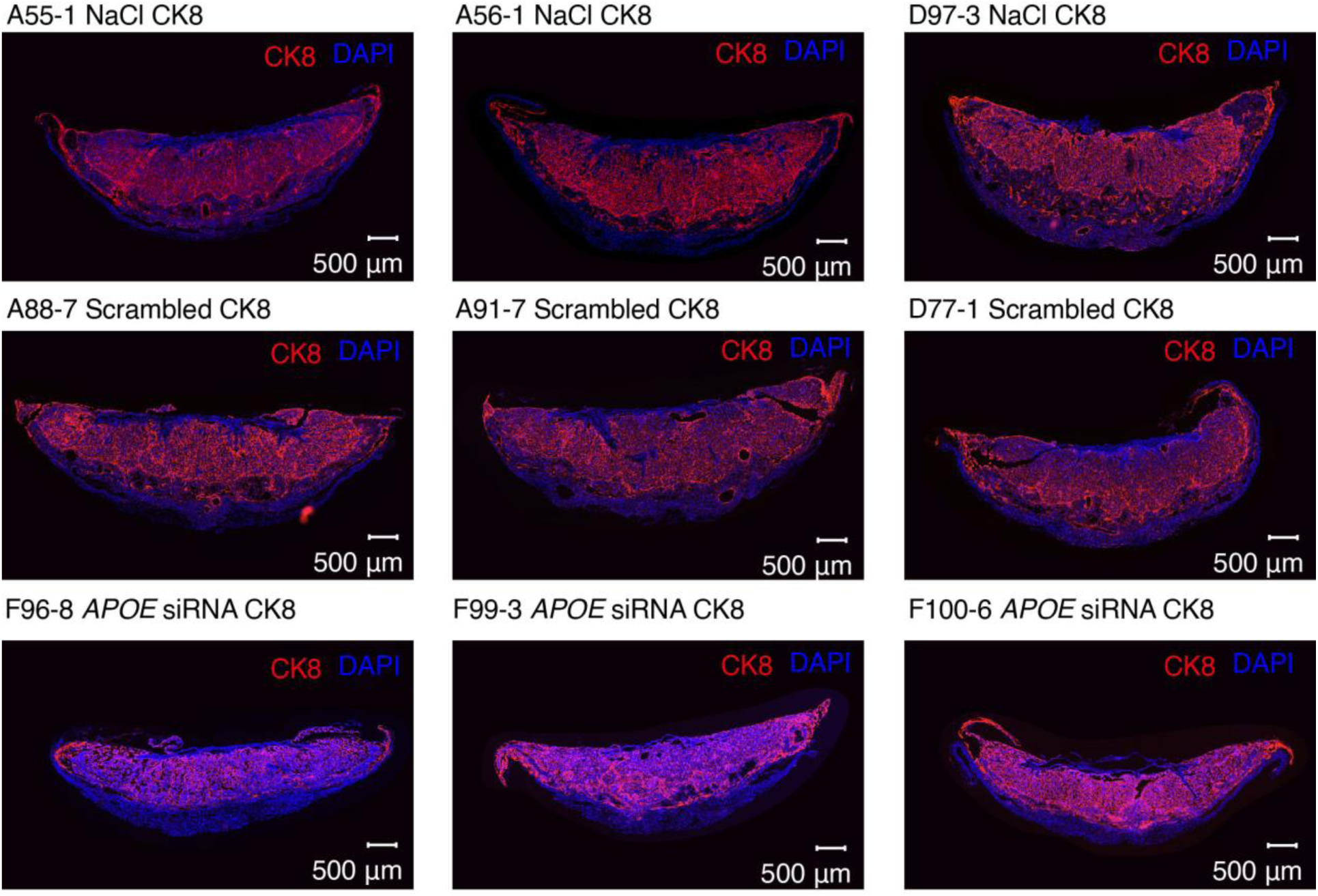
Immunofluorescence staining for CK8 (red) of E15.5 placental sections at 8 μm thickness from NaCl, scrambled siRNA, and *APOE* siRNA-treated groups. The prefix “xxx-xxx” refers to the maternal ID, and the suffix refers to the fetal ID.

**Extended Data Fig. 19.**
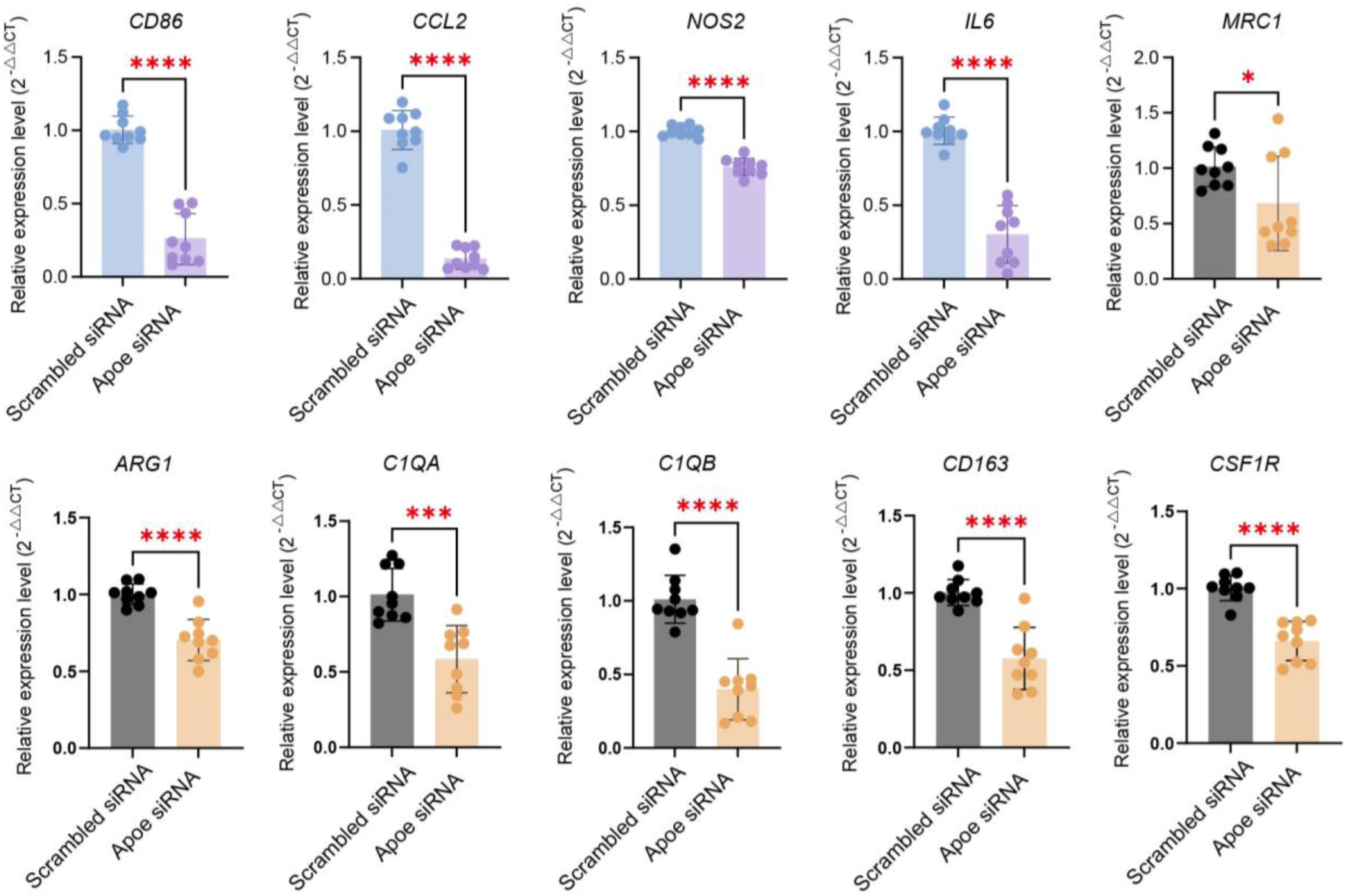
Expression of M1-type and M2-type marker genes of macrophage in E15.5 placentas treated with NaCl, scrambled siRNA, and *APOE*-targeting siRNA. qPCR analysis of M1-type macrophage genes (*CD86, CCL2, NOS2* and *IL6*) and M2-type macrophage genes (*MRC1, ARG1, C1QA, C1QB, CD163* and *CSF1R*) in E15.5 placentas treated with scrambled siRNA (negative control, *n* = 9) and *APOE* siRNA (*n* = 9). Statistical significance was assessed using an unpaired *t* test comparing scrambled siRNA (negative control) and *APOE* siRNA groups. Asterisks indicate significance levels: \*\*\*\**P* < 0.0001, \*\*\**P* < 0.001, \*\**P* < 0.01 and \**P* < 0.05.

**Extended Data Fig. 20.**
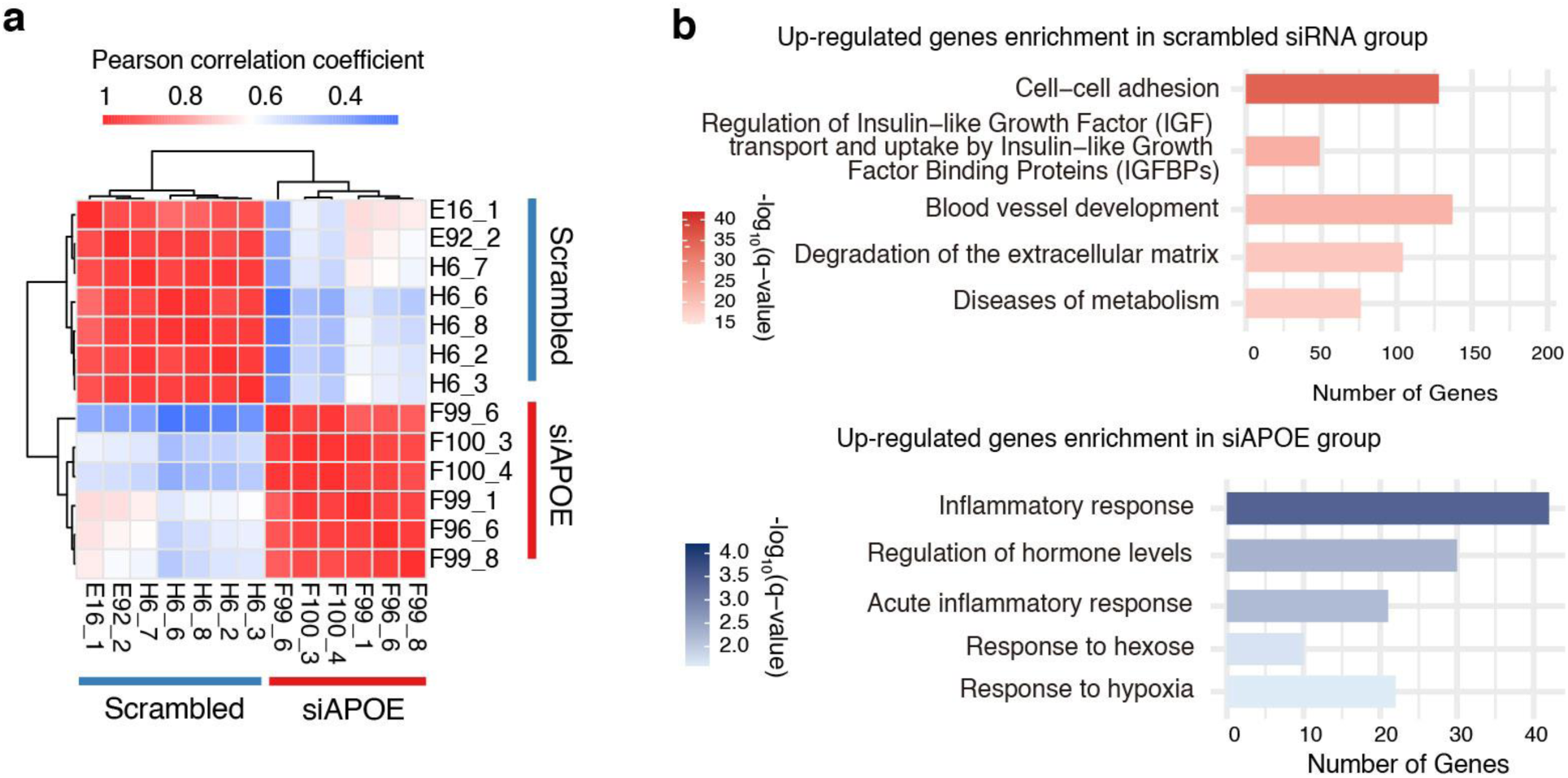
Bulk RNA-seq on placentas from the scrambled-control and *APOE-*knockdown groups. **a**, Pearson correlation coefficients between samples treated with scrambled siRNA and APOE-targeting siRNA. **b**, Pathway enrichment analysis of genes significantly upregulated in scrambled siRNA and APOE-targeting siRNA-treated group, respectively.

## Notes

### Competing Interest Statement

The authors have declared no competing interest.

## References

1. Roberts, R. M., Green, J. A. & Schulz, L. C. The evolution of the placenta. Reproduction 152, R179–189 (2016).

2. Mika, K., Whittington, C. M., McAllan, B. M. & Lynch, V. J. Gene expression phylogenies and ancestral transcriptome reconstruction resolves major transitions in the origins of pregnancy. Elife 11, e74297 (2022).

3. Griffith, O. W. & Wagner, G. P. The placenta as a model for understanding the origin and evolution of vertebrate organs. Nat Ecol Evol 1, 72 (2017).

4. Haig, D. Genetic conflicts in human pregnancy. Q Rev Biol 68, 495–532 (1993).

5. Moffett, A. & Loke, C. Immunology of placentation in eutherian mammals. Nat Rev Immunol 6, 584–594 (2006).

6. Hemberger, M. Immune balance at the foeto-maternal interface as the fulcrum of reproductive success. J Reprod Immunol 97, 36–42 (2013).

7. Carter, A. M. & Mess, A. Evolution of the placenta in eutherian mammals. Placenta 28, 259–262 (2007).

8. Grosser, O. Frühentwicklung, Eihautbildung Und Placentation Des Menschen Und Der Säugetiere. vol. 5 (Bergmann, 1927).

9. Wildman, D. E. et al. Evolution of the mammalian placenta revealed by phylogenetic analysis. Proc Natl Acad Sci U S A 103, 3203–3208 (2006).

10. Trivers, R. L. Parent-offspring conflict. American zoologist 14, 249–264 (1974).

11. Crespi, B. & Semeniuk, C. Parent-offspring conflict in the evolution of vertebrate reproductive mode. Am Nat 163, 635–653 (2004).

12. Kshitiz, null et al. Evolution of placental invasion and cancer metastasis are causally linked. Nat Ecol Evol 3, 1743–1753 (2019).

13. Wagner, G. P., Kshitiz, null, Dighe, A. & Levchenko, A. The Coevolution of Placentation and Cancer. Annu Rev Anim Biosci 10, 259–279 (2022).

14. Vornic, I. et al. Molecular Insights into Human Placentation: From Villous Morphogenesis to Pathological Pathways and Translational Biomarkers. Int J Mol Sci 26, 9483 (2025).

15. Holtan, S. G., Creedon, D. J., Haluska, P. & Markovic, S. N. Cancer and pregnancy: parallels in growth, invasion, and immune modulation and implications for cancer therapeutic agents. Mayo Clin Proc 84, 985–1000 (2009).

16. Ferretti, C., Bruni, L., Dangles-Marie, V., Pecking, A. P. & Bellet, D. Molecular circuits shared by placental and cancer cells, and their implications in the proliferative, invasive and migratory capacities of trophoblasts. Hum Reprod Update 13, 121–141 (2007).

17. Chew, L. C., Osuchukwu, O. O., Reed, D. J. & Verma, R. P. Fetal Growth Restriction. In StatPearls [Internet] (StatPearls Publishing, 2024).

18. Staff, A. C. et al. Failure of physiological transformation and spiral artery atherosis: their roles in preeclampsia. Am J Obstet Gynecol 226, S895–S906 (2022).

19. Windsperger, K. et al. Extravillous trophoblast invasion of venous as well as lymphatic vessels is altered in idiopathic, recurrent, spontaneous abortions. Hum Reprod 32, 1208–1217 (2017).

20. Illsley, N. P., DaSilva-Arnold, S. C., Zamudio, S., Alvarez, M. & Al-Khan, A. Trophoblast invasion: Lessons from abnormally invasive placenta (placenta accreta). Placenta 102, 61–66 (2020).

21. Elliot, M. G. & Crespi, B. J. Genetic recapitulation of human pre-eclampsia risk during convergent evolution of reduced placental invasiveness in eutherian mammals. Philos Trans R Soc Lond B Biol Sci 370, 20140069 (2015).

22. Beard, J. CANCER GENESIS. The Lancet 165, 56–57 (1905).

23. Pavličev, M. et al. Single-cell transcriptomics of the human placenta: inferring the cell communication network of the maternal-fetal interface. Genome Res 27, 349–361 (2017).

24. Scott, R. L. et al. Conservation at the uterine–placental interface. Proceedings of the National Academy of Sciences 119, e2210633119 (2022).

25. Stadtmauer, D. J. et al. Cell type and cell signalling innovations underlying mammalian pregnancy. Nat Ecol Evol 9, 1469–1486 (2025).

26. Tan, G. et al. Cross-species insights into placental evolution and diseases at the single-cell resolution. Nat Commun 10.1038/s41467-026-72652-w (2026) doi:10.1038/s41467-026-72652-w.

27. Afzal, J. et al. Evidence for coopetition at the maternal-fetal interface shaping placental invasion. Proc Natl Acad Sci U S A 122, e2323038122 (2025).

28. Yu, J. et al. Progestogen-driven B7-H4 contributes to onco-fetal immune tolerance. Cell 187, 4713–4732.e19 (2024).

29. Rosen, Y. et al. Toward universal cell embeddings: integrating single-cell RNA-seq datasets across species with SATURN. Nat Methods 21, 1492–1500 (2024).

30. Beal, J. R., Mukherjee, P., Bagchi, I. C. & Bagchi, M. K. Deciphering the maternal uterine signals that shape placenta development. Reproduction 171, xaag053 (2026).

31. Horn, L. C., Emmrich, P. & Bilek, K. The early placental trophoblast. I. Normal development and endometrial and non-tumor-induced disorders. Zentralbl Gynakol 118, 487–497 (1996).

32. Kumar, S. et al. TimeTree 5: An Expanded Resource for Species Divergence Times. Mol Biol Evol 39, msac174 (2022).

33. Wynn, R. M. & Corbett, J. R. Ultrastructure of the canine placenta and amnion. Am J Obstet Gynecol 103, 878–887 (1969).

34. Luckhardt, M., Kaufmann, P. & Elger, W. The structure of the tupaia placenta: I. Histology and vascularisation. Anat Embryol 171, 201–210 (1985).

35. Lin, K. C., Park, H. W. & Guan, K.-L. Regulation of the Hippo pathway transcription factor TEAD. Trends Biochem Sci 42, 862–872 (2017).

36. Plazyo, O. et al. Defective Trophoblast Differentiation, Endothelial Dysfunction, and Immune Dysregulation in Preeclampsia Coalesce on a Placental VGLL3-Centered Gene Network. Circulation 0,.

37. Wenqiang, D. et al. Scar matrix drives Piezo1 mediated stromal inflammation leading to placenta accreta spectrum. Nat Commun 15, 8379 (2024).

38. Meinhardt, G. et al. The multifaceted roles of the transcriptional coactivator TAZ in extravillous trophoblast development of the human placenta. Proc Natl Acad Sci U S A 122, e2426385122 (2025).

39. Gu, B. et al. The TEA domain transcription factors TEAD1 and TEAD3 and WNT signaling determine HLA-G expression in human extravillous trophoblasts. Proc Natl Acad Sci U S A 122, e2425339122 (2025).

40. Sato, Y. et al. Trophoblasts acquire a chemokine receptor, CCR1, as they differentiate towards invasive phenotype. Development 130, 5519–5532 (2003).

41. Zhao, X. et al. Up-regulation of microRNA-135 or silencing of PCSK6 attenuates inflammatory response in preeclampsia by restricting NLRP3 inflammasome. Mol Med 27, 82 (2021).

42. Pescador, N. et al. Hypoxia promotes glycogen accumulation through hypoxia inducible factor (HIF)-mediated induction of glycogen synthase 1. PLoS One 5, e9644 (2010).

43. Wang, P., Wang, F., Wang, L. & Pan, J. Proprotein convertase subtilisin/kexin type 6 activates the extracellular signal-regulated kinase 1/2 and Wnt family member 3A pathways and promotes in vitro proliferation, migration and invasion of breast cancer MDA-MB-231 cells. Oncol Lett 16, 145–150 (2018).

44. Amabebe, E., Ogidi, H. & Anumba, D. O. Matrix metalloproteinase-induced cervical extracellular matrix remodelling in pregnancy and cervical cancer. Reprod Fertil 3, R177–R191 (2022).

45. Longstreth, J. H. & Wang, K. The role of fibronectin in mediating cell migration. Am J Physiol Cell Physiol 326, C1212–C1225 (2024).

46. Wu, W.-B. et al. Decreased PGF may contribute to trophoblast dysfunction in fetal growth restriction. Reproduction 154, 319–329 (2017).

47. Pagani, E. et al. Placenta growth factor and neuropilin-1 collaborate in promoting melanoma aggressiveness. Int J Oncol 48, 1581–1589 (2016).

48. Fukuda, R., Beppu, S., Hinata, D., Kamada, Y. & Okiyoneda, T. Perturbation of EPHA2 and EFNA1 trans binding amplifies inflammatory response in airway epithelial cells. iScience 28, 111872 (2025).

49. Dunne, P. D. et al. EphA2 Expression Is a Key Driver of Migration and Invasion and a Poor Prognostic Marker in Colorectal Cancer. Clin Cancer Res 22, 230–242 (2016).

50. Inoue, M. et al. Endothelial cell-selective adhesion molecule modulates atherosclerosis through plaque angiogenesis and monocyte-endothelial interaction. Microvasc Res 80, 179–187 (2010).

51. Kanai, S. M. & Clouthier, D. E. Endothelin signaling in development. Development 150, dev201786 (2023).

52. Roberts, D. M. et al. The vascular endothelial growth factor (VEGF) receptor Flt-1 (VEGFR-1) modulates Flk-1 (VEGFR-2) signaling during blood vessel formation. Am J Pathol 164, 1531–1535 (2004).

53. Shattil, S. J. Function and regulation of the beta 3 integrins in hemostasis and vascular biology. Thromb Haemost 74, 149–155 (1995).

54. Wang, Q. et al. AR+TREM2+ macrophage induced pathogenic immunosuppression promotes prostate cancer progression. Nat Commun 16, 6964 (2025).

55. Ma, K., Chen, S., Chen, X., Zhao, X. & Yang, J. CD93 is Associated with Glioma-related Malignant Processes and Immunosuppressive Cell Infiltration as an Inspiring Biomarker of Survivance. J Mol Neurosci 72, 2106–2124 (2022).

56. Cai, Q., Dozmorov, M. & Oh, Y. IGFBP-3/IGFBP-3 Receptor System as an Anti-Tumor and Anti-Metastatic Signaling in Cancer. Cells 9, 1261 (2020).

57. Cabezón, R. et al. MERTK as negative regulator of human T cell activation. J Leukoc Biol 97, 751–760 (2015).

58. Batlle, E. & Massagué, J. Transforming Growth Factor-β Signaling in Immunity and Cancer. Immunity 50, 924–940 (2019).

59. Zhang, T., Yu, J., Wang, G. & Zhang, R. Amyloid precursor protein binds with TNFRSF21 to induce neural inflammation in Alzheimer’s Disease. Eur J Pharm Sci 157, 105598 (2021).

60. Johnson, P. & Ruffell, B. CD44 and its role in inflammation and inflammatory diseases. Inflamm Allergy Drug Targets 8, 208–220 (2009).

61. Kwon, B. CD137-CD137 Ligand Interactions in Inflammation. Immune Netw 9, 84–89 (2009).

62. Bassiouny, A. R., Zaky, A. & Kandeel, K. M. Alteration of AP-endonuclease1 expression in curcumin-treated fibrotic rats. Ann Hepatol 10, 516–530 (2011).

63. Dawkins, R. & Krebs, J. R. Arms races between and within species. Proc R Soc Lond B Biol Sci 205, 489–511 (1979).

64. Felsenstein, J. Phylogenies and the Comparative Method. The American Naturalist 125, 1–15 (1985).

65. Afzal, J. et al. Evidence for coopetition at the maternal-fetal interface shaping placental invasion. Proc Natl Acad Sci U S A 122, e2323038122 (2025).

66. Yoshie, M. et al. Small GTP-binding protein Rap1 mediates EGF and HB-EGF signaling and modulates EGF receptor expression in HTR-8/SVneo extravillous trophoblast cells. Reprod Med Biol 22, e12537 (2023).

67. Xie, Y. et al. LGR5 promotes tumorigenicity and invasion of glioblastoma stem-like cells and is a potential therapeutic target for a subset of glioblastoma patients. J Pathol 247, 228–240 (2019).

68. Zhu, Z., Yu, S., Niu, K. & Wang, P. LGR5 promotes invasion and migration by regulating YAP activity in hypopharyngeal squamous cell carcinoma cells under inflammatory condition. PLoS One 17, e0275679 (2022).

69. Zheng, Z. et al. Heterogeneous expression of Lgr5 as a risk factor for focal invasion and distant metastasis of colorectal carcinoma. Oncotarget 9, 30025–30033 (2018).

70. Rose, K. W., Taye, N., Karoulias, S. Z. & Hubmacher, D. Regulation of ADAMTS proteases. Frontiers in molecular biosciences 8, 701959 (2021).

71. Yang, H., Deng, Y., Dong, Y., Ma, Y. & Yang, L. Identification and Validation of Prognostic Markers for Endometriosis-Associated Ovarian Cancer. Int J Med Sci 21, 1903–1914 (2024).

72. Lu, Y., Zhang, S., Wang, Y., Ren, X. & Han, J. Molecular mechanisms and clinical manifestations of rare genetic disorders associated with type I collagen. Intractable Rare Dis Res 8, 98–107 (2019).

73. Mao, L. et al. Decorin deficiency promotes epithelial-mesenchymal transition and colon cancer metastasis. Matrix Biol 95, 1–14 (2021).

74. Gesteira, T. F., Verma, S. & Coulson-Thomas, V. J. Small leucine rich proteoglycans: Biology, function and their therapeutic potential in the ocular surface. Ocul Surf 29, 521–536 (2023).

75. Arpino, V., Brock, M. & Gill, S. E. The role of TIMPs in regulation of extracellular matrix proteolysis. Matrix Biol 44–46, 247–254 (2015).

76. Yu, J.-T. et al. Renal tubular epithelial IGFBP7 interacts with PKM2 to drive renal lipid accumulation and fibrosis. Mol Ther 33, 3757–3777 (2025).

77. Zhu, T. et al. IGFBP7: A novel biomarker involved in a positive feedback loop with TGF-β1 in idiopathic pulmonary fibrosis. Cell Signal 133, 111867 (2025).

78. Feng, Y. et al. Collagen I Induces Preeclampsia-Like Symptoms by Suppressing Proliferation and Invasion of Trophoblasts. Front Endocrinol (Lausanne) 12, 664766 (2021).

79. Baxter, R. C. Signaling Pathways of the Insulin-like Growth Factor Binding Proteins. Endocr Rev 44, 753–778 (2023).

80. Garrido-Gimenez, C. & Alijotas-Reig, J. Recurrent miscarriage: causes, evaluation and management. Postgrad Med J 91, 151–162 (2015).

81. Shi, Y. et al. Research trends and hotspots of recurrent spontaneous abortion with immune dysfunction: A bibliometric analysis from 2004 to 2024. Medicine (Baltimore) 104, e43059 (2025).

82. Zhang, L. et al. Deficient extravillous trophoblast invasion caused by impaired sialylation–Siglec-7 interaction contributes to recurrent pregnancy loss. Cell Death Dis 17, 291 (2026).

83. Zheng, Y. et al. Characterization of placental and decidual cell development in early pregnancy loss by single-cell RNA sequencing. Cell Biosci 12, 168 (2022).

84. Hanahan, D. & Weinberg, R. A. Hallmarks of cancer: the next generation. Cell 144, 646–674 (2011).

85. Valastyan, S. & Weinberg, R. A. Tumor metastasis: molecular insights and evolving paradigms. Cell 147, 275–292 (2011).

86. Knöfler, M. et al. Human placenta and trophoblast development: key molecular mechanisms and model systems. Cell Mol Life Sci 76, 3479–3496 (2019).

87. Liot, S. et al. Stroma Involvement in Pancreatic Ductal Adenocarcinoma: An Overview Focusing on Extracellular Matrix Proteins. Front Immunol 12, 612271 (2021).

88. Jena, M. K., Nayak, N., Chen, K. & Nayak, N. R. Role of Macrophages in Pregnancy and Related Complications. Arch Immunol Ther Exp (Warsz) 67, 295–309 (2019).

89. DeNardo, D. G. & Ruffell, B. Macrophages as regulators of tumor immunity and immunotherapy. Nat Rev Immunol 19, 369–382 (2019).

90. Huo, R. et al. APOE expression in papillary thyroid carcinoma: Influencing tumor progression and macrophage polarization. Immunobiology 229, 152821 (2024).

91. Tavazoie, M. F. et al. LXR/ApoE Activation Restricts Innate Immune Suppression in Cancer. Cell 172, 825–840.e18 (2018).

92. Hui, B. et al. Inhibition of APOE potentiates immune checkpoint therapy for cancer. Int J Biol Sci 18, 5230–5240 (2022).

93. Liu, C. et al. Pan-Cancer Single-Cell and Spatial-Resolved Profiling Reveals the Immunosuppressive Role of APOE+ Macrophages in Immune Checkpoint Inhibitor Therapy. Advanced Science 11, 2401061 (2024).

94. Procopciuc, L. M., Caracostea, G., Zaharie, G. & Stamatian, F. Newborn APOE genotype influences maternal lipid profile and the severity of high-risk pregnancy – preeclampsia: Interaction with maternal genotypes as a modulating risk factor in preeclampsia. Hypertension in Pregnancy 34, 271–283 (2015).

95. Chen, Y. et al. Subtype specific immune-metabolic reprogramming in preeclampsia revealed by multiomics and serum biomarkers. Hypertens Res 49, 641–657 (2026).

96. Crosley, E. J., Elliot, M. G., Christians, J. K. & Crespi, B. J. Placental invasion, preeclampsia risk and adaptive molecular evolution at the origin of the great apes: evidence from genome-wide analyses. Placenta 34, 127–132 (2013).

97. Khong, Y. & Brosens, I. Defective deep placentation. Best Pract Res Clin Obstet Gynaecol 25, 301–311 (2011).

98. Morlando, M. & Collins, S. Placenta Accreta Spectrum Disorders: Challenges, Risks, and Management Strategies. Int J Womens Health 12, 1033–1045 (2020).

99. Manus, M. B. Evolutionary mismatch. Evol Med Public Health 2018, 190–191 (2018).

100. Nesse, R. M. & Williams, G. C. Why We Get Sick: The New Science of Darwinian Medicine. (Vintage, 2012).

101. Stearns, S. C. & Medzhitov, R. Evolutionary Medicine. (Oxford University Press, 2024).

102. Thomas, J. R., Naidu, P., Appios, A. & McGovern, N. The Ontogeny and Function of Placental Macrophages. Front Immunol 12, 771054 (2021).

103. Freyer, L. et al. Erythro-myeloid progenitor origin of Hofbauer cells in the early mouse placenta. Development 149, dev200104 (2022).

104. Vento-Tormo, R. et al. Single-cell reconstruction of the early maternal-fetal interface in humans. Nature 563, 347–353 (2018).

105. Jiang, X. et al. Identifying a dynamic transcriptomic landscape of the cynomolgus macaque placenta during pregnancy at single-cell resolution. Dev Cell 58, 806–821.e7 (2023).

106. Yang, Y. et al. Transcriptomic Profiling of Human Placenta in Gestational Diabetes Mellitus at the Single-Cell Level. Front Endocrinol (Lausanne) 12, 679582 (2021).

107. Berger, H., Gagnon, R. & Sermer, M. Guideline No. 393-Diabetes in Pregnancy. J Obstet Gynaecol Can 41, 1814–1825.e1 (2019).

108. Zhu, L. et al. Single-Cell Sequencing of Peripheral Mononuclear Cells Reveals Distinct Immune Response Landscapes of COVID-19 and Influenza Patients. Immunity 53, 685–696.e3 (2020).

109. Dobin, A. et al. STAR: ultrafast universal RNA-seq aligner. Bioinformatics 29, 15–21 (2013).

110. Laughney, A. M. et al. Regenerative lineages and immune-mediated pruning in lung cancer metastasis. Nat Med 26, 259–269 (2020).

111. Lee, H.-O. et al. Lineage-dependent gene expression programs influence the immune landscape of colorectal cancer. Nat Genet 52, 594–603 (2020).

112. Zhang, P. et al. Dissecting the Single-Cell Transcriptome Network Underlying Gastric Premalignant Lesions and Early Gastric Cancer. Cell Rep 27, 1934–1947.e5 (2019).

113. Mao, X. et al. Single-cell and spatial transcriptome analyses revealed cell heterogeneity and immune environment alternations in metastatic axillary lymph nodes in breast cancer. Cancer Immunol Immunother 72, 679–695 (2023).

114. Davidson, G. et al. Mesenchymal-like Tumor Cells and Myofibroblastic Cancer-Associated Fibroblasts Are Associated with Progression and Immunotherapy Response of Clear Cell Renal Cell Carcinoma. Cancer Res 83, 2952–2969 (2023).

115. Chen, J., Liu, Z., Wu, Z., Li, W. & Tan, X. Identification of a chemoresistance-related prognostic gene signature by comprehensive analysis and experimental validation in pancreatic cancer. Front Oncol 13, 1132424 (2023).

116. Goad, J. et al. Single-cell sequencing reveals novel cellular heterogeneity in uterine leiomyomas. Hum Reprod 37, 2334–2349 (2022).

117. Hirz, T. et al. Dissecting the immune suppressive human prostate tumor microenvironment via integrated single-cell and spatial transcriptomic analyses. Nat Commun 14, 663 (2023).

118. Qian, J. et al. A pan-cancer blueprint of the heterogeneous tumor microenvironment revealed by single-cell profiling. Cell Res 30, 745–762 (2020).

119. Li, K. et al. Single-cell dissection of the multicellular ecosystem and molecular features underlying microvascular invasion in HCC. Hepatology 79, 1293–1309 (2024).

120. Guo, C. et al. Spatiotemporally deciphering the mysterious mechanism of persistent HPV-induced malignant transition and immune remodelling from HPV-infected normal cervix, precancer to cervical cancer: Integrating single-cell RNA-sequencing and spatial transcriptome. Clin Transl Med 13, e1219 (2023).

121. Ganier, C. et al. Multiscale spatial mapping of cell populations across anatomical sites in healthy human skin and basal cell carcinoma. Proc Natl Acad Sci U S A 121, e2313326120 (2024).

122. Tripathi, S. et al. Pediatric glioma immune profiling identifies TIM3 as a therapeutic target in BRAF fusion pilocytic astrocytoma. J Clin Invest 134, e177413 (2024).

123. Hao, Y. et al. Dictionary learning for integrative, multimodal and scalable single-cell analysis. Nat Biotechnol 42, 293–304 (2024).

124. McGinnis, C. S., Murrow, L. M. & Gartner, Z. J. DoubletFinder: Doublet Detection in Single-Cell RNA Sequencing Data Using Artificial Nearest Neighbors. Cell Syst 8, 329–337.e4 (2019).

125. Korsunsky, I. et al. Fast, sensitive and accurate integration of single-cell data with Harmony. Nat Methods 16, 1289–1296 (2019).

126. Han, X. et al. Mapping the Mouse Cell Atlas by Microwell-Seq. Cell 172, 1091–1107.e17 (2018).

127. Han, X. et al. Construction of a human cell landscape at single-cell level. Nature 581, 303–309 (2020).

128. Jiang, X. et al. A differentiation roadmap of murine placentation at single-cell resolution. Cell Discov 9, 30 (2023).

129. Emms, D. M. & Kelly, S. OrthoFinder: phylogenetic orthology inference for comparative genomics. Genome Biol 20, 238 (2019).

130. Brawand, D. et al. The evolution of gene expression levels in mammalian organs. Nature 478, 343–348 (2011).

131. Paradis, E. & Schliep, K. ape 5.0: an environment for modern phylogenetics and evolutionary analyses in R. Bioinformatics 35, 526–528 (2019).

132. Murat, F. et al. The molecular evolution of spermatogenesis across mammals. Nature 613, 308–316 (2023).

133. Lin, Z. et al. Evolutionary-scale prediction of atomic-level protein structure with a language model. Science 379, 1123–1130 (2023).

134. Tarashansky, A. J. et al. Mapping single-cell atlases throughout Metazoa unravels cell type evolution. Elife 10, e66747 (2021).

135. Renfree, M. B. Implantation and Placentation Vol 2, Reproduction in Mammals: Embryonic and Fetal Development; Austin, CR, Short, RV, Eds. (1982).

136. Zhou, Y. et al. Metascape provides a biologist-oriented resource for the analysis of systems-level datasets. Nature communications 10, 1523 (2019).

137. Crowell, H. L. et al. muscat detects subpopulation-specific state transitions from multi-sample multi-condition single-cell transcriptomics data. Nat Commun 11, 6077 (2020).

138. Szklarczyk, D. et al. The STRING database in 2023: protein-protein association networks and functional enrichment analyses for any sequenced genome of interest. Nucleic Acids Res 51, D638–D646 (2023).

139. Bader, G. D. & Hogue, C. W. V. An automated method for finding molecular complexes in large protein interaction networks. BMC Bioinformatics 4, 2 (2003).

140. Shannon, P. et al. Cytoscape: a software environment for integrated models of biomolecular interaction networks. Genome Res 13, 2498–2504 (2003).

141. Efremova, M., Vento-Tormo, M., Teichmann, S. A. & Vento-Tormo, R. CellPhoneDB: inferring cell-cell communication from combined expression of multi-subunit ligand-receptor complexes. Nat Protoc 15, 1484–1506 (2020).

142. Carver, A. J., Taylor, R. J. & Stevens, H. E. Mouse In Vivo Placental Targeted CRISPR Manipulation. J Vis Exp 10.3791/64760 (2023) doi:10.3791/64760.

143. Ferguson, C. M. et al. Silencing Apoe with divalent-siRNAs improves amyloid burden and activates immune response pathways in Alzheimer’s disease. Alzheimers Dement 20, 2632–2652 (2024).

144. Livak, K. J. & Schmittgen, T. D. Analysis of relative gene expression data using real-time quantitative PCR and the 2(-Delta Delta C(T)) Method. Methods 25, 402–408 (2001).

145. Schneider, C. A., Rasband, W. S. & Eliceiri, K. W. NIH Image to ImageJ: 25 years of image analysis. Nat Methods 9, 671–675 (2012).

146. Chen, S., Zhou, Y., Chen, Y. & Gu, J. fastp: an ultra-fast all-in-one FASTQ preprocessor. Bioinformatics 34, i884–i890 (2018).

147. Kim, D., Paggi, J. M., Park, C., Bennett, C. & Salzberg, S. L. Graph-based genome alignment and genotyping with HISAT2 and HISAT-genotype. Nat Biotechnol 37, 907–915 (2019).

148. Danecek, P. et al. Twelve years of SAMtools and BCFtools. Gigascience 10, giab008 (2021).

149. Liao, Y., Smyth, G. K. & Shi, W. featureCounts: an efficient general purpose program for assigning sequence reads to genomic features. Bioinformatics 30, 923–930 (2014).

150. Love, M. I., Huber, W. & Anders, S. Moderated estimation of fold change and dispersion for RNA-seq data with DESeq2. Genome Biol 15, 550 (2014).

